# DNA origami 2.0

**DOI:** 10.1101/2022.12.29.522100

**Authors:** Nayan P. Agarwal, Ashwin Gopinath

## Abstract

DNA origami is a technique that allows the creation of precise, modular, and programmable nanostructures using DNA. These nanostructures have found use in several fields like biophysics, molecular biology, nanoelectronics, and nanophotonic due to their programmable nature as well as ability to organize other nanomaterials with high accuracy. However, they are fragile and unstable when removed from their optimal aqueous conditions. In contrast, other commonly used bottom-up methods for creating inorganic nanoparticles do not have these issues, but it is difficult to control the shape or spatial organization of ligands on these nanoparticles. In this study, we present a simple, highly controlled method for templated growth of silica on top of DNA origami while preserving all the salient features of DNA origami. Using the polyplex micellization (PM) strategy, we create DNA nanostructures that can withstand salt-free, buffer-free, alcohol-water mixtures, enabling us to control the material growth conditions while maintaining the monodispersity and organization of nanoelements. We demonstrate the growth of silica shells of different thicknesses on brick and ring-shaped DNA origami structures using the standard Stöber process. We also demonstrate the thermostability of the silica-coated nanostructures as well as accessibility of surface sites programmed into the DNA origami after the silica growth in the final inorganic nanostructure.

## Introduction

One of the main goals of nanotechnology, as laid out by Feynman in his seminal lecture “There’s plenty of room at the bottom” in 1959, was achieving absolute compositional and organizational control over matter with atomic precision. Since then, several techniques have attempted to achieve this stated goal through either a top-down lithographic approach or a bottom-up self-assembly approach. Top-down approaches, owing to its modular nature, has fundamentally transformed our society by enabling integrated electronics, optics, MEMS, microfluidics, and more. However, fundamental limitations associated with patterning capabilities as well as dependence to silicon prevent this approach from faithfully achieving Feynman’s vision. Bottom-up approaches, especially ones that rely on biomolecular assembly, have demonstrated a remarkable ability to create nearly atomically precise structures. However, from a compositional point of view bottom-up techniques suffer from being highly incompatible with each other, i.e., these bottom-up nanostructures stability and viability is tightly tied to their media in which they are synthesized making them incompatible with each other. There are however bottom-up techniques, like DNA nanotechnology and DNA origami, that has the qualities to overcome these difficulties.

By viewing DNA as a programmable polymer rather than an information-carrying molecule, practitioners of this field exploit the sequence-dependent Watson-Crick-Franklin base pair interaction to create nanoparticles of arbitrary two and three-dimensional shapes. These nanoparticles range in size from a few 10s of nanometers to a few microns, and they can take a variety of shapes, including nanotubes, discs, rings, polyhedra, and crystal lattices.^1–8^ In addition to controlling the shape and size, the technique allows the resultant nanoparticles to be viewed as molecular breadboards on which other nanoparticles could be organized with positional accuracy approaching 0.5 A (Figure 1-A).^9^ This capability has been particularly helped by the nearly six decades of development in DNA chemistry^10^ that enabled a variety of biological molecules and non-biological nanomaterials (like Quantum dots, carbon nanotubes, etc.) to be decorated with DNA.^11–17^ It is also worth noting that the yield and purity of the DNA origami nanostructures are nearly 100%, and its synthesis cost is also relatively low at ∼$300/gram.^18^ DNA origami can also be organized using top-down lithographic nanofabrication to enable ∼5 nm positional control^19^ and ∼3.5 degrees orientational control of single molecules on planar substrate with.^20^ In fact, it would be accurate to state that no other nanofabrication technique simultaneously offers such a diverse range of capabilities However, these DNA origami nanoparticles are structurally fragile, the particles are incompatible with organic solvents, have limited thermostability, and the material properties are not particularly relevant for any novel property or relevant applications. This limits the usage of DNA nanoparticles since their establishment, as technologically relevant materials, processes, or chemical reactions are typically most stable in non-polar organic solvents and are incompatible with the presence of cations.^21–26^

**Figure 1.**
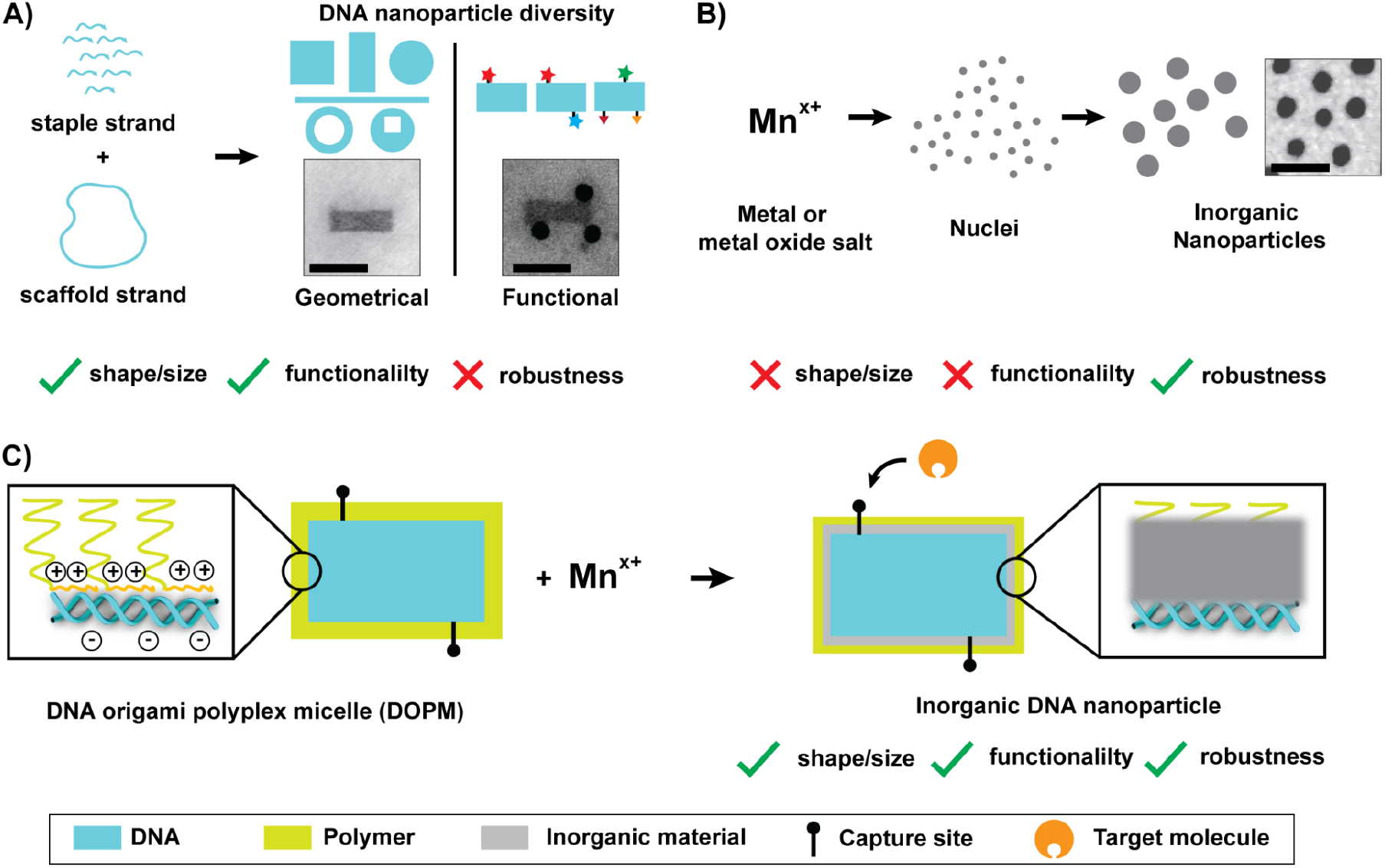
Illustration of the strategy to manufacture robust nanoparticles with programmable shape, size, and surface functionality. A) The self-assembly of a DNA nanoparticle utilizes long single-stranded DNA (ssDNA) called “scaffold strand”, that is held together by distinct short ssDNA oligonucleotides called “staple strands”, to create diverse two- and three-dimensional nanoparticles with ease and high-purity. The surface of DNA nanoparticles can be used to place any number of molecules (inorganic or biological) at pre-determined locations with unprecedented precisions. However, these nanoparticles lack robustness. B) In contrast, inorganic nanoparticles are typically synthesized by reducing metal or metal oxide salts to obtain nuclei, which condense to form nanoparticles. These nanoparticles are robust; however, the manufacturing process faces certain fundamental, chemical, and physical limitations to obtain programmability of shape, size, or surface functionality. C) As a solution, we propose combining the two approaches. To achieve this, we utilize the salt- and solution-independent DNA origami polyplex micelles (DOPMs), which can grow inorganic materials in solutions that favor the material growth. These DNA nanoparticles can be designed to carry a specific number of functional ligands placed at distinct locations, which will be maintained during the material growth. This strategy can be used to produce functional inorganic DNA nanoparticles.

A completely different approach to achieving Feynman’s vision involves synthesizing inorganic nanoparticles from the bottom-up using techniques like the Stöber process (also known as the sol-gel method),^27^ hydrothermal,^28^ co-precipitation,^29^ or microemulsion.^30^ Inorganic nanoparticles (metal or metal-oxide nanomaterials) produced by such techniques possess material properties unachievable by any other top-down approaches. These particles are stable over various environmental conditions, such as temperature, solution, pH, and solvents. While these techniques can be adapted to change the material composition and size of the nanoparticles with relative ease, however, it is extraordinarily difficult to change the surface ligand composition, shape of the particle, to break the symmetry of the particle, or to tightly control the size of the particles while maintaining high monodispersity (Figure 1-B). Such difficulties arise from the complex dependency of such chemical processes on various reaction parameters, each of which can influence the nanoparticle’s stability, reactivity, and other physical properties. Achieving the desired material property requires a deep understanding of the underlying chemistry and physics of the synthesis process, as well as careful experimentation and optimization of the synthesis conditions.

While both these dramatically different approaches for nanoparticle synthesis have independently matured over the last several decades, neither has achieved what can be regarded as modular, compositional, and organizational control over matter at the atomic or molecular scale. However, by combining the best aspects of both these techniques, it should be possible to create a generalized technique and framework for manufacturing nanoparticles that offer the immense programmability of DNA origami and the material purity and compositional control of the inorganic nanoparticle synthesis. A few attempts to achieve this goal have also been made, during the last decade, by using DNA origami as templates or molds to grow inorganic materials such as gold (Au), platinum (Pt), palladium (Pd), silver (Ag), silica (SiO_2_), etc.^31–38^ However, these attempts have not been successful since the key benefits of one or both techniques were compromised. The primary reason for this lack of success is the incompatibility between the experimental conditions best suited for inorganic material growth and those for DNA origami stability. Specifically, the quality of the material during inorganic nanoparticle synthesis is completely dependent on the solution conditions (such as precursor concentration, temperature, presence of impurities, etc.), and these reactions are typically done in organic solvents. In contrast, DNA origami is very robust to synthesis conditions; however, its stability typically requires it to be in an aqueous solution with a set concentration of cations and the solution temperature not to exceed ∼50-100 °C.^39–41^ This fundamental incompatibility forced efforts for templated material growth on DNA origami to be conducted in a media that is most suited for the stability of the DNA nanoparticle. This decision leads to inferior results with little control over the material purity, the structural dimensions (like thickness), and the loss of DNA origami’s ability to be treated as a molecular breadboard.

In this paper, we introduce a new, facile approach to templated growth of silica on DNA origami nanoparticles while preserving all the salient features of the DNA origami, i.e., tightly controlled shape, which is already accepted to be programmable, monodispersity, and most importantly, the ability to organize nanoelements on the final inorganic nanoparticle arbitrarily. We believe the resultant nanoparticle is a fundamental step towards a new class of programmable nanoparticles and thus refer to it as DNA origami 2.0. Our choice to grow silica on the DNA origami was driven by the wealth of literature detailing silica growth reaction and its compatibility with other inorganic material growth.^42,43^ We view these two properties of silica as making it a proxy for other inorganic material, i.e., if silica can be templated on DNA origami, any other inorganic material could also be templated analogously on DNA origami or top of the silica shell already grown on the origami. Further, silica is highly biocompatible, inert, and optically transparent, making it amenable for downstream practical applications.

Typically, silica nanoparticles are synthesized using the Stöber process (also known as the sol-gel process), which is conducted by introducing a molecular precursor (an alkoxide) into an alcohol-water mixture. The growth is driven by the interplay between the hydrolysis of the precursor into silanols as well as the condensation of silanols into silica nanoclusters that, in turn, grow into nanoparticles. Conceptually, we can represent this using the following three equations.^44^

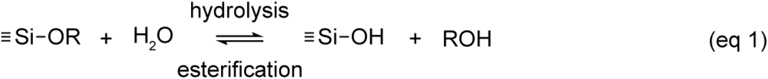

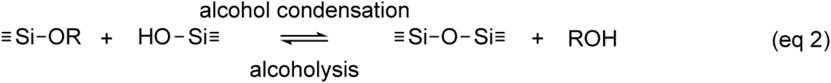

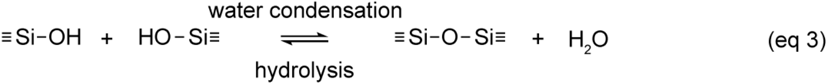

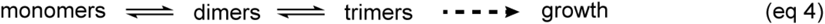

In practice, these reactions occur concurrently, and the detailed kinetics of nanoparticle formation is dependent on several independent parameters, primary among them are the choice of precursor and precursor concentration, choice of solvent, pH conditions, alcohol-to-water ratio as well as the water-to-silica ratio. The control of nanoparticle formation was typically achieved by tuning these parameters such that the three steps of the growth process were rendered independent, and one could move the system from one condition to another. All previous attempts to grow silica on top of DNA origami involved working in an aqueous solution, which is not conducive to enabling such precise control.^33–35^ Further, these previous works also had buffering agents (Tris, MOPS, etc.) and cations (Na^+^, Mg^2+^, etc.) which serve as impurities, having a varied effect on nucleation and growth of nanoparticles while also leading to spontaneous uncontrolled nucleation.^45^ Our approach to creating DNA origami 2.0 involved using the polyplex micellization (PM) strategy^46,47^ to make DNA origami stable in a salt-free, buffer-free, alcohol-water mixture, a condition in which silica growth has been demonstrated to be tightly controlled. The PM is achieved using poly(ethylene glycol)-b-poly(L-lysine) block copolymers (PEG-PLL), where the cationic PLL section strongly binds electrostatically to the DNA structures. At the same time, the PEG chains form an uncharged polymeric shell around the DNA origami that enables it to be structurally stable in arbitrary solvents.^46–48^

To demonstrate and optimize the synthesis of DNA origami 2.0 particles, we use DNA origami in the shape of a brick (42-helix bundle or 42HB).^49^ The polyplex micellization of the DNA brick was achieved using poly(ethylene glycol)-*b*-poly(L-lysine) block copolymers (PEG-PLL) with either 1, 5, or 20 kDa long PEG units, but with 10 lysine units in all cases (Figure 1-C).^46,47^ We characterized the results using dynamic light scattering (DLS). This bulk measurement technique enables observing changes in the hydrodynamic diameter (Z_avg_, measured in nanometers) and the dispersity of particles for the entire duration of the growth as a function of time. A brief description of the measurement process and interpretation of the observed values are detailed in supporting note 1.

The DLS-based size analysis of 42HBs in the native buffers (10 mM Tris, 1 mM EDTA, and 5 mM MgCl_2_ maintained at pH 8.3) resulted in a single distribution peak, indicating a monodispersed solution with the Z_avg_ of 52.15 ± 0.73 nm (Figure S 1-A, tube 1). Here, despite the 42HBs having a cuboidal shape of dimensions 55×20×10 nm, the DLS algorithm simplifies it to an equivalent sphere of diameter Z_avg_ because of the Stokes-Einstein equation (supporting note 1). Next, we assessed the effect of different buffer conditions on the 42HB DNA origami polyplex micelles (DOPM) prepared using 5 kDa long PEG units (Figure S 1-A, tubes 2 to 5). From the DLS results, we observed that in the native buffers, 42HB DOPMs were monodispersed with a Z_avg_ of 61.48 ± 0.42 nm (Figure S 1-A, tube 2), which is 17.89% (9.33 ± 0.84 nm) more than the non-polyplexed 42HBs. This indicated that the thickness of the 5 kDa PEG shell around the 42HBs in the DOPMs is 4.67 ± 0.42 nm, consistent with the previous studies.^46–48^ Transferring the 42HB DOPMs into ultrapure water by buffer exchange led to a further increase in the size of the particles to Z_avg_ of 63.73 ± 0.51 nm (Figure S 1-A, tube 3), while still exhibiting a single distribution peak in DLS. This suggests that the particles are intact and stable in a salt-free solution. The additional 4.3% increase (2.25 ± 0.66 nm) in hydrodynamic radius can be explained by the core of the DNA origami slightly expanding due to the removal of the divalent cations known to condense and give structural rigidity to DNA origami.^50–52^ Finally, the 42HB DOPMs were transferred into an 85% ethanol solution. The DLS analysis suggested that the particles were intact and non-aggregating, but the effective size of the particle increased to a Z_avg_ of 89.32 ± 1.38 nm (Figure S 1-A, tube 4). We hypothesize that this 25.59 ± 1.47 nm increase in the thickness is due to the PEG chains acquiring an extended conformation due to its interaction with ethanol rather than the compressed mushroom conformation observed in aqueous conditions.^48,53^ We added concentrated ammonium hydroxide to the 85% ethanol solution with 42HB DOPM to increase the effective pH of the solution to 11 and observed a 9.8% decrease in the Z_avg_ to 80.55 ± 1.03 nm (Figure S 1-A, tube 5) while no change in the shape of the distribution. Such behavior indicates that the PEG chain length shortens in the presence of hydroxide ions, disrupting the interactions between the PEG chains and ethanol molecules. Hydroxide ions are highly reactive and can potentially react with both the PEG chains and the alcohol molecules, breaking the bonds between them and reducing the overall strength of the interaction. These experiments demonstrate the structural viability and monodispersity of DOPMs in salt-free, partially non-polar, alcohol-water solutions.^48^ The ethanol-based solvent system is also well-accepted for the Stöber process.

The SiO_2_ growth process involved adding tetraethyl orthosilicate (TEOS) as a precursor to the 85% ethanol solution at pH 11 with the 42HB DOPMs, followed by incubation at room temperature for a fixed time interval. We characterized the growth using DLS (Figure S 1-B). For the first 15 mins of the reaction, we observed no change in the hydrodynamic diameter of the particles in the solution (Z_avg_ fluctuating between 77.29 nm and 81.46 nm, consistent with 42HB DOPMs), followed by a significant increase of Z_avg_ around 188 nm at the 16 min time point (Figure S 1-B, blue curve). After this time, we also observed additional size distribution peaks, indicating polydispersity and particle aggregation. The scattering intensity measured by the detector is directly proportional to the inorganic material growth that supposedly takes place on the surface of the DNAO. By analyzing the scattering intensity data as a function of time (Figure S 1-B, orange curve), we observed a 34% increase in scattering intensity during the first 15 min of incubation, which confirmed that prior to aggregation, the silica growth takes place in the solution, on the surface of the DNAO. These observations suggest that small nucleation centers are constantly being generated in solution. At the same time, the silica shell grows on the DNA underneath the PEG layer on the 42HB DOPMs, which restricts any observable change to the Z_avg_. The PEG shell also prevents the formation of aggregates. However, aggregates appear after 15 minutes when the silica shell on the DNAO is thicker than the PEG shell. We repeated the growth without DOPMs templates as negative control while maintaining all the other solution conditions (Figure S 2). Here, we did not observe any condensed particles during the 60 min incubation, which suggests that it is necessary to have a template in the solution that directs the condensation of the nuclei. From this, we can conclude that in the absence of any impurity (multivalent cations, buffering agents, and other non-participating molecules), the formation of side products is restricted, potentially limiting the availability of nuclei and the subsequent growth of the inorganic material to the DOPMs only.

These results indicated that templated growth of silica is possible on DOPMs in an alcohol-water solution. However, in order to have tighter control over the silica growth process as well as to minimize spurious nucleation of silica nanoclusters, we investigated the following parameters: the pH of the reaction solution (pH 7, pH 8.3 or pH 11), the choice of solvent (methanol or ethanol), alcohol content (0%, 10%, 50%, 85% or 95%), the choice of the precursor (TMOS or TEOS), the concentration of the precursor with respect to the DOPMs, the length of PEG units in PEG-PLL (1 kDa, 5 kDa or 20 kDa), as well as pH transitions and the addition of PEG-silanes for reaction termination.

We assessed the effect of the different pH conditions (pH 7, pH 8.5, and pH 11) in 85% ethanol solution on 42HB DOPMs with 5 kDa PEG chain lengths (Figure 2-A). We observed from the DLS measurements that the Z_avg_ of the DOPMs at pH 7 and 8.5 are comparable at 87.56 ± 1.2 nm and 88 ± 2 nm, respectively (Figure 2-B). While for pH 11, the Z_avg_ reduces to 80 ± 0.64 nm (Figure 2-B). We attributed the decrease in Z_avg_ to the presence of excess OH^-^ ion in the solution, which led to a change in PEG confirmation. Next, we initiated the growth reaction at these pH conditions by adding 1x concentration of TEOS (equivalent to 37.2 mM of TEOS per 1 nM of DOPMs, precursor concentrations explained in the following section). From the DLS results, we conclude that no silica growth had taken place at pH 7 or 8.5 over a period of 60 min (Figure 2-C and D, overlapping grey and orange curves). We believe this is due to the slow rate of hydrolysis at these conditions that produce little to no nuclei. However, at pH 11, catalyzed by OH^-^ ions, the hydrolysis of TEOS proceeds at a faster rate that reaches completion, immediately followed by condensation, which takes place at a steady rate due to a constant supply of monomers. From this, we can conclude that pH 11 is ideal for silica shell growth.

**Figure 2.**
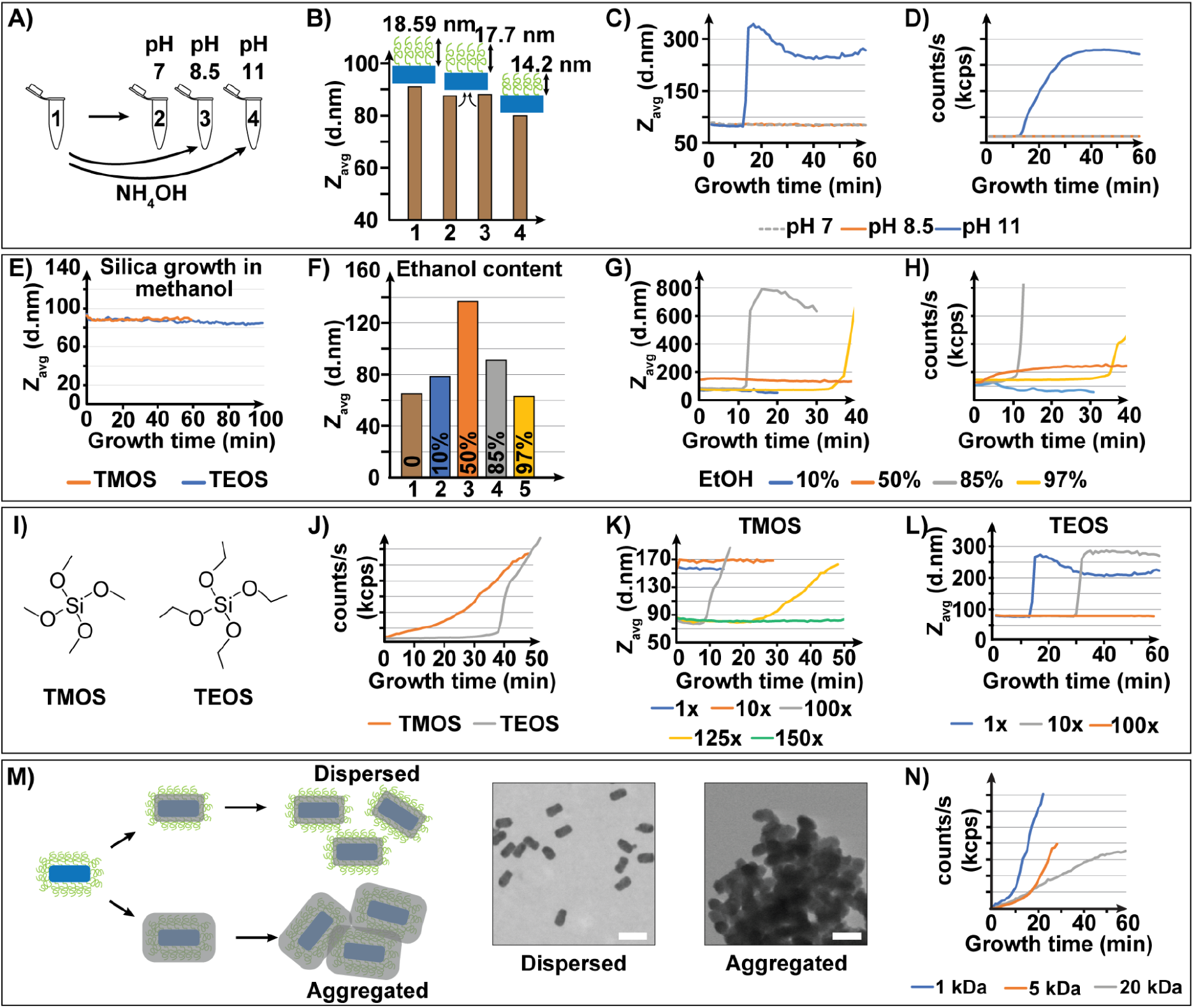
DNA brick origami structures are subjected to various optimized conditions that impact the silica shell growth to obtain fine-tuned control over the growth process. A-D) DLS data for the optimization tests for solvent pH conditions. E-H) DLS data for alcohol-to-water content optimizations. I-L) DLS data for optimizing the choice of precursor (molecular structure of the tested precursors in I) and precursor concentrations. M-N) DLS and TEM data for PEG-length optimization studies. Scale bars = 100 nm.

To further investigate the reliability of each step (hydrolysis, condensation, and growth) on the solvent pH condition, we added hydrochloric acid (HCl) after 10 min of reaction to change the pH of the solution from pH 11 to pH 7. This pH transition during the growth reaction led to an uncontrolled condensation that was observed as aggregate formation immediately (Figure S 3). In contrast to the reaction at pH 7, where we did not observe any growth (Figure 2-C and D), this uncontrolled condensation is due to the pH transition, which confirms our earlier belief that at pH 11, the reaction undergoes complete hydrolysis and moving to pH 7 led to an increase in the rate of condensation resulting in the formation of gel-like products. We concluded that such pH transitions could be used to terminate the reaction.

Next, we investigated the dependency of the silica shell growth process on the solvent type. Typically, the use of a precursor-miscible solvent facilitates the rate of hydrolysis. For this, we compared methanol to ethanol as a solvent for silica growth while keeping all the other parameters constant at 85% alcohol composition, pH 11, and using 42HB DOPMs with 5 kDa PEG length. We initiated the growth reaction by adding tetramethyl orthosilicate (TMOS) or TEOS as silica precursors. From the DLS measurements for the samples with methanol as the solvent, we did not observe any growth over 60 min and 100 min for either TMOS or TEOS as a precursor, respectively (Figure S 4). We hypothesize that this behavior could be a combination of several reasons. Firstly, methanol forms a stronger hydrogen bond with water molecules than the one between them. As a result, at 85% methanol content, the diffusion of water is restricted, which reduces the rate of hydrolysis of the precursor. Secondly, methanol stabilizes the silanols (-Si-OH) produced after hydrolysis, reducing the condensation rate. Thirdly, methanol hydrogen bonds to silicates (-SiO^-^), which reduces their nucleophilicity, further affecting the condensation of monomers. Lastly, methanol reduces the redistribution of the growing silica shell by breaking siloxane bonds *via* alcoholysis or hydrolysis.^44^ Due to these reasons, we did not further investigate methanol as a solvent and used ethanol for all the subsequent studies.

### Alcohol/water content

Typically, undiluted silanes (precursors) remain unhydrolyzed upon exposure to water due to the liquid/liquid phase separation.^45^ To get around this issue, homogenization agents like alcohols, tetrahydrofuran (THF), acetone, etc., are added to prevent phase separations as well as supporting the silica growth reaction. However, the growth can’t occur in pure organic solvent because some amount of water is necessary to promote hydrolysis. A delicate balance is thus required between the hydrolysis and alcoholysis (equations 1 to 4) for controlled growth and it is crucial to understand the precise influence of alcohol/water content in the reaction solution.

We assessed the following ethanol compositions: 10%, 50%, 85%, and 97%, using 42HB DOPMs with 5 kDa PEG lengths. To begin, we measured the Z_avg_ of DOPMs in these ethanol compositions. We found that the increase in the alcohol composition increased the Z_avg_ from 65 ± 1.83 nm for pure water condition to 78.32 ± 2.63 nm for 10% ethanol, with the maximum Z_avg_ of 136 ± 5 nm at 50% ethanol, followed by a decrease to 90.97 ± 2.7 nm and 62.88 ± 0.9 nm for 85% and 95% ethanol compositions, respectively (Figure 2-E). We hypothesize that such a trend results from the hydrogen bonding between the glycol units in PEG chains with ethanol and water. The chains are maximally stretched to 42 nm for 50%, depending on the ethanol composition. Following a decrease in the water content leads to the relaxation of the PEG chains to 19.41 nm and 5.36 nm at 85% and 97% ethanol composition, respectively. Interestingly, the PEG chains in both pure water and 97% ethanol composition exhibit comparable lengths of 3.52 nm and 5.36 nm, respectively (Figure 2-F, 1, and 5). We conclude that, independently, both water and ethanol have a similar impact on the PEG chain lengths. For specific applications, 50% ethanol composition can promote spatial separation of terminal functional groups on PEG chains further away from the structure, making them more accessible for target molecules.

Next, we initiated the growth reaction at these ethanol compositions at pH 11 by adding 1x concentration of TEOS. By observing the Z_avg_ and the scattering intensity for 40 min, we concluded that no growth took place for the entire duration at 10% and 50% ethanol content (Figure 2-G and H, no changes were observed in either, even after 180 min of incubation, data not shown). While for 97% ethanol composition, the growth was observed to be nearly 3 times slower compared to 85%, as the aggregation time point for 97% ethanol appears around the 35 min time point compared to 12 min for 85% composition. We hypothesize that this is due to the slower rate of hydrolysis at a low water content setting (3%), which led to the lack of monomers required for the growth. We concluded that the 85% ethanol composition was ideal for silica growth.

### Choice of precursor (TMOS v/s TEOS)

Hydrolysis of the precursor is the first step toward the silica shell’s growth. The kinetics of this reaction is influenced by the steric bulk and the electron-providing/withdrawing characteristics of the organic substituents attached to the silicon atom. Thus, the real growth will be intimately dependent on the choice of precursor. We compared the choice between TMOS or TEOS (Figure 2-G) as precursors in our reaction with the hypothesis that the methoxy (-O-CH_3_) substituent in TMOS would undergo hydrolysis at a faster rate compared to ethoxy (-O-C_2_H_5_) in TEOS, as its more electronegative and a better leaving group.

We used 42HB DOPMs with 5 kDa PEG lengths to grow silica shells using either precursor, keeping all the other parameters constant at 85% ethanol and pH 11. We performed the growth reaction for 50 min and plotted the scattering intensity as a function of time (Figure 2-H). The DLS results supported our hypothesis that the reaction proceeds 4-5 times faster for TMOS than TEOS. The plot suggested that the silica growth using TMOS follows a linearly (Figure 2-H, orange curve), while for TEOS, the growth is switch-like where large aggregates starts forming ∼38 mins (Figure 2-H, grey curve). This behavior is due to the slower rate of hydrolysis for TEOS compared to TMOS, which delays the condensation of silica. We further believe that for TEOS, the condensation of monomers only takes place after the complete displacement of all four substituents. However, only the displacement of the first substituent is the rate-limiting step, and the following displacement reactions take place simultaneously. In contrast, for TMOS, due to the faster rate of hydrolysis, there is a steady supply of nuclei throughout the process, giving it a consistent, linear model for shell growth. From this, we concluded that either of the precursors is suitable for silica growth. However, TMOS provides a better opportunity for facile and controllable growth.

To further assess the two precursors, we investigated the effect of their concentrations on silica growth. For this, we observed the aggregation time points for varying concentrations of TMOS or TEOS while keeping all the other parameters constant at 85% ethanol, pH 11, and 5 kDa PEG. The precursor concentration was varied between 1x and 150x, where x represents the dilution factor, with 1x representing the highest concentration corresponding to 9,300 molecules per nm^2^ of the origami or 37.2 mM of precursor per 1 nM of origami. We conducted the silica growth for 50 min and plotted the Z_avg_ as a function of time. We observed that 1x and 10x concentrations of TMOS (Figure 2-I) led to aggregation immediately after precursor addition. In comparison, 1x and 10x concentrations of TEOS exhibit slower growth, and the structures aggregated after 16 and 31 minutes, respectively (Figure 2-J). This result is consistent with our previous observation.

Finally, we investigated the impact of the PEG chain on the growth process. We used three variations of PEG-PLL with a constant cationic segment formed with 10 lysine repeats while changing the PEG chain lengths to 1 kDa, 5 kDa, or 20 kDa. We hypothesize that, like a surfactant, the external PEG shell prevents aggregation during the silica growth process, keeping the structures monodispersed and allowing a controlled deposition of silica (Figure 2-M). Initially, we measured the Z_avg_ of 42HB DOPMs with varying PEG chain lengths in different buffer conditions (Figure S 5). As per expectations, in the native buffers, the Z_avg_ for 42HB DOPMs with 20 kDa PEG increased to 83 ± 1.4 nm from 52.15 ± 0.73 nm for 42HB DNAO and 61.48 ± 0.42 nm for 42HB DOPMs with 5 kDa PEG. Like the 5 kDa PEG in previous sections, the Z_avg_ in 85% ethanol for DOPMs with 20 kDa PEG increased by 46 ± 0.9 nm to 128.8 ± 2.5 nm. This increase is proportional to the increase in the length of the PEG chain, with an average chain length of 20 kDa in 85% ethanol being 38.33 ± 1.3 nm and 18.59 ± 0.78 nm for 5 kDa. However, the transition to pH 11 solution decreased slightly to 127.5 ± 1.5 nm. During this study, we observed aggregation for 1 kDa PEG even in native buffers (Figure S 5). We hypothesize that smaller PEG lengths (≤1 kDa) are insufficient to avoid the aggregation and the structures form aggregates during the polyplex micellization process. This observation further confirmed the reliance on longer PEG lengths for the stability and monodispersity of the structures.^47^

We used the 42HB DOPMs with varying PEG chain lengths for silica growth while keeping all the other parameters constant at 85% ethanol, pH 11, and using 125x concentration of TMOS as a precursor. We observed the silica shell growth for 60 min. We plotted the scattering intensity data as a function of time (Figure 2-L, corresponding Z_avg_ data in Figure S 6). We observed that for 1 kDa PEG the growth occurs steadily during the initial 10 min of incubation, followed by rapid growth causing aggregation in addition to the aggregates formed in the polyplex micellization step (Figure 2-L, blue curve). It is crucial to note that for 1 kDa PEG length, even prior to the silica growth, the 42HB DOPMs exhibited polydispersity and a cumulative Z_avg_ of 336.5 ± 11 nm. From these results, we concluded that the 1 kDa PEG lengths were not suited for our DNA origami 2.0. In the case of 5 kDa PEG, we observed steady growth until the aggregation timepoint of ∼16 min. While for 20 kDa PEG the growth takes place steadily for the entire incubation, which suggests that longer PEG lengths maintain a monodispersed solution enabling uniform shell growth (Figure 2-L, orange curve for 5 kDa and grey curve for 20 kDa. Corresponding Z_avg_ data in Figure S 6). We believe that to maintain a similar steady growth condition for 5 kDa PEG, it would be necessary to use lower precursor concentrations, albeit this would result in thinner silica shells as the rate of silica shell growth is directly proportional to the concentration of the precursor.

We extended this further and evaluated the silica shell growth at varying TMOS concentrations for 20 kDa PEG lengths while keeping all the other parameters constant (Figure 3-A and Figure S 7). We concluded that longer PEG chain lengths ensure a steady rate of silica deposition despite the increased rate of silica condensation at higher concentrations. By maintaining a constant growth time of 3 hours, we obtained different shell thicknesses at different TMOS concentrations. Specifically, we observed 6 nm growth for 125x TMOS, 5.2 nm for 150x TMOS, and 3.63 nm for 200x TMOS concentrations. (Figure 3-C; supporting note 2, Figure S 23 and Figure S 24 for more details regarding size measurements). From these results, we concluded that the silica shell thickness could be tuned by performing the silica shell growth at lower precursor concentrations, followed by longer incubation times, and undergoing complete exhaustion of the precursor to stop the reaction. However, such a reliance on the exhaustion of the precursors for reaction termination increases reaction times (3-24 hours) and is accompanied by side products (non-templated silica particles, aggregates) which reduced the reproducibility of the technique as well as introduces the need for downstream purification.

**Figure 3.**
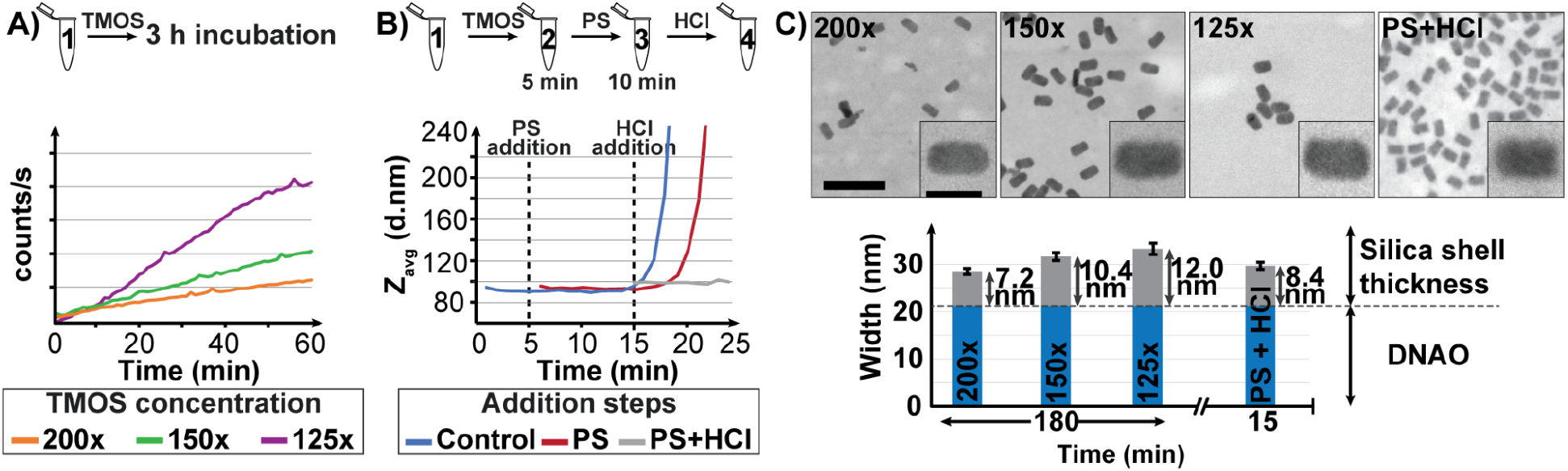
A-D) DLS and TEM data for growth reactions using varying TMOS concentrations were conducted for 3 hours on 42HB DOPMs with 20 kDa PEG chain lengths. For D and G, Scale bars = 200 nm for the top row and 50 nm for the inset. The blue part of the histograms depicts the width contributed by the DNAO core, while the grey part of the bars represents the total silica shell width obtained after the growth reaction. E-H) DLS and TEM data for optimizing conditions necessary to stop the reaction using PEG-silane (molecular structure in A) and pH transition using HCl.

Alternative approaches to controlling the thickness of silica layer growth would involve pH transitions using acids like HCl to terminate the reaction or use of capping agents like PEG-silanes (PS for short, 2-[methoxy (polyethyleneoxy) propyl] trimethoxy silane with 6–9 (CH2CH2O) units).^54–56^ To investigate these alternatives, we studied the impact of the addition of PS, HCl, or a combination of both on the growth reaction.

The capping agent’s introduction or the pH change was investigated independently. We conducted the silica shell growth on 42HB DOPMs with 5 kDa PEG lengths, using a 10x concentration of TEOS as the precursor. At first, the silica shell growth was terminated by adding varying concentrations of PS after performing the growth for 10 min (Figure S 8-A). The PS concentration was relative to the precursor concentration, described by the PS/precursor molar ratio. For stoichiometry optimization, the ratio was varied between 10:1 and 0.01:1. From the observed Z_avg_ and scattering intensity plots we concluded that the addition of PS does indeed stop the growth reaction for all the variants (Figure S 8-B and C). Further, the Z_avg_ remained constant over the next 24 hours (and 75 hours for the variant with 1:1 PS concentration), confirming our hypothesis that PS does indeed terminate the growth reaction.

Next, the silica shell growth was terminated by adding HCl (in place of PS) to the above reaction using TEOS. Adding HCl changed the solution pH from 11 to 7 (Figure S 9-A). We hypothesized that the transition to pH 7 would minimize the hydrolysis rate, reducing the monomers’ supply. In contrast, the higher rate of condensation will lead to rapid growth followed by reaction termination.^44^ We performed the growth for varied times (10, 20, 25, and 30 min), followed by adding HCl. From the Z_avg_ and scattering plots, we concluded that the pH transition did indeed lead to the termination of the growth reaction (Figure S 9-B and C). From the results, we concluded that the Z_avg_ of the resulting structures was directly dependent on the time points at which HCl was added.

Finally, we investigated the optimized PS concentration (0.1:1) to stop the growth reaction using TMOS (in place of TEOS) as a silica precursor. In contrast to the above observations, adding PS did not terminate the silica shell growth. Instead of stopping the silica growth completely, the capping agent only delayed the aggregation by 4 min (Figure 3-B orange curve). We hypothesized that this inability of PS to terminate the growth using TMOS as a precursor is due to the higher rate of hydrolysis of TMOS compared to TEOS, which out-competes the addition of a capping agent to the growing shell. We made similar observations for reaction termination using HCl, where despite termination, the shell growth was observed to be 5 times faster as compared to TEOS (Figure S 9 to Figure S 12). To circumvent this, we employed a combination of adding PS and HCl. We added PS after 5 min of growth reaction and HCl after 10 min (Figure 3-B). The DLS results confirmed the termination of the growth reaction (Figure 3-B gray curve), and we obtained a shell thickness of 4.23 nm with a total reaction time of 15 min (Figure 3-C).

It is necessary to note that the time points for adding PEG-silane and HCl need to be optimized using the DLS data. These additional time points can vary from batch to batch (Figure S 13) and across different structures. However, such optimizations are unnecessary for growth reactions performed using lower precursor concentrations that do not require the addition of PS and/or HCl.

### Optimized growth conditions

With the optimized growth conditions in hand, we studied the modularity of the technique by applying it to a dramatically different DNA origami shape, a ring (Figure 4-F).^57^ The growth was performed on DNA rings DOPMs prepared using 20 kDa PEG chain length in 85% ethanol solution at pH 11, using 125x TMOS concentration. We incubated the reaction for 3 hours to allow the reagents to exhaust, terminating the growth process. From the electron micrographs, we observed that the structures across all the conditions were monodispersed and do not exhibit any form of aggregation at any step of the process (Figure 4 B-D for DNA bricks and G-I for DNA rings, wide-field images in Figure S 14 to Figure S 22). We observed that for the DNA brick-silica nanoparticles, the width increased by 40%, from 21.27 ± 1.1 nm for DNAO to 30.61 ± 2.12 nm for silica structures (Figure 4 E), while the length decreased by 8%, from 57.15 ± 0.83 nm for DNAO to 54.12 ± 1.85 nm for silica structures. Similarly, the pore diameter of DNA ring structures decreased by 59%, from 31.2 ± 2.415 nm to 12.85 ± 1.69 nm (Figure 4 J), while the outer diameter decreased by 3.6%, from 51.86 nm for DNAO to 49.97 nm for silica structures. These results show consistent material growth of 4.12 nm thickness for DNA brick and 4.67 nm for DNA ring. However, the structure seems to have undergone some conformation change during the silica templating process. While it is beyond the scope of this work to prove the precise reasons for this conformational change, we believe it is due to the solvent conditions and the non-uniform growth of silica on different parts of the origami. The control DNAO and DOPMS were stained with uranyl formate to allow better contrast in the resulting images. At the same time, we did not perform such a DNA staining process for the silica-coated structure. The difference in contrast and edge clarity of the structures provided a clear indication of material growth. We also imaged the DNAO structures and the structures with the silica shells on the same TEM grid, and we observed a clear difference in material-related contrast due to the silica shell (Figure 5 A and D and supporting Figure S 25 and Figure S 26).

**Figure 4.**
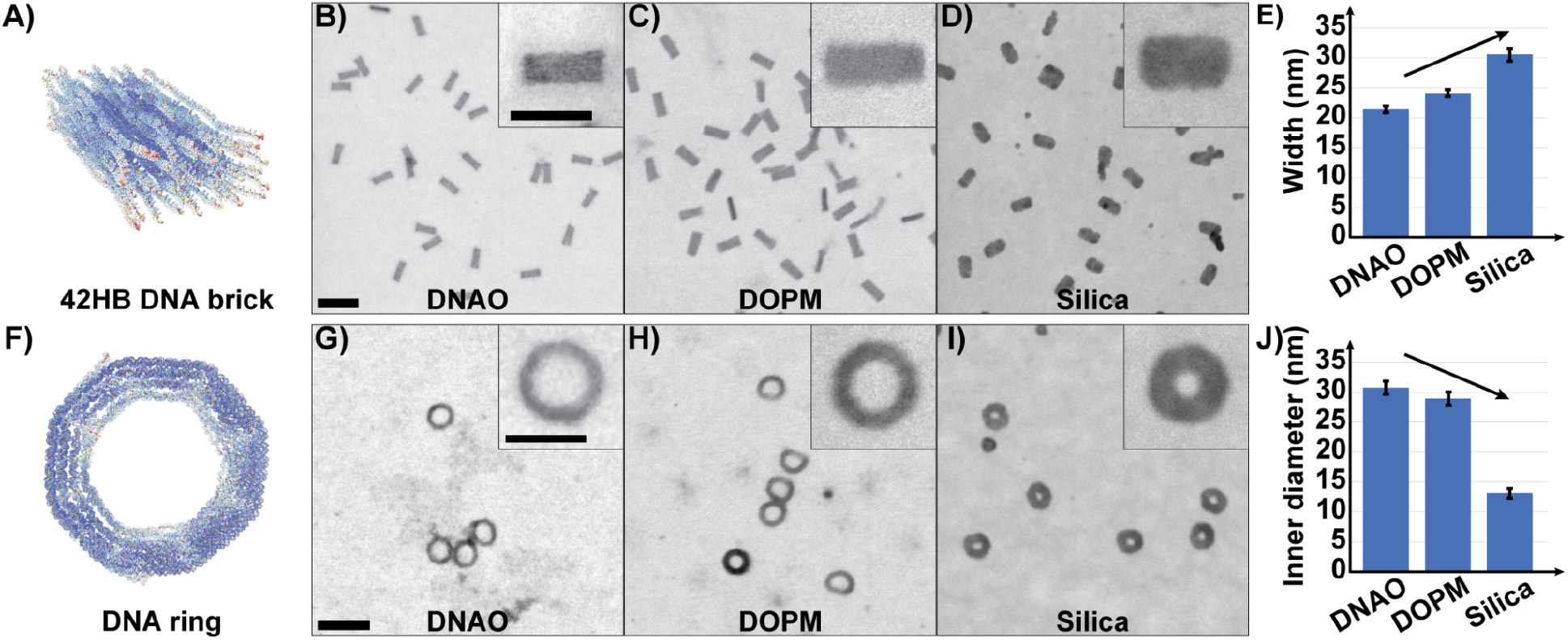
Test structures for inorganic material growth using DNA origami as a template. A-E) 42HB DNA origami (DNA brick), F-J) Delftagon DNA origami (DNA ring). A and F) Representative oxDNA models for the corresponding DNAO structures. TEM micrographs B and G) DNAO, C and H) DOPM, and D and I) Silica-coated structures. For all the TEM images, Scale bars = 100 nm and 50 nm for the inset. E and J) Graphs comparing the width (for DNA brick) and the inner diameter (for DNA ring) of structures across all the variations as labeled on the x-axis (Please refer to Figure S 23, Figure S 24 and supporting note 2 for more details regarding the measurement process).

**Figure 5.**
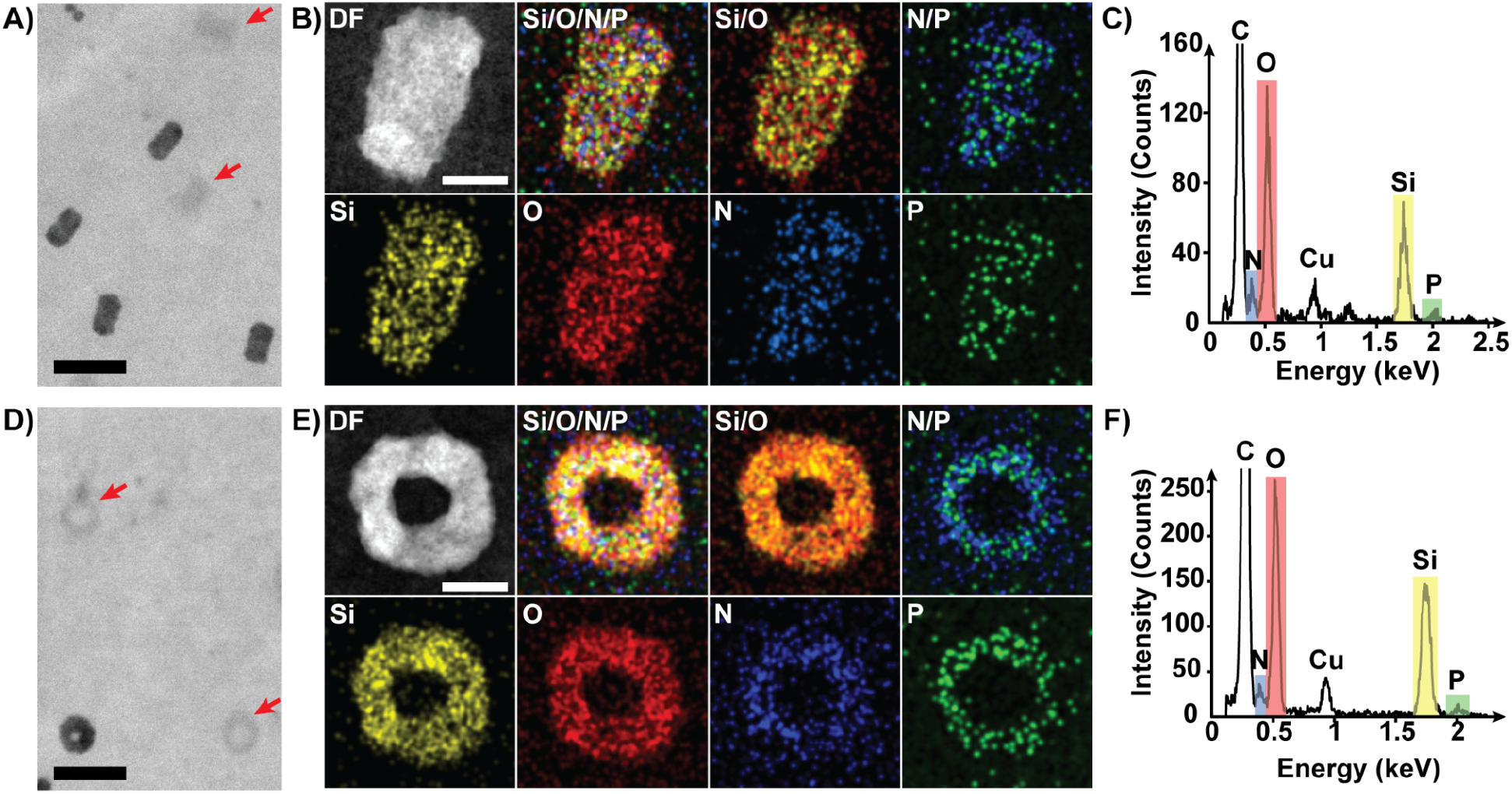
The electron micrographs compare the non-coated control DNA structures with the silica-coated ones (A and D). Scale bars = 100 nm. Energy-Dispersive Spectroscopy (EDS) elemental analysis to confirm inorganic material growth of B-C) Silica coated DNA brick structures and E-F) Silica coated DNA ring structures. B and E) Represent a collection of electron micrographs overlayed with elemental signal maps as mentioned in the top left of each image (Silica, Si = yellow; Oxygen, O = red; Nitrogen, N = blue; Phosphorus, P = green). Scale bars = 20 nm. C and F) Represent the energy-dispersive spectra against intensity, which has been correlated with the respective elements.

In addition, we performed elemental analysis and plotted the energy dispersive spectra (EDS) for both nanoparticles (Figure 5 B-C and E-F). This analysis provided the clearest, unambiguous evidence of the presence of silica around the DNAO. The overlay of the dark field image of the structure with the energy signal from the constituent elements (silicon, oxygen, nitrogen, and phosphorous) confirmed that the silica growth takes place on the DNA origami structures. The dense silicon elemental map (Labeled “Si”) for the DNA brick and ring structures suggested consistent growth. A comparison study between the control DNAO and silica-coated structures on the same grid is in the supporting information (Figure S 27). As expected, the DNAO structures lacked the silicon intensity peak along with the decreased peak for oxygen.

### Robustness

For the entire duration of the tests and optimizations, the silica-templated DNAO nanoparticles exhibited stability for long durations (over 4 days, data not included) in salt-free, aqueous, and organic solvents, where typically, native DNAO nanoparticles either degrade or aggregate. These structures also survived high pH solutions (pH 11) and pH transitions from pH 11 to 7. From these observations, we concluded that the structures could sustain a wide range of working conditions, which enables various applications that do not favor aqueous saline solutions.

### Thermal stability

Next, we evaluated the robustness of silica DNA nanoparticles at elevated temperatures. Typically, most native DNA nanoparticles disintegrate at temperatures above 50 °C in favorable buffer conditions or room temperature in low-salt or no-salt aqueous buffers.^40^ This poses a major challenge for solution-based applications that require elevated working temperatures. To investigate the thermal stability of the silica-coated DNA nanoparticles, we performed the silica shell growth on 42HB DOPMs using 5 kDa PEG length, 125x TMOS as a precursor in 85% ethanol at pH 11. After 5 min of growth, we stopped the reaction using PS, followed by adding HCl after an additional 15 min of incubation. We incubated these structures along with native structures and DOPMs as controls in a thermocycler for 30 min at 90 °C. Upon AGE analysis (Figure S 28), we observed that the structures did not exhibit any staple leakage. However, from the TEM images (Figure S 29), we concluded that the structures lost their shape and resembled rod-like structures. We made these observations for DOPMs and the structure with silica shells and concluded that despite staple retention, there was a loss of structural integrity. We hypothesize that the observed thermal degradation is due to the condensation forces exerted by the 10-unit long polylysines on the DNA being the most prominent component at elevated temperatures, especially since the silica shell is thin and likely to be highly porous. Interestingly, the lack of staple strand leakage suggests that despite the structural distortion, the strong electrostatic attractions of the polylysines retain all the staples strand within the structure. The same behavior is observed with DOPMs (Data not shown). In order to give our templated silica nanoparticles thermal stability, we utilized the well-established glutaraldehyde crosslinking method (that interconnects primary amines on the polylysines)^58^ to reduce the DNA condensation forces and improve the thermal stability. Upon repeating the thermostability test with crosslinked DOPMs (DOPMx), we observed that despite some staple leakage, the structures maintained the shape (Figure S 28 and Figure S 29). These observations confirm our hypothesis, and the reduction in the overall positive charge on the polylysines led to a significant reduction in the condensation forces. To circumvent the observed staple leakage, we incorporated minor design modifications by removing weakly bound staples (staples with domain length of ≤7), which completely stopped the staple strand leakage (Figure S 30.).

### Accessibility of functional groups after silica growth

One of the primary goals of this work was to demonstrate not only the facile, templated growth of silica on DNA origami but also the retention of spatial programmability of the DNA origami. To demonstrate this capability, we initially performed a bulk accessibility test. For this, we utilized azido-modified PEG-PLL (-N_3_ groups on the terminal ends of PEG) to form the DOPMs. The presence of this terminal group enables click-chemistry to a dibenzylcyclooctyne (DBCO) functionalized fluorescent dye. Please check the supporting information for a complete setup description (Figure S 31). We used these azido-DOPMs for silica shell growth using 125x TMOS, 85% ethanol at pH 11, and PS, HCl combination to stop the growth reaction. To assess the accessibility of the azido group after the growth, we added DBCO-functionalized Atto-488 fluorescent dye. We observed the atto-488 fluorescence from the AGE results, and we confirmed that the terminal groups were accessible after the silica growth. This result further supports our hypothesis that the silica deposition occurs directly on the surface of the DNA and does not embed the PEG chains in the process. This result also suggested that by incorporating PEG chains, with terminal reactive groups, at select sites on the DNA, we can create defined reaction sites on the silica-templated nanoparticle.

We modified the DNA brick design to accommodate 1-, 2-, or 3-single-stranded DNA extensions with unique sequences at select locations on the outer surface of the structure (Figure S 32). Next, to these extensions, we added complementary oligonucleotides conjugated with 5 kDa long PEG chains carrying a biotin group at the terminal position. This conjugate can bind to entities carrying streptavidin. A detailed protocol for the synthesis, and purification, of the conjugate, is presented in the supporting information (Figure S 33). We used these DNA brick structures with either 1, 2, or 3-biotin groups (Figure 6 A) to grow silica shells and evaluated the accessibility by adding streptavidin-functionalized 10 nm iron oxide nanoparticles (Strep-IONPs). We performed the silica growth by forming DOPMs with the 5 kDa PEG chain length, using 200x TMOS as the precursor, and conducted the growth reaction for 3 hours. We characterized the resulting structures with TEM, and for reference, we used native DNA nanoparticles and DOPMs without silica shells. From the TEM images, we confirmed the accessibility of Biotin groups across the controls and the silica-coated structures (Figure 6 A and supporting Figure S 34, Figure S 35 and Figure S 36). Further, we observe that the interaction between the nanoparticle and the silica-templated DNA origami is statistically identical to that between the nanoparticle and DNA origami and DNA origami polyplex (Figure 6 B-C). This suggests that while the interaction between the nanoparticle and the terminal reactive group might require improvement, the silica growth process does not adversely affect the accessibility of the reactive groups organized on the DNA nanostructure.

**Figure 6.**
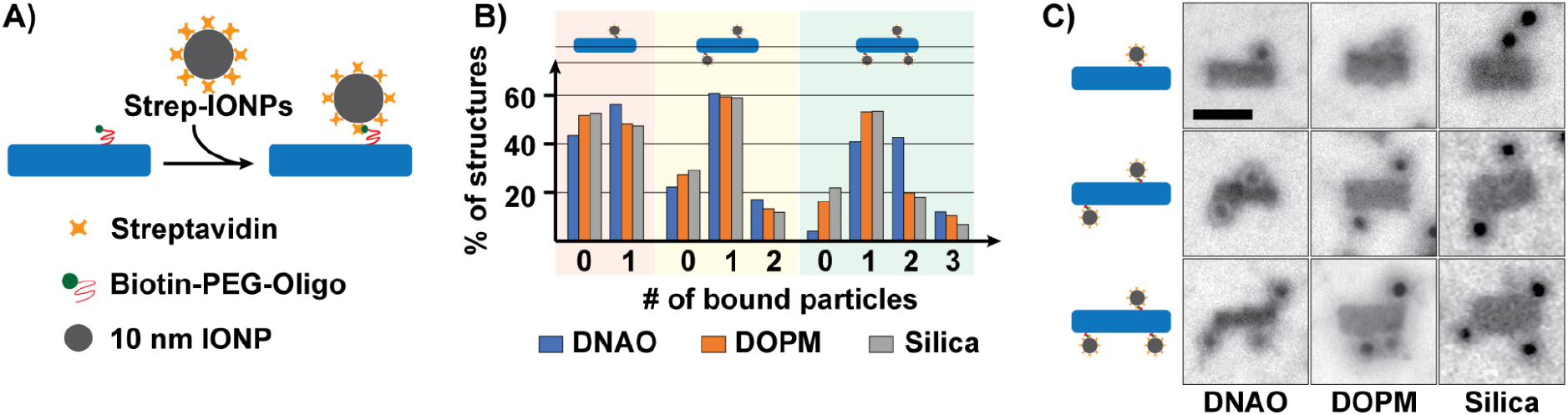
Attaching iron oxide nanoparticles (IONPs) to the functional silica DNA nanoparticle. A) Schematic of the binding strategy. Biotin moieties placed at specific locations on the surface of the DNA nanoparticle were bound to streptavidin-functionalized 10 nm IONPs. B) Binding statistics. C) TEM images representing the three variations of the binding sites. DNAO and DOPMs were used as controls for this study. Scale bars = 50 nm.

## Conclusion

The scaffolded DNA origami, over the last 15 years, has found widespread use as a tool in fundamental research in the areas of biophysics, nanooptics as well as other adjacent fields. However, its utility as a mainstream technique to rival conventional nanoparticles has been inhibited by manufacturing challenges associated with large-scale production of DNA origami, instability of native DNA structures in-vivo as well as DNA being materially fragile. And, only when all these three shortcomings are solved simultaneously will the DNA origami be accepted as a practical nanoparticle synthesis technique. While great strides have been made in mitigating issues associated with large-scale manufacturing of DNA origami as well as increasing the in-vivo stability of DNA origami without losing any of the functional capabilities of DNA origami, our present work is the first attempt solving the material fragility of DNA through controlled inorganic material growth without degrading the salient functional features of DNA origami. Further, the method presented here is easy to implement, has a high yield and is inexpensive both in terms of reaction time as well as capital. The main outstanding issue with our technique is optimization of coupling chemistry to maximize the yield associated with the decoration of nanoparticles and molecules on top of silicified DNA origami.

A few obvious applications that can emerge from our approach include: (a) Rapid silicification or other material growth on DNA origami crystals in the service of developing high tensile strength composite materials, (b) by converting the DNA origami to silica we can leverage the extensive Nanoparticle therapeutics,^59^ (c) inorganic scaffold allows the nanoparticle to act as scaffold for metal growth that in turn enables complex metallic structure for practical optical applications as well as biomedical imaging needs. More generally we see the ease, and speed, associated with the synthesis of our particles to further accelerate the growth and utility of DNA origami.

## Acknowledgements

We acknowledge funding from National Science Foundation (MCB2027165) and DARPA (140D0422C0020). We thank Dr. Aubrey Penn for help with EDS measurements, Dr. Simon Roessler for use of his HPLC purification setup, Ms. Emily Wu for assistance with silica growth measurements and Prof. Giovanni Traverso for the use of Malvern Zetasizer Ultra for DLS experiments. We acknowledge Prof. Ulrich Wiesner for providing valuable feedback regarding the silica growth at the initial stages of the project.

## Contributions

A.G. conceived the project. N.P.A. performed all the experiments and characterization. Both authors contributed to data interpretation and manuscript preparation.

## References

(1) Rothemund, P. W. K. Folding DNA to Create Nanoscale Shapes and Patterns. Nature 2006, 440 (7082), 297–302. https://doi.org/10.1038/nature04586.

(2) He, Y.; Ye, T.; Su, M.; Zhang, C.; Ribbe, A. E.; Jiang, W.; Mao, C. Hierarchical Self-Assembly of DNA into Symmetric Supramolecular Polyhedra. Nature 2008, 452 (7184), 198–201. https://doi.org/10.1038/nature06597.

(3) Douglas, S. M.; Dietz, H.; Liedl, T.; Högberg, B.; Graf, F.; Shih, W. M. Self-Assembly of DNA into Nanoscale Three-Dimensional Shapes. Nature 2009, 459 (7245), 414–418. https://doi.org/10.1038/nature08016.

(4) Matthies, M.; Agarwal, N. P.; Schmidt, T. L. Design and Synthesis of Triangulated DNA Origami Trusses. Nano Lett. 2016, 16 (3), 2108–2113. https://doi.org/10.1021/acs.nanolett.6b00381.

(5) Wagenbauer, K. F.; Sigl, C.; Dietz, H. Gigadalton-Scale Shape-Programmable DNA Assemblies. Nature 2017, 552 (7683), 78–83. https://doi.org/10.1038/nature24651.

(6) Tikhomirov, G.; Petersen, P.; Qian, L. Fractal Assembly of Micrometre-Scale DNA Origami Arrays with Arbitrary Patterns. Nature 2017, 552 (7683), 67. https://doi.org/10.1038/nature24655.

(7) Matthies, M.; Agarwal, N. P.; Poppleton, E.; Joshi, F. M.; Šulc, P.; Schmidt, T. L. Triangulated Wireframe Structures Assembled Using Single-Stranded DNA Tiles. ACS Nano 2019, 13 (2), 1839–1848. https://doi.org/10.1021/acsnano.8b08009.

(8) Dey, S.; Fan, C.; Gothelf, K. V.; Li, J.; Lin, C.; Liu, L.; Liu, N.; Nijenhuis, M. A. D.; Saccà, B.; Simmel, F. C.; Yan, H.; Zhan, P. DNA Origami. Nat. Rev. Methods Primer 2021, 1 (1), 1–24. https://doi.org/10.1038/s43586-020-00009-8.

(9) Hong, F.; Zhang, F.; Liu, Y.; Yan, H. DNA Origami: Scaffolds for Creating Higher Order Structures. Chem. Rev. 2017, 117 (20), 12584–12640. https://doi.org/10.1021/acs.chemrev.6b00825.

(10) Madsen, M.; Gothelf, K. V. Chemistries for DNA Nanotechnology. Chem. Rev. 2019, 119 (10), 6384–6458. https://doi.org/10.1021/acs.chemrev.8b00570.

(11) Kuzyk, A.; Schreiber, R.; Fan, Z.; Pardatscher, G.; Roller, E.-M.; Högele, A.; Simmel, F. C.; Govorov, A. O.; Liedl, T. DNA-Based Self-Assembly of Chiral Plasmonic Nanostructures with Tailored Optical Response. Nature 2012, 483 (7389), 311–314. https://doi.org/10.1038/nature10889.

(12) Mikkilä, J.; Eskelinen, A.-P.; Niemelä, E. H.; Linko, V.; Frilander, M. J.; Törmä, P.; Kostiainen, M. A. Virus-Encapsulated DNA Origami Nanostructures for Cellular Delivery. Nano Lett. 2014, 14 (4), 2196–2200. https://doi.org/10.1021/nl500677j.

(13) Gopinath, A.; Miyazono, E.; Faraon, A.; Rothemund, P. W. K. Engineering and Mapping Nanocavity Emission via Precision Placement of DNA Origami. Nature 2016, 535 (7612), 401–405. https://doi.org/10.1038/nature18287.

(14) Liu, N.; Liedl, T. DNA-Assembled Advanced Plasmonic Architectures. Chem. Rev. 2018, 118 (6), 3032–3053. https://doi.org/10.1021/acs.chemrev.7b00225.

(15) Linko, V.; Ora, A.; Kostiainen, M. A. DNA Nanostructures as Smart Drug-Delivery Vehicles and Molecular Devices. Trends Biotechnol. 2015, 33 (10), 586–594. https://doi.org/10.1016/j.tibtech.2015.08.001.

(16) Tian, Y.; Lhermitte, J. R.; Bai, L.; Vo, T.; Xin, H. L.; Li, H.; Li, R.; Fukuto, M.; Yager, K. G.; Kahn, J. S.; Xiong, Y.; Minevich, B.; Kumar, S. K.; Gang, O. Ordered Three-Dimensional Nanomaterials Using DNA-Prescribed and Valence-Controlled Material Voxels. Nat. Mater. 2020, 19 (7), 789–796. https://doi.org/10.1038/s41563-019-0550-x.

(17) Kahn, J. S.; Gang, O. Designer Nanomaterials through Programmable Assembly. Angew. Chem. 2022, 134 (3), e202105678. https://doi.org/10.1002/ange.202105678.

(18) Praetorius, F.; Kick, B.; Behler, K. L.; Honemann, M. N.; Weuster-Botz, D.; Dietz, H. Biotechnological Mass Production of DNA Origami. Nature 2017, 552 (7683), 84. https://doi.org/10.1038/nature24650.

(19) Shetty, R. M.; Brady, S. R.; Rothemund, P. W. K.; Hariadi, R. F.; Gopinath, A. Bench-Top Fabrication of Single-Molecule Nanoarrays by DNA Origami Placement. ACS Nano 2021, 15 (7), 11441–11450. https://doi.org/10.1021/acsnano.1c01150.

(20) Gopinath, A.; Thachuk, C.; Mitskovets, A.; Atwater, H. A.; Kirkpatrick, D.; Rothemund, P. W. K. Absolute and Arbitrary Orientation of Single-Molecule Shapes. Science 2021, 371 (6531), eabd6179. https://doi.org/10.1126/science.abd6179.

(21) Gartner, Z. J.; Tse, B. N.; Grubina, R.; Doyon, J. B.; Snyder, T. M.; Liu, D. R. DNA-Templated Organic Synthesis and Selection of a Library of Macrocycles. Science 2004, 305 (5690), 1601–1605. https://doi.org/10.1126/science.1102629.

(22) Rozenman, M. M.; Liu, D. R. DNA-Templated Synthesis in Organic Solvents. ChemBioChem 2006, 7 (2), 253–256. https://doi.org/10.1002/cbic.200500413.

(23) McKee, M. L.; Milnes, P. J.; Bath, J.; Stulz, E.; Turberfield, A. J.; O’Reilly, R. K. Multistep DNA-Templated Reactions for the Synthesis of Functional Sequence Controlled Oligomers. Angew. Chem. Int. Ed. 2010, 49 (43), 7948–7951. https://doi.org/10.1002/anie.201002721.

(24) He, Y.; Liu, D. R. Autonomous Multistep Organic Synthesis in a Single Isothermal Solution Mediated by a DNA Walker. Nat. Nanotechnol. 2010, 5 (11), 778–782. https://doi.org/10.1038/nnano.2010.190.

(25) Acuna, G. P.; Bucher, M.; Stein, I. H.; Steinhauer, C.; Kuzyk, A.; Holzmeister, P.; Schreiber, R.; Moroz, A.; Stefani, F. D.; Liedl, T.; Simmel, F. C.; Tinnefeld, P. Distance Dependence of Single-Fluorophore Quenching by Gold Nanoparticles Studied on DNA Origami. ACS Nano 2012, 6 (4), 3189–3195. https://doi.org/10.1021/nn2050483.

(26) Bathe, M.; Rothemund, P. W. K. DNA Nanotechnology: A Foundation for Programmable Nanoscale Materials. MRS Bull. 2017, 42 (12), 882–888. https://doi.org/10.1557/mrs.2017.279.

(27) Stöber, W.; Fink, A.; Bohn, E. Controlled Growth of Monodisperse Silica Spheres in the Micron Size Range. J. Colloid Interface Sci. 1968, 26 (1), 62–69.

(28) Darr, J. A.; Zhang, J.; Makwana, N. M.; Weng, X. Continuous Hydrothermal Synthesis of Inorganic Nanoparticles: Applications and Future Directions. Chem. Rev. 2017, 117 (17), 11125–11238. https://doi.org/10.1021/acs.chemrev.6b00417.

(29) Ahn, T.; Kim, J. H.; Yang, H.-M.; Lee, J. W.; Kim, J.-D. Formation Pathways of Magnetite Nanoparticles by Coprecipitation Method. J. Phys. Chem. C 2012, 116 (10), 6069–6076. https://doi.org/10.1021/jp211843g.

(30) López-Quintela, M. A. Synthesis of Nanomaterials in Microemulsions: Formation Mechanisms and Growth Control. Curr. Opin. Colloid Interface Sci. 2003, 8 (2), 137–144. https://doi.org/10.1016/S1359-0294(03)00019-0.

(31) Mertig, M.; Colombi Ciacchi, L.; Seidel, R.; Pompe, W.; De Vita, A. DNA as a Selective Metallization Template. Nano Lett. 2002, 2 (8), 841–844. https://doi.org/10.1021/nl025612r.

(32) Sun, W.; Boulais, E.; Hakobyan, Y.; Wang, W. L.; Guan, A.; Bathe, M.; Yin, P. Casting Inorganic Structures with DNA Molds. Science 2014, 346 (6210). https://doi.org/10.1126/science.1258361.

(33) Liu, X.; Zhang, F.; Jing, X.; Pan, M.; Liu, P.; Li, W.; Zhu, B.; Li, J.; Chen, H.; Wang, L.; Lin, J.; Liu, Y.; Zhao, D.; Yan, H.; Fan, C. Complex Silica Composite Nanomaterials Templated with DNA Origami. Nature 2018, 559 (7715), 593–598. https://doi.org/10.1038/s41586-018-0332-7.

(34) Nguyen, L.; Döblinger, M.; Liedl, T.; Heuer-Jungemann, A. DNA-Origami-Templated Silica Growth by Sol–Gel Chemistry. Angew. Chem. Int. Ed. 2019, 58 (3), 912–916. https://doi.org/10.1002/anie.201811323.

(35) Nguyen, M.-K.; Nguyen, V. H.; Natarajan, A. K.; Huang, Y.; Ryssy, J.; Shen, B.; Kuzyk, A. Ultrathin Silica Coating of DNA Origami Nanostructures. Chem. Mater. 2020, acs.chemmater.0c02111. https://doi.org/10.1021/acs.chemmater.0c02111.

(36) Aryal, B. R.; Ranasinghe, D. R.; Westover, T. R.; Calvopiña, D. G.; Davis, R. C.; Harb, J. N.; Woolley, A. T. DNA Origami Mediated Electrically Connected Metal—Semiconductor Junctions. Nano Res. 2020, 13 (5), 1419–1426. https://doi.org/10.1007/s12274-020-2672-5.

(37) Pang, C.; Aryal, B. R.; Ranasinghe, D. R.; Westover, T. R.; Ehlert, A. E. F.; Harb, J. N.; Davis, R. C.; Woolley, A. T. Bottom-Up Fabrication of DNA-Templated Electronic Nanomaterials and Their Characterization. Nanomaterials 2021, 11 (7), 1655. https://doi.org/10.3390/nano11071655.

(38) Majewski, P. W.; Michelson, A.; Cordeiro, M. A. L.; Tian, C.; Ma, C.; Kisslinger, K.; Tian, Y.; Liu, W.; Stach, E. A.; Yager, K. G.; Gang, O. Resilient Three-Dimensional Ordered Architectures Assembled from Nanoparticles by DNA. Sci. Adv. 2021, 7 (12), eabf0617. https://doi.org/10.1126/sciadv.abf0617.

(39) Hahn, J.; Wickham, S. F. J.; Shih, W. M.; Perrault, S. D. Addressing the Instability of DNA Nanostructures in Tissue Culture. ACS Nano 2014, 8 (9), 8765–8775. https://doi.org/10.1021/nn503513p.

(40) Gerling, T.; Kube, M.; Kick, B.; Dietz, H. Sequence-Programmable Covalent Bonding of Designed DNA Assemblies. Sci. Adv. 2018, 4 (8), eaau1157. https://doi.org/10.1126/sciadv.aau1157.

(41) Kielar, C.; Xin, Y.; Shen, B.; Kostiainen, M. A.; Grundmeier, G.; Linko, V.; Keller, A. On the Stability of DNA Origami Nanostructures in Low-Magnesium Buffers. Angew. Chem. Int. Ed. 2018, 57 (30), 9470–9474. https://doi.org/10.1002/anie.201802890.

(42) Wu, S.-H.; Mou, C.-Y.; Lin, H.-P. Synthesis of Mesoporous Silica Nanoparticles. Chem. Soc. Rev. 2013, 42 (9), 3862–3875. https://doi.org/10.1039/C3CS35405A.

(43) Kankala, R. K.; Han, Y.-H.; Na, J.; Lee, C.-H.; Sun, Z.; Wang, S.-B.; Kimura, T.; Ok, Y. S.; Yamauchi, Y.; Chen, A.-Z.; Wu, K. C.-W. Nanoarchitectured Structure and Surface Biofunctionality of Mesoporous Silica Nanoparticles. Adv. Mater. 2020, 32 (23), 1907035. https://doi.org/10.1002/adma.201907035.

(44) Brinker, C. J.; Scherer, G. W. Sol-Gel Science: The Physics and Chemistry of Sol-Gel Processing; Academic Press, 2013.

(45) Issa, A. A.; Luyt, A. S. Kinetics of Alkoxysilanes and Organoalkoxysilanes Polymerization: A Review. Polymers 2019, 11 (3), 537. https://doi.org/10.3390/polym11030537.

(46) Agarwal, N. P.; Matthies, M.; Gür, F. N.; Osada, K.; Schmidt, T. L. Block Copolymer Micellization as a Protection Strategy for DNA Origami. Angew. Chem. Int. Ed. 2017, 56 (20), 5460–5464. https://doi.org/10.1002/anie.201608873.

(47) Ponnuswamy, N.; Bastings, M. M. C.; Nathwani, B.; Ryu, J. H.; Chou, L. Y. T.; Vinther, M.; Li, W. A.; Anastassacos, F. M.; Mooney, D. J.; Shih, W. M. Oligolysine-Based Coating Protects DNA Nanostructures from Low-Salt Denaturation and Nuclease Degradation. Nat. Commun. 2017, 8 (1), 15654. https://doi.org/10.1038/ncomms15654.

(48) Agarwal, N. P.; Chandrasekhar, S.; Prakash, P. S.; Joffroy, K.; Schmidt, T. L. Block Copolymer Micellization of DNA Origami Promotes Solubility in Organic Solvents. Langmuir 2022, 38 (38), 11650–11657. https://doi.org/10.1021/acs.langmuir.2c01508.

(49) Engelhardt, F. A. S.; Praetorius, F.; Wachauf, C. H.; Brüggenthies, G.; Kohler, F.; Kick, B.; Kadletz, K. L.; Pham, P. N.; Behler, K. L.; Gerling, T.; Dietz, H. Custom-Size, Functional, and Durable DNA Origami with Design-Specific Scaffolds. ACS Nano 2019, 13 (5), 5015–5027. https://doi.org/10.1021/acsnano.9b01025.

(50) Dietz, H.; Douglas, S. M.; Shih, W. M. Folding DNA into Twisted and Curved Nanoscale Shapes. Science 2009, 325 (5941), 725–730. https://doi.org/10.1126/science.1174251.

(51) Roodhuizen, J. A. L.; Hendrikx, P. J. T. M.; Hilbers, P. A. J.; de Greef, T. F. A.; Markvoort, A. J. Counterion-Dependent Mechanisms of DNA Origami Nanostructure Stabilization Revealed by Atomistic Molecular Simulation. ACS Nano 2019, 13 (9), 10798–10809. https://doi.org/10.1021/acsnano.9b05650.

(52) Bertosin, E.; Stömmer, P.; Feigl, E.; Wenig, M.; Honemann, M. N.; Dietz, H. Cryo-Electron Microscopy and Mass Analysis of Oligolysine-Coated DNA Nanostructures. ACS Nano 2021, 15 (6), 9391–9403. https://doi.org/10.1021/acsnano.0c10137.

(53) Hansen, C. M. Hansen Solubility Parameters: A User’s Handbook, 2nd ed.; CRC Press: Boca Raton, 2007.

(54) Sun, Y.; Sai, H.; von Stein, F.; Riccio, M.; Wiesner, U. Water-Based Synthesis of Ultrasmall PEGylated Gold–Silica Core–Shell Nanoparticles with Long-Term Stability. Chem. Mater. 2014, 26 (18), 5201–5207. https://doi.org/10.1021/cm501348r.

(55) Ma, K.; Mendoza, C.; Hanson, M.; Werner-Zwanziger, U.; Zwanziger, J.; Wiesner, U. Control of Ultrasmall Sub-10 Nm Ligand-Functionalized Fluorescent Core–Shell Silica Nanoparticle Growth in Water. Chem. Mater. 2015, 27 (11), 4119–4133. https://doi.org/10.1021/acs.chemmater.5b01222.

(56) Wu, W.-C.; Tracy, J. B. Large-Scale Silica Overcoating of Gold Nanorods with Tunable Shell Thicknesses. Chem. Mater. 2015, 27 (8), 2888–2894. https://doi.org/10.1021/cm504764v.

(57) Fragasso, A.; De Franceschi, N.; Stömmer, P.; van der Sluis, E. O.; Dietz, H.; Dekker, C. Reconstitution of Ultrawide DNA Origami Pores in Liposomes for Transmembrane Transport of Macromolecules. ACS Nano 2021, acsnano.1c01669. https://doi.org/10.1021/acsnano.1c01669.

(58) Anastassacos, F. M.; Zhao, Z.; Zeng, Y.; Shih, W. M. Glutaraldehyde Cross-Linking of Oligolysines Coating DNA Origami Greatly Reduces Susceptibility to Nuclease Degradation. J. Am. Chem. Soc. 2020, 142 (7), 3311–3315. https://doi.org/10.1021/jacs.9b11698.

(59) Janjua, T. I.; Cao, Y.; Yu, C.; Popat, A. Clinical Translation of Silica Nanoparticles. Nat. Rev. Mater. 2021, 6 (12), 1072–1074. https://doi.org/10.1038/s41578-021-00385-x.

