## Supporting information for "DNA origami 2.0"

#### SUPPORTING INFORMATION

**This PDF file includes:**

Materials and Methods

Figures S1 to S36

References and Notes

### Methods and materials.

#### DNA origami design.

The DNA brick (42HBs) design described previously[1] was used without any design modification (sequences and the caDNAno design occurs as 42HB_staples.xls and 42HB.json, as a part of the zip archive 42HBdesgin.zip). Briefly, the design consists of a total of 213 oligonucleotides (staple strands), distributed across three 96-well plates at a concentration of 200 µM each. These oligonucleotides were pooled to a stock solution and diluted to a working concentration of 800 nM each. The design was modified to accommodate staple extensions at three locations as described in the Figure S 32. These extensions were 20 bases long and had the following sequence: "CAACCTCATACCACTACAAC", while the complementary oligonucleotide had the following sequence: "GTTGTAGTGGTATGAGGTTG".

The DNA ring (delftagon) design described previously[2] was used without any design modification (sequences and the caDNAno design occurs as ring_staples.xls and ring.json, as a part of the zip archive Ringdesign.zip). Briefly, the design consists of a total of 240 staple strands, distributed across three 96-well plates at a concentration of 200 µM each. These oligonucleotides were pooled to a stock solution and diluted to a working concentration of 1600 nM each.

#### DNA origami folding.

In the folding reaction for DNA brick, 20 nM of p7560 scaffold (Tilibit nanosystems GmbH, Germany), respective staple strand set (Integrated DNA Technologies, IDT) at 100 nM (each staple, a 5:1 staple to scaffold ratio), in 1x Tris EDTA buffer, pH 8.0 at 25 °C (Sigma Aldrich, 10 mM Tris, 1 mM EDTA) with 15 mM magnesium chloride (Sigma Aldrich) were mixed. The mixture was annealed in a thermal cycler (Bio-Rad C1000 Touch) for 65 °C for 15 min for denaturation, followed by a folding ramp from 60 to 20 °C at the rate of -1 °C per 3 min and held at 20 °C for storage.

For DNA ring, the reaction mixture consisted of 10 nM of p7560 scaffold, staple mix at 300 nM (30:1 ratio), in 1x Tris EDTA buffer with 24 mM magnesium chloride. The mixture was annealed for 65 °C for 15 min for denaturation, followed by a 16-hour hold at 50 °C. The folded structures were stored at 20 °C until further purification.

Excess staple strands were then removed by ultrafiltration columns (50 kDa MWCO Amicon Ultra-0.5 Centrifugal Filter unit, AMD Millipore).

#### Purification with ultrafiltration columns.

**Passivation.** Prior to purification, the columns were passivated by adding 500 µL of 5% Pluronic F-127 (Sigma Aldrich) solution to the filters and incubating for 30 min at room temperature. After the incubation, the filters were flipped to discard the solution and spun in the flipped state for 1 min at 2,000 g to remove any remaining solution. The filters were then washed with 500 µL of ultrapure water and flipped to discard the solution. This step was repeated for a total of four times, after which the filter was used right away.

NOTE 1: Do not leave the filter unused or without any solution for a long duration as the filter membrane can dry out, leading to poor yields.

NOTE 2: Do not let the Pluronic solution pass through the filter, as it will block the pore completely.

**Purification.** To the passivated filter, the crude DNAO mixture was added along with the wash buffer (1x Tris EDTA, pH 8.0 with 5 mM $MgCl_2$) to bring the total volume to 400 µL. The filter was spun at 14,000 g for 2 min at room temperature, the flow through was discarded and 400 µL of the wash buffer was added. The total washes were calculated as per the total staple concentration. Typically, 8 washes were performed for 200 µL of the impure reaction mixture consisting of 100 nM of each staple. The last wash step was performed for 5 min to obtain a concentrated purified mixture. After the final wash the filter was reversed, placed in a fresh tube, and centrifuged at 1,000 rcf for 2 min. The purified DNA origami solution was then collected for further characterization and reactions.

NOTE 3: Add the wash buffer prior to the addition of the crude mixture. E.g.: If the DNAO mixture if 40 µL, add 360 µL of the wash buffer to the filter followed by the DNA mixture.

NOTE 4: Do not have more than 50 fmoles of total staple concentration in the filter as it leads to poor purification yields. Use multiple filters if necessary.

#### Preparation of DOPMs.

The PEG-PLys with varying PEG chain lengths (1, 5 or 20 kDa) or the terminal azide group (N3-PEG-PLys with 5 kDa PEG chain length) were purchased from Alamanda Polymers Inc. It is composed of a PEG segment and a cationic PLys segment with DP 10.

##### DNA origami polyplex micelle preparation.

**Standard preparation.** For the preparation of DOPMs, the polyplex micellization strategy was used as described previously.[3] For this, the block copolymer solution was added at the desired N/P ratio to DNA origami solution and the mixture was immediately vortexed. The required volume of PEG-PLys needed for DOPM formation were calculated using the excel calculator (calculator in the supporting files). After a brief incubation the DOPM solution was purified to remove excess PEG-PLys, and buffer exchanged to ultrapure water.

##### Purification of DOPMs using ultrafiltration columns.

For purifying the mixture after complexation and to remove excess PEG-PLys, the passivation and purification protocol as mentioned earlier were used. For purification, 400 µl of ultrapure water were used instead and a total of 4 washes were performed to completely remove salts. After the final wash, the filters were reversed, placed in a fresh tube, and centrifuged at 1,000 rcf for 2 min. The purified DOPMs solution was then collected and used further.

##### Glutaraldehyde crosslinking of polylysines.

Purified DOPMs in ultrapure water were used for glutaraldehyde crosslinking. For this the N/P ratio was assumed to be 2 after purification and accordingly, 200 times glutaraldehyde (50% in water, Grade 1 for electron microscopy, Sigma Aldrich) per amine was added to the DOPMs and mixed thoroughly. The reaction mixture was incubated at room temperature overnight. The crosslinked DOPMs (DOPMx) were purified using Pluronic passivated 50 kDa ultrafiltration columns using the protocol mentioned earlier. Briefly, three washes were performed with 450 µL ultrapure water to obtained purified DOPMx.

#### Silica growth on DOPMs.

**Transfer to 85% ethanol, pH 11.** DOPMs in ultrapure water at high concentrations (>50 nM) were diluted to a final concentration of 2.5 nM using non-denatured ethanol (200% proof for molecular biology, Sigma Aldrich) to get the final ethanol content to 85%. Next, ammonium hydroxide (28.0-30.0% $NH_3$ basis, ACS reagent, Sigma Aldrich) was added to a final concentration of 0.14 M to get to pH 11. For this, 695 µL were taken in 250 µL water to get 5 M ammonium hydroxide solution. This mixture was vortexed thoroughly to obtain a homogenous mixture.

**Addition of precursor.** To the above mixture, precursor (TMOS, 99% Thermo Scientific; or TEOS, reagent grade 98% Sigma Aldrich) was added to initiate silica growth and mixed immediately. The mixture was incubated at room temperature. For experiments with DLS measurements, the mixture was immediately used for measurement after the precursor addition. Dilution of the precursor was performed using dimethyl sulfoxide (DMSO, 99.9% ACS reagent Sigma Aldrich). The diluted precursors were always freshly prepared and used immediately.

**Calculations for the precursor concentration.** From several preliminary tests (data not shown), the precursor concentration was adapted to an arbitrary number of 9,300 molecules per $nm^2$ of origami surface, which corresponded to 37.2 mM precursor per 1 nM for DNA origami. This amount is denoted as 1x precursor concentration. Any further dilution factor of the precursor is denoted with respective number, for instance 10x or 125x dilution for 10 times or 125 times dilution to the above mentioned 37.2 mM precursor concentration. The appropriate dilution factor was worked out by observing the DLS results. Further, the dilution factor can be adjusted as per the requirement of the silica coating thickness.

**Variations.** Several adjustments were made to the above protocols to verify the optimal growth conditions. In all the instances, the growth protocol was maintained while changing single parameters. For experiments with methanol, like ethanol, DOPMs in ultrapure water at high concentrations were diluted using pure methanol (99.9%, HPLC grade, Sigma Aldrich). For experiments requiring pH transition from pH 11 to 7, hydrochloric acid (37%, reagent grade, Sigma Aldrich) were used. Final concentration of 0.2 M of HCl were added to 100 µL of the reaction mixture and mixed thoroughly.

**Stopping the growth.** For stopping the growth reaction PEG-silane (3-[Methoxy(polyethyleneoxy)$_{6\text{-}9}$] propyltrimethoxysilane, tech grade, Gelest, Inc.) was used. Any necessary dilutions were performed using dimethyl sulfoxide (DMSO, 99.9% ACS reagent Sigma Aldrich). The diluted solutions were always freshly prepared and used immediately.

**Purification/moving to ultrapure water.** For purifying the silica structures from unreacted excess reagents, 350 µL of 85% ethanol, pH 11 wash buffer was added to 100 µL of the reaction mixture. This solution was added to Pluronic passivated 50 kDa ultrafiltration column and spun at 14,000 g for 2 min at room temperature. The flow-through was discarded and 400 µL of fresh wash buffer was added. This wash step was repeated for a total of three times followed by another three washes with 400 µL of ultrapure water for buffer exchange. After the final wash the filter was reversed, placed in a fresh tube, and centrifuged at 1,000 rcf for 2 min. The purified DNA origami solution was then collected for further characterization and reactions.

#### Agarose gel electrophoresis.

For the AGE analysis, 0.3% agarose gels (Type I, Sigma Aldrich) were casted with 0.5x Tris Borate EDTA buffer (prepared used 10x Tris-borate-EDTA buffer, ultra-pure grade VWR life science) containing 12 mM $MgCl_2$ and pre-stained with 1x Sybr safe DNA gel stain (Invitrogen by ThermoFisher Scientific). As a running buffer, 0.5x TBE buffer with 12 mM $MgCl_2$ was used. For sample preparation, 15 µL of sample solution were mixed with 3 µL of 6x gel loading dye (15% Ficoll 400, 5 mM Tris, 1 mM EDTA, 12 mM $MgCl_2$, 0.03% bromophenol blue and 0.03% of xylene cyanol). Electrophoresis was performed at 70 V for 2 h at room temperature. As a reference, 3 µL of 1 kb plus DNA ladder (ThermoFisher Scientific) were added. The gel was imaged with the UVP Gel Studio gel scanner (Analytik Jena GmbH). For experiments without salt, the gel and the running buffers were prepared without $MgCl_2$.

#### Dynamic light scattering measurements.

All the measurements were performed using the Zetasizer Ultra (Zetasizer advanced range, Malvern Panalytical). Low volume disposable cuvettes were used with adaptors to avoid any evaporation during measurements (ZEN0040 UV-Cuvette with 8.5 mm window height, BrandTech Brand). All the samples were equilibrated at 25 °C for all measurements. All the results were analyzed using the ZS xplorer software available at the manufacturer website. For all the control measurements, the size measurements were performed 10 times. For all the growth reactions, the number of repeats were set as per the desired incubation time.

#### tSEM characterization.

Carbon-coated TEM grids (400 mesh copper, formvar/carbon film 10 nm/1 nm thick, Electron Microscopy Sciences) were plasma-treated for 15 s. 2 µL of the sample solution were drop casted on the grid and incubated for 5 min. The excess solution was removed from the grid with a filter paper. Next, 5 µL of a 2% uranyl formate solution was applied for 90 s to stain the DNA origami structures, and the solution was removed with a filter paper. Additionally, to remove excess stain, two washing steps were performed with 10 µL of ultrapure water for 10 s each and the water removed with a filter paper. The samples were scanned on Gemini SEM450 (Zeiss) operated at 20 kV.

**Preparation of 2% uranyl formate solution.** 100 mg of uranyl formate (Electron Microscopy Sciences) were added to 5 ml of boiled ultrapure water. The mixture was stirred for a minimum of 5 min and filtered using a 0.22 µm syringe filter. The filtered solution was stored at -20 °C in 100 µL aliquots. Prior to staining, 2.5 µL of 1 M NaOH solution were added to 100 µL aliquot of 2 % uranyl formate and mixed at room temperature for 5 min.

**Semiautomated size measurements.**

TEM images were used to plot grey value profiles using imageJ ROI manager. The grey-values profiles were further analyzed by python script (https://github.com/ewuirl/TEM-ruler) to obtain average size distribution.

#### Scanning Transmission Electron Microscopy and Energy-Dispersive X-ray Spectroscopy.

Scanning TEM and elemental mapping of DNAO and silica structures was performed at 200 kV using a FEI Titan Themis equipped with a SuperX EDX detector in annular dark field (ADF) mode.

#### PAGE gel analysis.

**Casting 15% denaturing PAGE gel.** 18.2 g of Urea (BioXtra, pH 7.5-9.5 (20 °C, 5 M in H2O), Sigma Aldrich) was dissolved in 20 mL of ultrapure water. To the mixture following components were added: 5 mL of 10x Tris-EDTA-Borate buffer, 18.75 mL of Acrylamide/Bis-acrylamide solution (Ambion, 19:1 40% w/v solution, Invitrogen, Thermofisher Scientific), 400 µL of freshly dissolved Ammonium persulfate, 10% w/v (for molecular biology, >98% Sigma Aldrich), 30 µL of N,N,N′,N′-Tetramethylethylenediamine (TEMED, Surecast, Invitrogen, Thermofisher Scientific). This solution was mixed by inverting the tube. Then the solution was carefully poured into empty PAGE cassettes (1.5 mm thickness, mini, ThermoFisher Scientific). The prepared volume was sufficient for 4 cassettes. Combs were carefully placed into the cassettes without spilling the solution. The cassettes were incubated at room temperature for 30 min. For storage, the cassettes were placed in a bag with 0.5x TBE buffer and placed at 4 °C until further usage.

**Sample loading preparation.** 5 µL of the sample were mixed with 5 µL of 2x denaturing loading solution (50% formamide, 10 mM NaOH, traces of bromophenol blue and xylene cyanol). For the ladder, 0.2 µL of Ultra low Range DNA ladder (ThermoFisher Scientific) were mixed with 0.3 µL 10X folding buffer, 5 µL of 2X denaturing loading dye and 4.7 µL of ultrapure water. The empty lanes were filled with a blank solution composed of 5 µL of 2x loading dye and 5 µL of ultrapure water.

**Protocol for running the gel.** 1x TBE buffer warmed to ~65 °C was used as the running buffer. The lanes were rinsed with 1x TBE buffer to remove excess urea. The gel was pre-run at 230 V for 30 min inside a thermal box filled with hot water (~65 °C). The lanes were rinsed again prior to loading the prepared samples. The gel was run under the same condition as mentioned before. After the run, the gel was post-stained with SYBR gold nucleic acid gel stain (10,000x in DMSO, Invitrogen, ThermoFisher Scientific). The gel staining solution was prepared by mixing 45 mL ultrapure water, 5 mL absolute ethanol. To this mixture 5 µL of the SYBR gold dye was added. The gel was imaged with the UVP Gel Studio gel scanner.

#### Thermal stability tests.

For heat denaturation tests, the reaction mixture containing DNAO or DOPMs or silica structures were incubated in a thermocycler for 30 min at 90 °C. After the incubation, the solutions were placed in -20 °C freezer for 10 min to stop any further degradation. The solutions were immediately used for characterization with AGE and TEM.

**Preparation of fluorescently labelled block copolymers.**

For preparing fluorescently labelled PEG-PLys, terminal azido functionalized PEG-PLys with 5 kDa PEG chain lengths were used. As a fluorescent label, DBCO-PEG$_4$-Atto448 (Lumiprobe) was used. The obtained powder from the manufacturer was dissolved in DMSO to obtain 10 mM stock solution, which was stored in -20 °C freezer and wrapped in a foil to avoid any light exposure.

**Labelling reaction prior polyplex micellization.** For the click reaction, calculations were performed to fluorescently label 50% of the PEG-PLys to avoid any additional purification steps. The reaction mixture was incubated overnight at room temperature and wrapped in a foil to avoid any light exposure. The fluorescently labelled PEG-PLys were used without any purification for DOPM formation. However, purification steps were performed to remove excess PEG-PLys and for buffer exchange to ultrapure water.

**Labelling reaction after polyplex micellization or after silica growth.** Similar calculations were performed with the aim to label 50% PEG-PLys on purified DOPMs or silica structures in ultrapure water. After the addition of the dye, the reaction mixture was incubated overnight at room temperature. The fluorescently tagged structures were purified using ultrafiltration columns with a total of three wash steps with ultrapure water.

**AGE analysis.** The structures were analyzed using 0.3% agarose gels not pre-stained with DNA dye. The rest of the protocol was like the one mentioned earlier. The gel was scanned for Atto488 signal, followed by post-staining with SYBR safe DNA gel stain dye. For this, 120 mL of 0.5x TBE buffer was mixed with 12 µL of DNA stain. The gel was incubated in the staining solution for 30 min at room temperature. Prior to imaging, excess stain was removed by washing the gel with ultrapure water. This step was performed to reduce any background noise.

NOTE: Wash all the gel chamber and combs with 70% ethanol to remove any previous excess of the gel staining dye. This was performed to reduce the occurrence of any false positives.

#### Preparation and purification of DBCO or biotin-oligonucleotide-PEG conjugates.

For this, oligonucleotides with sequences complimentary to the single stranded extensions on the DNAO were order with amino terminal groups from IDT. In addition, DBCO-PEG-NHS, and Biotin-PEG-NHS, with 5 kDa PEG chain lengths were obtained from JenKem Technology USA.

**Click chemistry between amino-oligo and NHS-PEG-DBCO or biotin.** The reaction was performed in 50% DMSO, at pH 8.3 buffered using 1 M sodium bicarbonate solution. 25 times excess of NHS over amino was added to ensure high yields. Prior to the reaction the heterobifunctional PEG was dissolved in DMSO to prepare a 10 mM stock solution. The reaction was incubated overnight at room temperature. The resulting mixture was characterized by PAGE gel using the protocol mentioned above.

**HPLC purification.** For purification, the crude reaction mixture was lyophilized overnight. Next, to the lyophilized sample, 5 µl of acetonitrile (HPLC gradient grade, Sigma Aldrich) and 95 µL of 0.05 M, pH 7 Triethylammonium acetate (TEAA) buffer (Sigma Aldrich) were added and mixed thoroughly. The mixture was equilibrated for 30 min at room temperature and transferred to the HPLC vial. The solution was purified by reverse-phase high performance liquid chromatography (HPLC 1200 Agilent Technologies) with C3 column (Agilent Zorbax 300SB-C3) using a gradient of acetonitrile (concentration was increased from 5% to 50% over 30 min) in 0.05 mM TEAA (pH 7) at a flow rate of 0.4 mL/min as a mobile phase. The unreacted oligonucleotides were eluted at around 11-13 min, and PEG-modified oligonucleotides were eluted at around 20-22 min. The collected fractions were lyophilized and dissolved in 1x PBS for analysis with PAGE gel as mentioned above.

**Validating DBCO activity after purification.** To validate the activity of DBCO, homobifunctional PEG with Azido moiety were used to form dimers. The results were characterized using PAGE gels.

**Adding purified oligo-PEG-DBCO or biotin to DNAO.** The purified conjugates were added to the DNAO at ten times excess over each extension. The mixture was incubated for 1 hour at room temperature. The excess conjugates were removed using 50 kDa Pluronic passivated ultrafiltration columns.

#### Assembly of Iron oxide nanoparticles (IONPs) functionalized structures.

Purified DNAO, DOPMs or silica coated structures carrying 1, 2 or 3 binding sites with biotin groups were mixed with Streptavidin coated IONPs (Purchased from Ocean Nanotech, LLC USA) at three times excess over each binding site. The reaction mixture was incubated overnight at room temperature. The IONPs functionalized structures were immediately used for characterization TEM.

### Supporting Note

#### Supporting Note 1.

**Dynamic light scattering (DLS)** is used for characterizing the size distribution profile of particles in a suspension. The principal of the technique is that the particles are in constant thermal motion, called Brownian motion, and diffusion speed of these particles is a function of their size. Smaller particles diffuse faster than larger particles. To measure the diffusion speeds, the particles are illuminated with a laser and the pattern (scattering intensity) produced is observed using a sensitive avalanche photodiode detector (APD). The change in intensity can be analyzed and the fluctuations produces a curve which is correlated with time to measure size and size distribution. The size of the particles is measured using the Stokes-Einstein equation which gives the hydrodynamic radius (or diameter, $R_d$) of the particle using the diffusion coefficient (D) according to this:

$$R_d = \frac{k_B T}{3\pi\eta D}$$

Where, $k_B$ is the Boltzmann constant, T is the absolute temperature and $\eta$ is the liquid viscosity.

DLS measures the apparent size of the dynamic hydration layer that exists around a solid spherical particle in a solution and gives in the form of **Z-average** ($Z_{avg}$). However, the assumption that particles are spherical, does not reflect the true particle size for non-spherical particles. This suggests that using a highly asymmetric structure such as a rod or a flat sheet would result in multiple peaks. In this paper, we have used the DNA brick design which as per the DLS results is assumed to be a sphere with a diameter of 52 ± 2 nm instead of a cuboid of 55 x 20 x 10 nm dimensions. This was consistent across all the measurements and therefore adapted as control for comparison.

Another important parameter that is measured is the **Derived mean count rate (kcps)**, which is the theoretical count rate one would obtain at 100% laser power with zero attenuation. Collectively, this is the total number of scattered photons detected and is usually stated in a per second basis while accounting for the different attenuation levels. This parameter allowed to observe the addition of inorganic material on the DNA origami as a function of time. The increase in the scattering intensity directly correlated to the increase in the inorganic material on the origami. This parameter allows tracking the material growth when the $Z_{avg}$ stays constant up until the aggregation time point.

### Supporting Note 2.

**Semi-automated size measurements of structures using TEM images.** To quantitatively analyze the size of the structures imaged using transmission electron microscopy (TEM), we used ImageJ to plot the grey value profiles of the structures and then used the python script (https://github.com/ewuirl/TEM-ruler) to analyze the profiles and calculate the average size distribution. Specifically, we imported the TEM images into ImageJ and used the ROI manager to draw regions of interest (ROIs) around the structures of interest. Then, we selected "plot profile" from the ImageJ menu to obtain the grey value profiles for each ROI. Finally, we used the python script to analyze the grey value profiles and calculate the average size distribution of the structures based on the profile data.

### Supporting Figures.

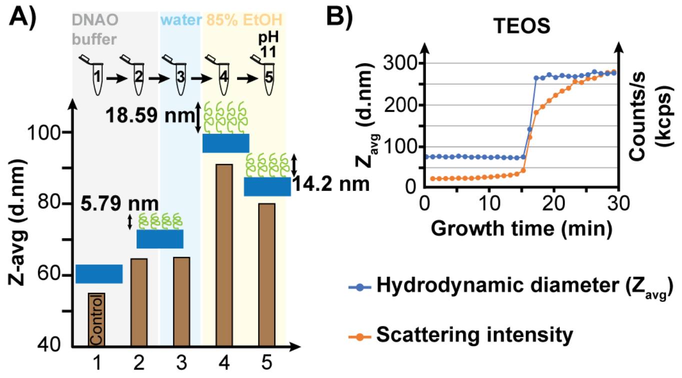

Figure S 1. A) The effect of different buffer conditions on the 42HB DOPMs (tubes 2 to 5; the blue rectangular structures depict 42HBs with the PEG chains in green). Tube 1 represents the 42HB DNAO in native buffer. The structural stability and dispersity across the different solvent conditions (depicted in grey background for native buffer, blue for water and yellow for 85% ethanol) was analyzed using DLS. B) Preliminary test to grow silica shell on 42HB DOPMs in 85% ethanol at pH 11 using tetraethyl orthosilicate (TEOS) as precursor. The silica growth on 42HBs was observed using DLS (Dynamic light scattering) for an incubation period of 30 min. The blue and orange curves represent the measured hydrodynamic diameter (in nm) and scattering intensity curves (in counts/s) as a function of the growth time, respectively.

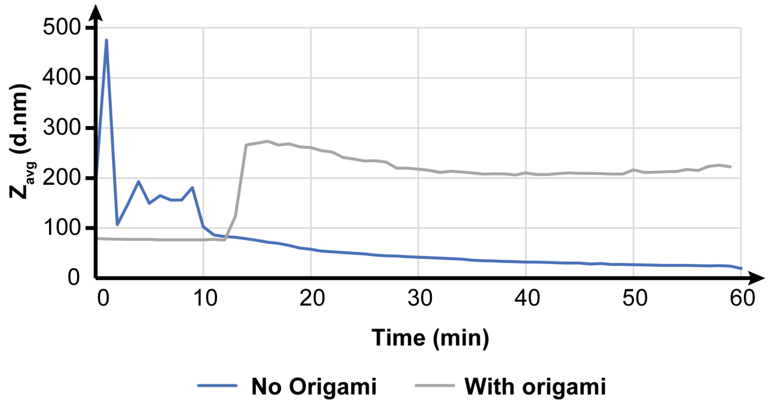

Figure S 2. DLS measurements of silica growth without the presence of DNA origami as a template (blue curve). Changes in the Zavg were plotted as a function of the incubation time. As a comparison the silica growth with DNA origami as a template is overlayed in the graph (grey curve). For this, the growth was conducted in 85% ethanol, pH 11 reaction solution using TEOS as a precursor at 1x concentration. The growth reaction with origami was performed with DNA brick polyplexed with PEG-PLL, PEG length 5 kDa.

From the DLS results, random fluctuations were observed for no origami growth in the first 14 min before attaining a stable, progressively declining Zavg. We hypothesize that these fluctuations arise from random aggregation of the silicates as it undergoes hydrolysis and subsequent condensation and redistribution. In the absence of a template these aggregates undergo rapid changes that are not appropriately averaged by the DLS algorithm. We concluded that without a template these random fluctuations indicate the absence of any significant particle growth, which eventually undergoes steady decrease in the Zavg in the remaining duration of the growth.

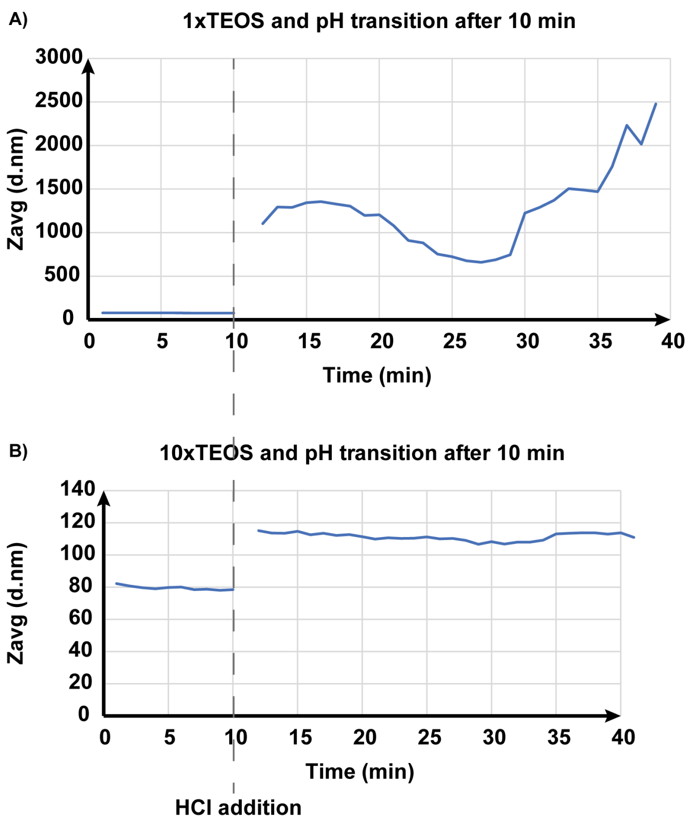

Figure S 3. DLS measurements of pH transition from 11 to 7 using HCl during the silica growth for DOPMs with 5 kDa PEG chain length at A) 1x or B) 10x TEOS concentration. Changes in the Zavg were plotted as a function of the incubation time. For this, the growth was conducted in 85% ethanol, pH 11 reaction solution. The growth reaction was performed with DNA brick polyplexed with PEG-PLL, PEG length 5 kDa.

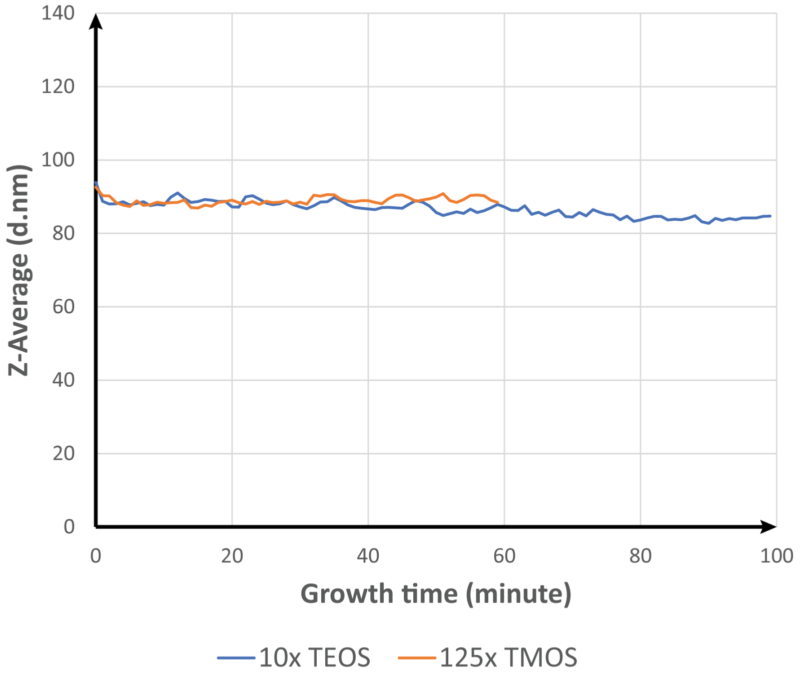

Figure S 4. DLS measurements of silica growth in methanol for DOPMs with 5 kDa PEG chain length using either TEOS or TMOS as precursor (blue curve for 10x TEOS and orange curve for 125x TMOS concentration). Changes in the Zavg were plotted as a function of the incubation time. For this, the growth was conducted in 85% methanol, pH 11 reaction solution. The growth reaction was performed with DNA brick polyplexed with PEG-PLL, PEG length 5 kDa.

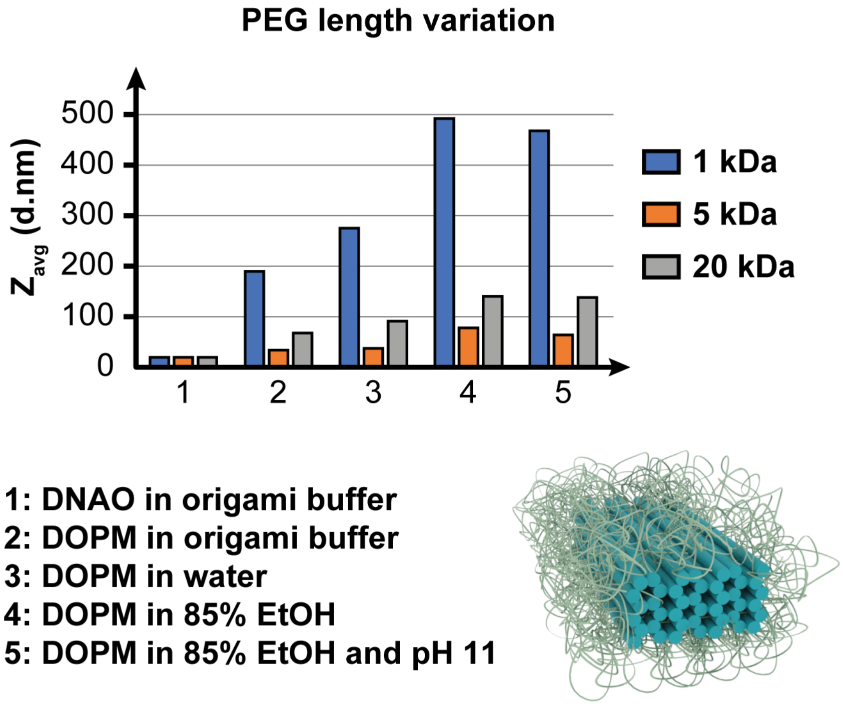

Figure S 5. DLS measurements of the DOPMs with varying lengths of the PEG chain in different buffer conditions. For this, three variations of PEG-PLL were used with a constant cationic segment of 10 lysine repeats while using either of the lengths for the PEG chain: 1 (Blue), 5 (Orange) or 20 (Grey) kDa.

For 1 kDa PEG length, the condition 2 with standing origami buffer (1x TE + 12 mM $MgCl_2$), was observed to aggregate formation, that upon further changes in the buffer condition led to higher-order aggregates. From this we suspect that 1 kDa PEG lengths are not sufficiently long to avoid aggregation. Due to this observation, 1 kDa PEG lengths were not further investigated.

For 5 and 20 kDa PEG lengths, the Zavg average increased to a maximum for 85% ethanol buffer conditions (15% ultrapure water) with ~90 nm for 5 kDa and ~129 nm for 20 kDa PEG lengths. Further changes to pH 11 only had a slight decrease in the Zavg for either of the PEG lengths.

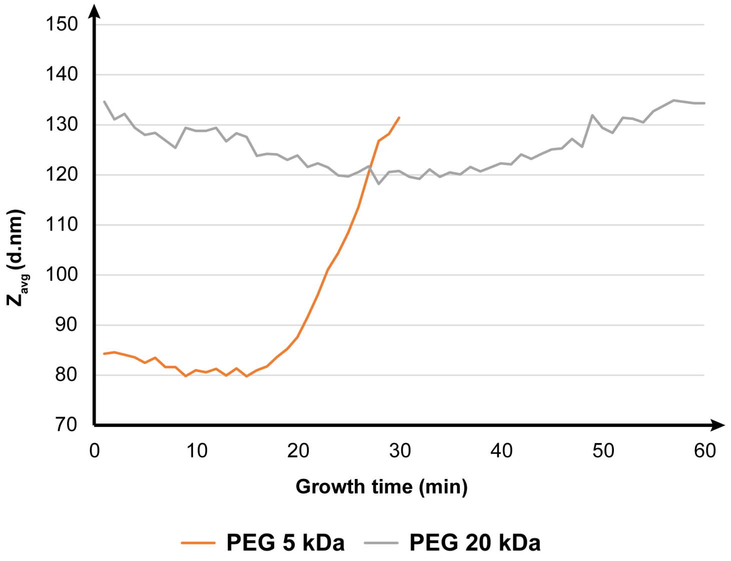

Figure S 6. DLS measurements of silica growth for DOPMs with varying PEG lengths (orange curve for 5 kDa and grey curve for 20 kDa PEG length). Changes in the Zavg were plotted as a function of the incubation time. For this, the growth was conducted in 85% ethanol, pH 11 reaction solution using TMOS as a precursor at 125x concentration. The growth reaction was performed with DNA brick.

A)

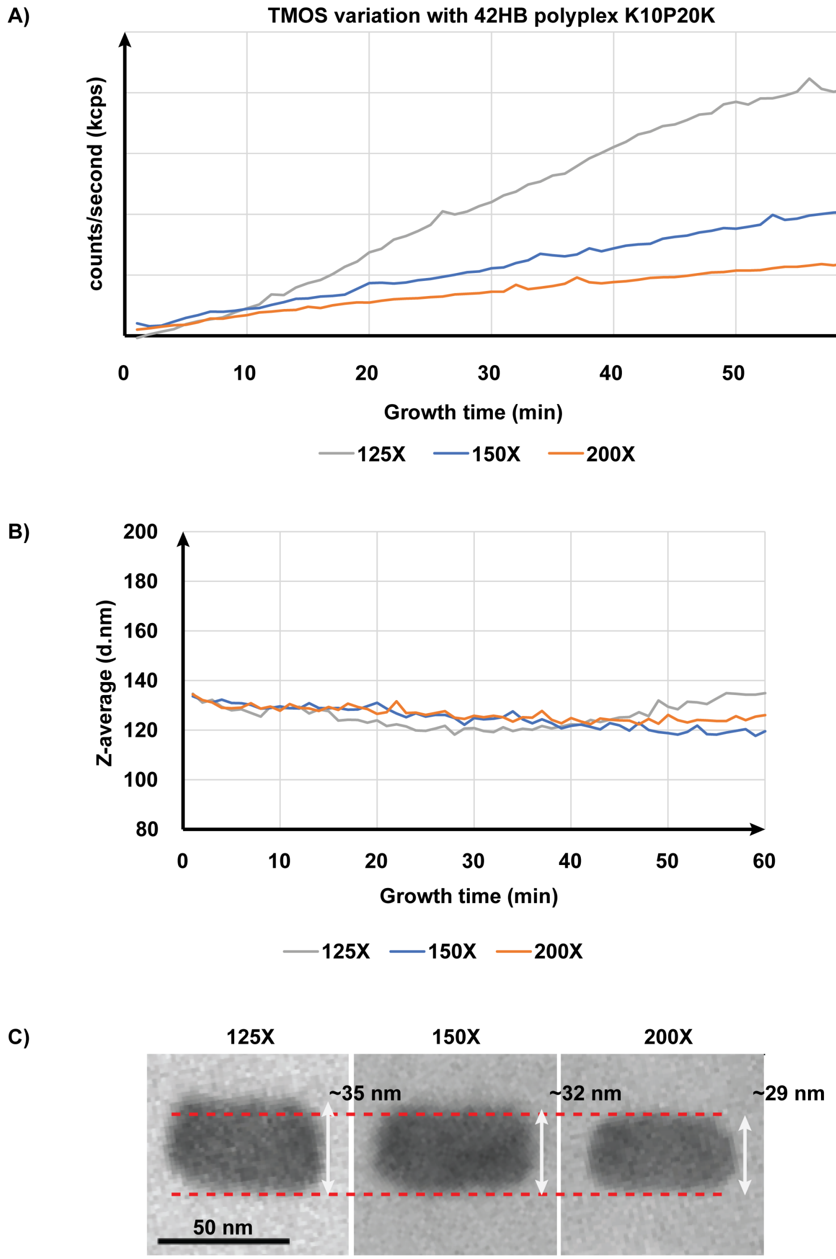

Figure S 7. DLS measurements of silica growth for DOPMs with 20 kDa PEG chain length at varying TMOS concentration (grey curve for 125x, blue curve for 150x and orange curve for 200x TMOS concentration). A) Changes in the derived mean count rate (counts/second) were plotted as a function of incubation time along with B) Changes in the Zavg were plotted as a function of the incubation time. For this, the growth was conducted in 85% ethanol, pH 11 reaction solution using TMOS as a precursor at 125x concentration. The growth reaction was performed with DNA brick polyplexed with PEG-PLL, PEG length 20 kDa. C) Corresponding TEM images. Scale bars = 50 nm.

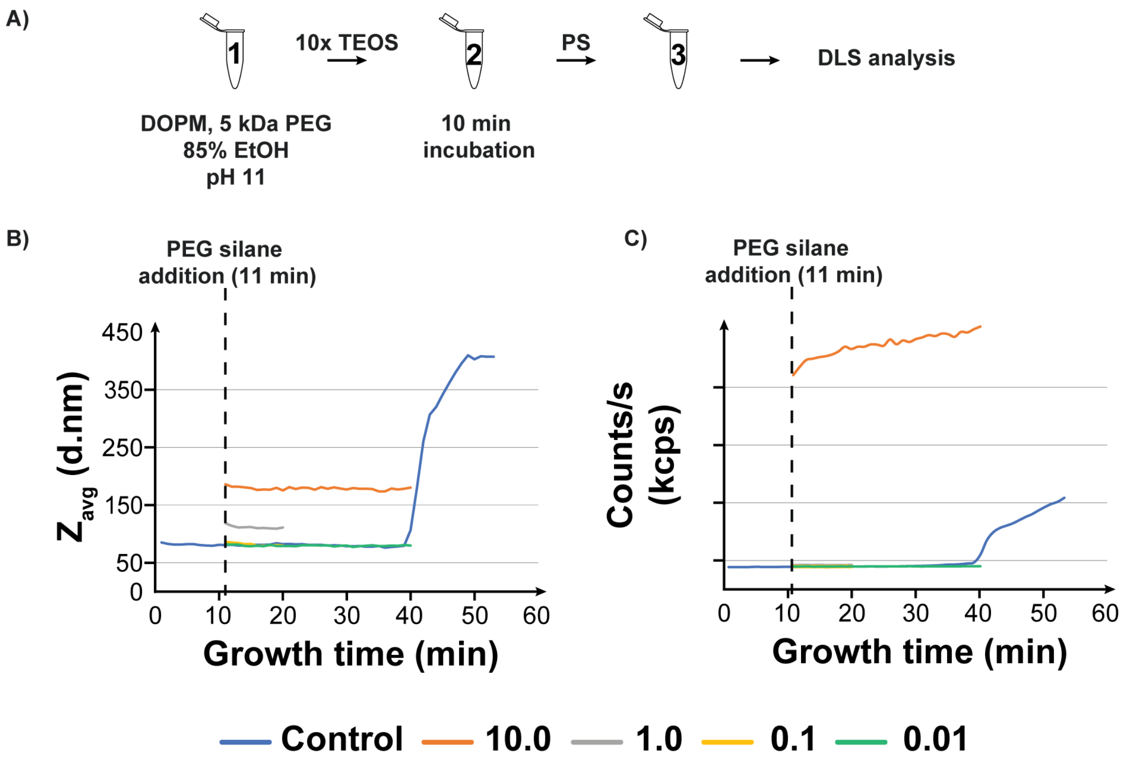

Figure S 8. DLS data for silica shell growth on 42HB DOPMs with 5 kDa PEG lengths using a 10x concentration of TEOS as the precursor. The silica shell growth in these set of reactions were terminated by the addition of PEG-Silane (PS). The PS was added to the reaction after 10 min growth. The addition steps are outlined in (A). PS concentration was varied relative to the precursor concentration. B) and C) $Z_{avg}$ and scattering plots respectively.

In case of addition of 10:1 PS concentration and 1:1 PS concentration we observed a significant increase in the Zavg when compared to the control (178.8 ± 2.56 nm in case of 10:1 and 111.9 ± 2.62 nm in case of 1:1). However, for both 0.1:1 and 0.01:1 PS concentrations, no such increase in the Zavg we observed. We hypothesize that, like TMOS, PS undergo rapid hydrolysis as well as condensation, like what was observed in case of 10x TMOS drive silica shell growth (Figure 2. DNA brick origami structures are subjected to various optimized conditions that impact the silica shell growth to obtain fine-tuned control over the growth process. A-D) DLS data for the optimization tests for solvent pH conditions. E-H) DLS data for alcohol-to-water content optimizations. I-L) DLS data for optimizing the choice of precursor (molecular structure of the tested precursors in I) and precursor concentrations. M-N) DLS and TEM data for PEG-length optimization studies. Scale bars = 100 nm.-I). The higher PS concentrations (equivalent to 12.5x TMOS) led to an increased uncontrolled rate of deposition, adding to the growing silica shell, thus increasing the thickness in addition to the reaction termination. Reduction of the PS molar concentration led to a controlled rate of deposition, as observed from 0.1:1 and 0.01:1 PS variations.

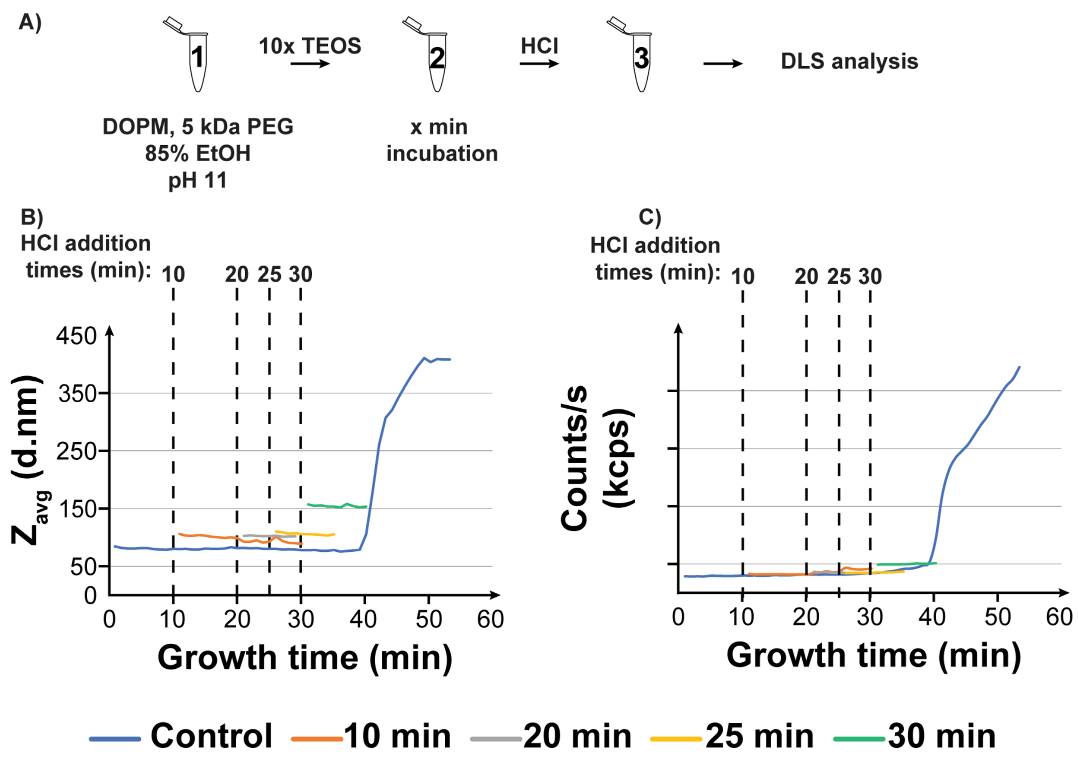

Figure S 9. DLS data for silica shell growth on 42HB DOPMs with 5 kDa PEG lengths using a 10x concentration of TEOS as the precursor. The silica shell growth in these set of reactions were terminated by the addition of HCl, enforcing the change of pH of the reaction mixture from pH 11 to 7. HCl was added at different time points to assess its effect. The steps are outlined in (A), whole B) and C) represents the $Z_{avg}$ and scattering plots respectively.

The largest $Z_{avg}$ of 154.9 ± 2 nm was observed for the 30 min HCl addition, while the smallest observed Zavg was 100.2 ± 1.4 nm for the 10 min HCl addition. In addition, we also observed that the addition of HCl at (or after) 40 min led to the aggregation of the structures (data not shown).

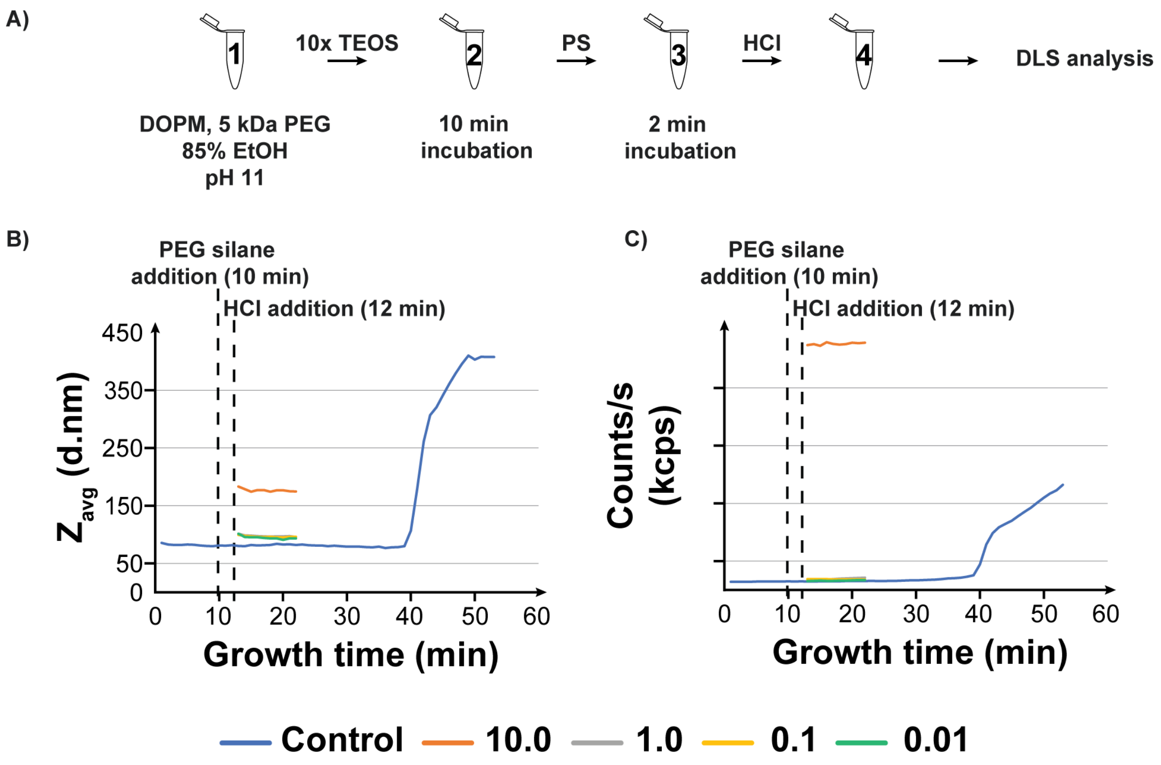

Figure S 10. DLS data for silica shell growth on 42HB DOPMs with 5 kDa PEG lengths using a 10x concentration of TEOS as the precursor. The silica shell growth in these set of reactions were terminated by the addition of PS and HCl. PS was added after 10 min incubation and followed the addition of HCl 2 min later. PS concentration was varied relative to the precursor concentration. The steps are outlined in (A), whole B) and C) represents the $Z_{avg}$ and scattering plots respectively.

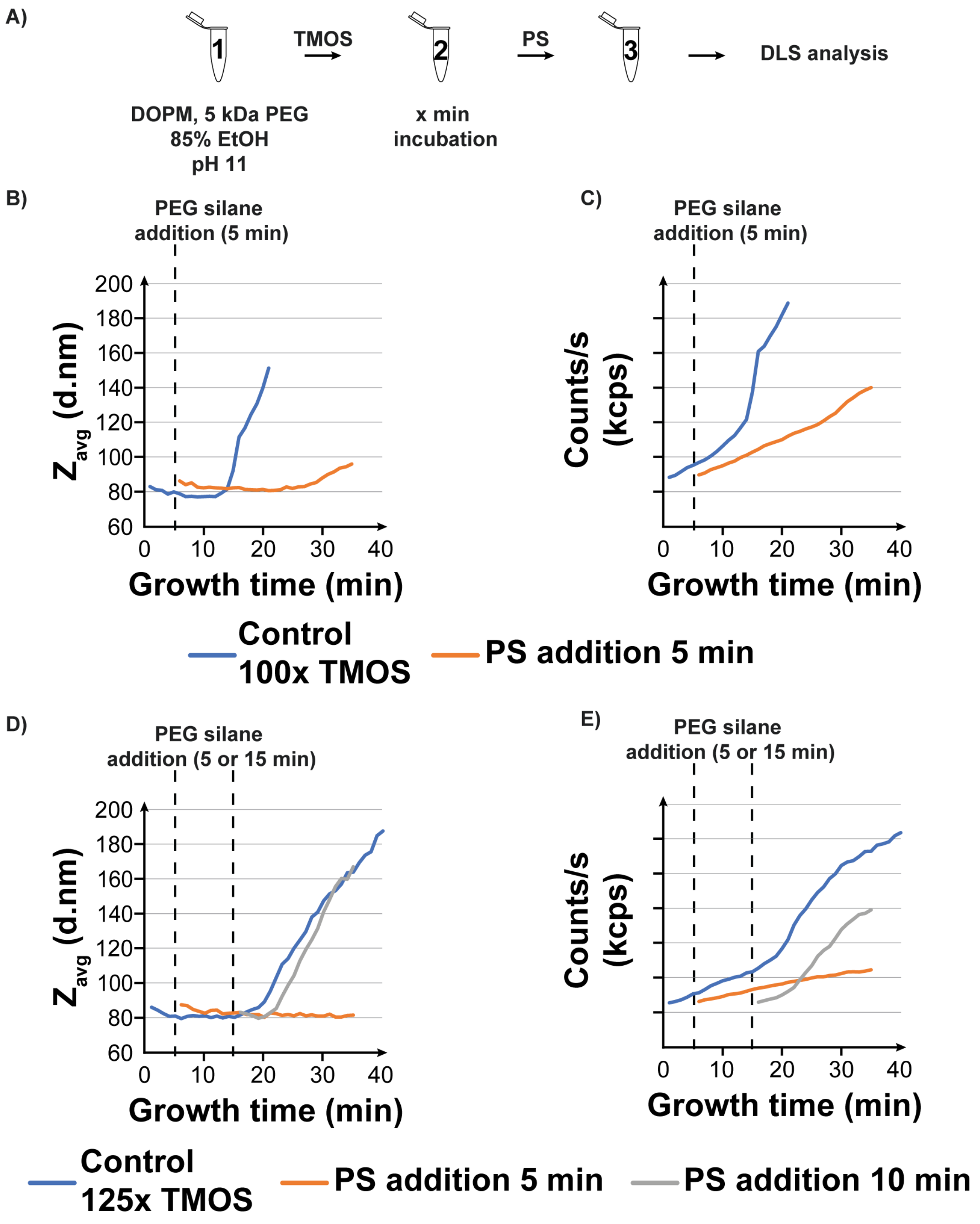

Figure S 11. DLS data for silica shell growth on 42HB DOPMs with 5 kDa PEG lengths using a 100x and 125x concentrations of TMOS as the precursor. The silica shell growth in these set of reactions were terminated by the addition of PEG-Silane (PS). The PS was added to the reaction after 5 or 15 min growth. The addition steps are outlined in (A). PS concentration was varied relative to the precursor concentration. B), D) and C, E) $Z_{avg}$ and scattering plots respectively.

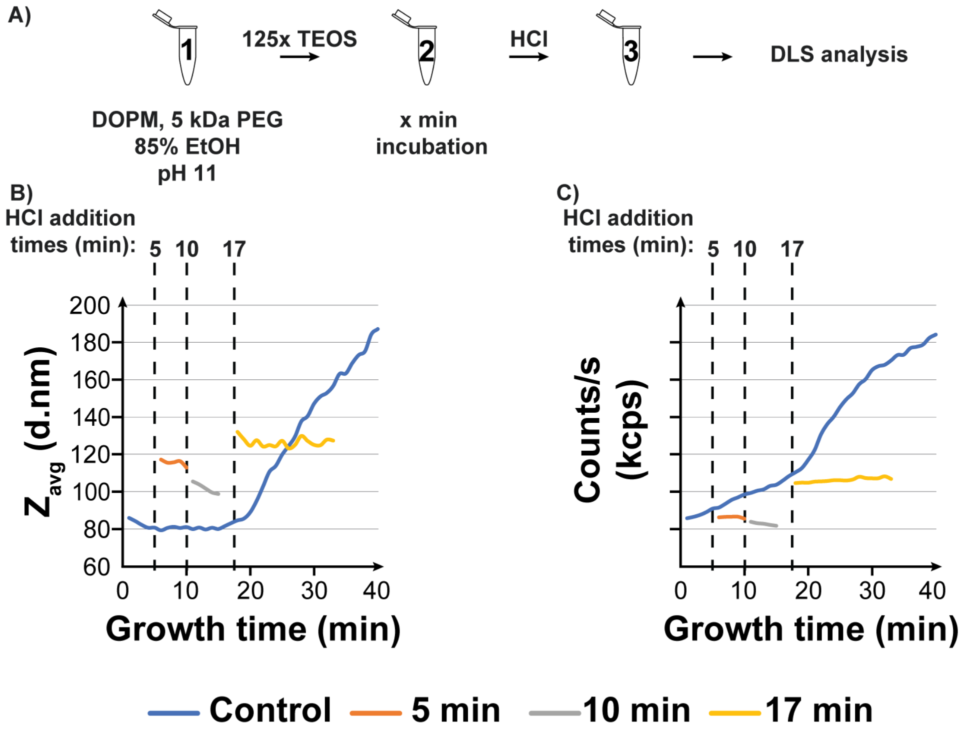

Figure S 12. DLS data for silica shell growth on 42HB DOPMs with 5 kDa PEG lengths using a 125x concentration of TMOS as the precursor. The silica shell growth in these set of reactions were terminated by the addition of HCl, enforcing the change of pH of the reaction mixture from pH 11 to 7. HCl was added at different time points to assess its effect. The steps are outlined in (A), whole B) and C) represents the $Z_{avg}$ and scattering plots respectively.

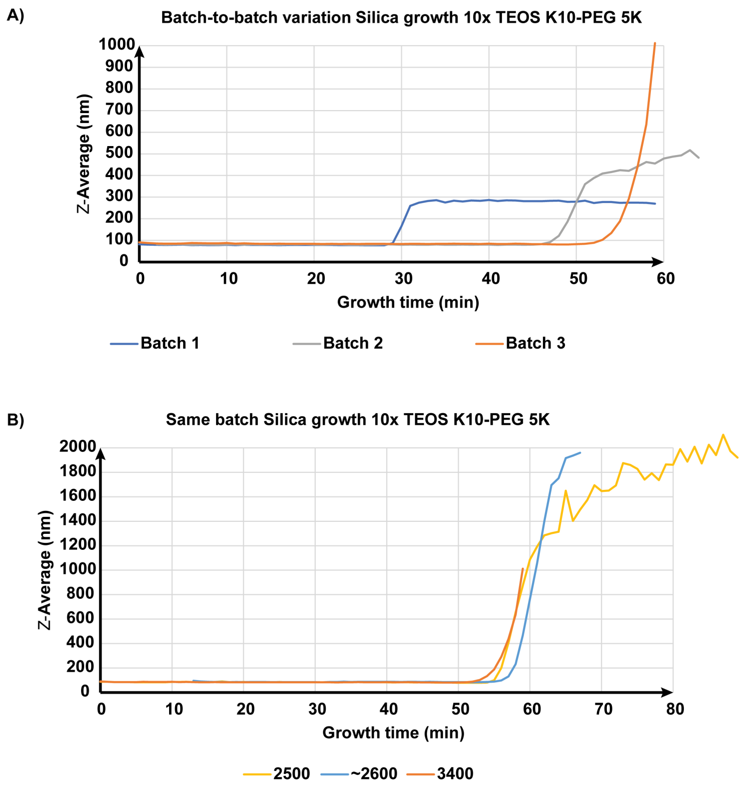

Figure S 13. DLS measurements to understand the batch-to-batch impact on the aggregation time point. Changes in the Zavg were plotted as a function of the incubation time. For this, the growth was conducted in 85% ethanol, pH 11 reaction solution and using TEOS as the precursor. The growth reaction was performed with DNA brick polyplexed with PEG-PLL, PEG length 5 kDa. A) Batch-to-batch variations were compared with B) Same batch variations.

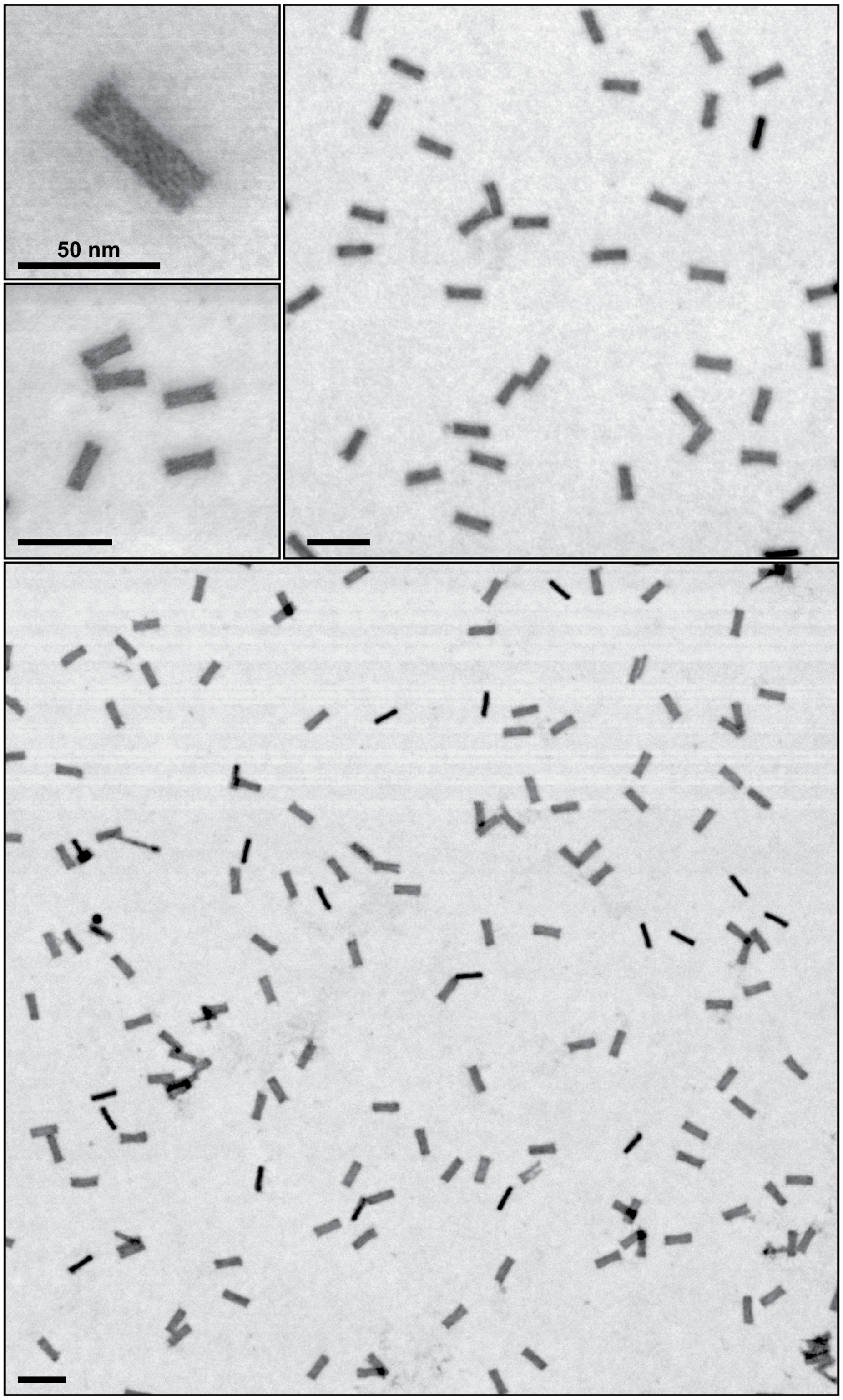

Figure S 14. Electron micrographs representing control DNA brick structures. Scale bars = 100 nm (unless specified otherwise). For these structures, the TEM grids were stained with uranyl formate to facilitate superior contrast to observe the structures.

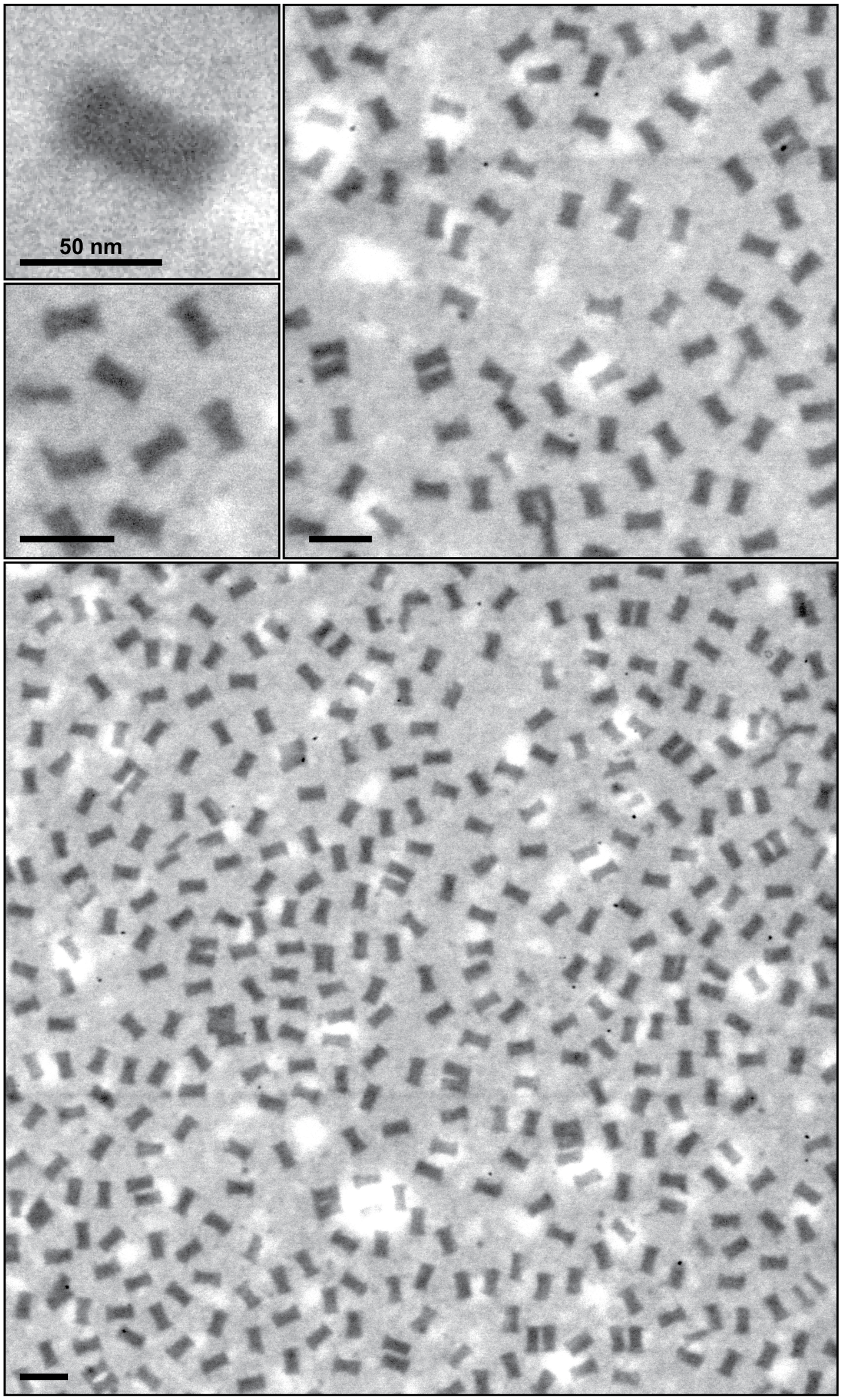

Figure S 15. Electron micrographs representing polyplexed DNA brick structures. For micellization, the K10P20k block co-polymers were used. Scale bars = 100 nm (unless specified otherwise). For these structures, the TEM grids were not stained with uranyl formate.

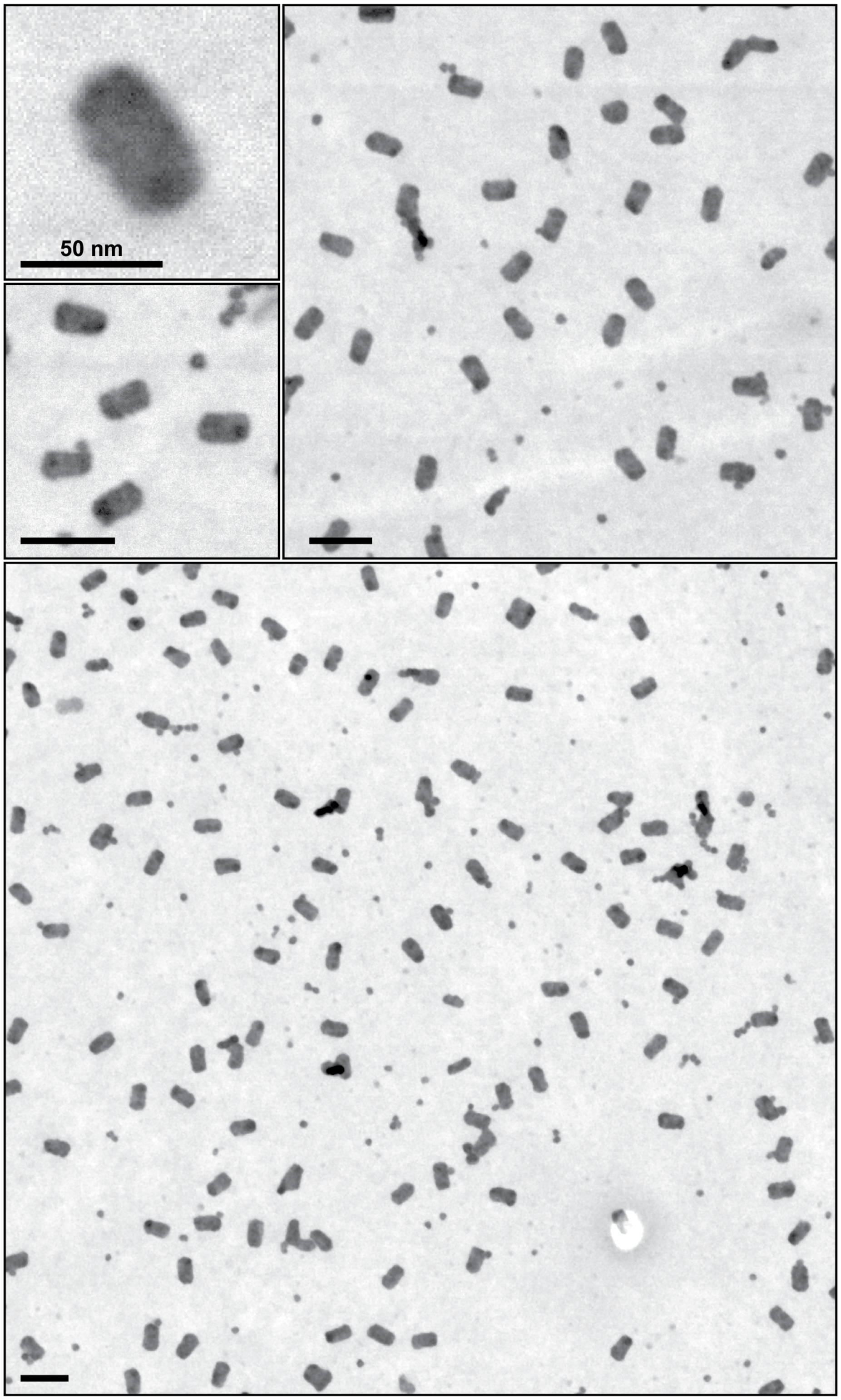

Figure S 16. Electron micrographs representing silica coated DNA brick structures. For micellization, the K10P20k block co-polymers were used. For silica shell growth 125x concentration of TMOS was used. Scale bars = 100 nm (unless specified otherwise). For these structures, the TEM grids were not stained with uranyl formate.

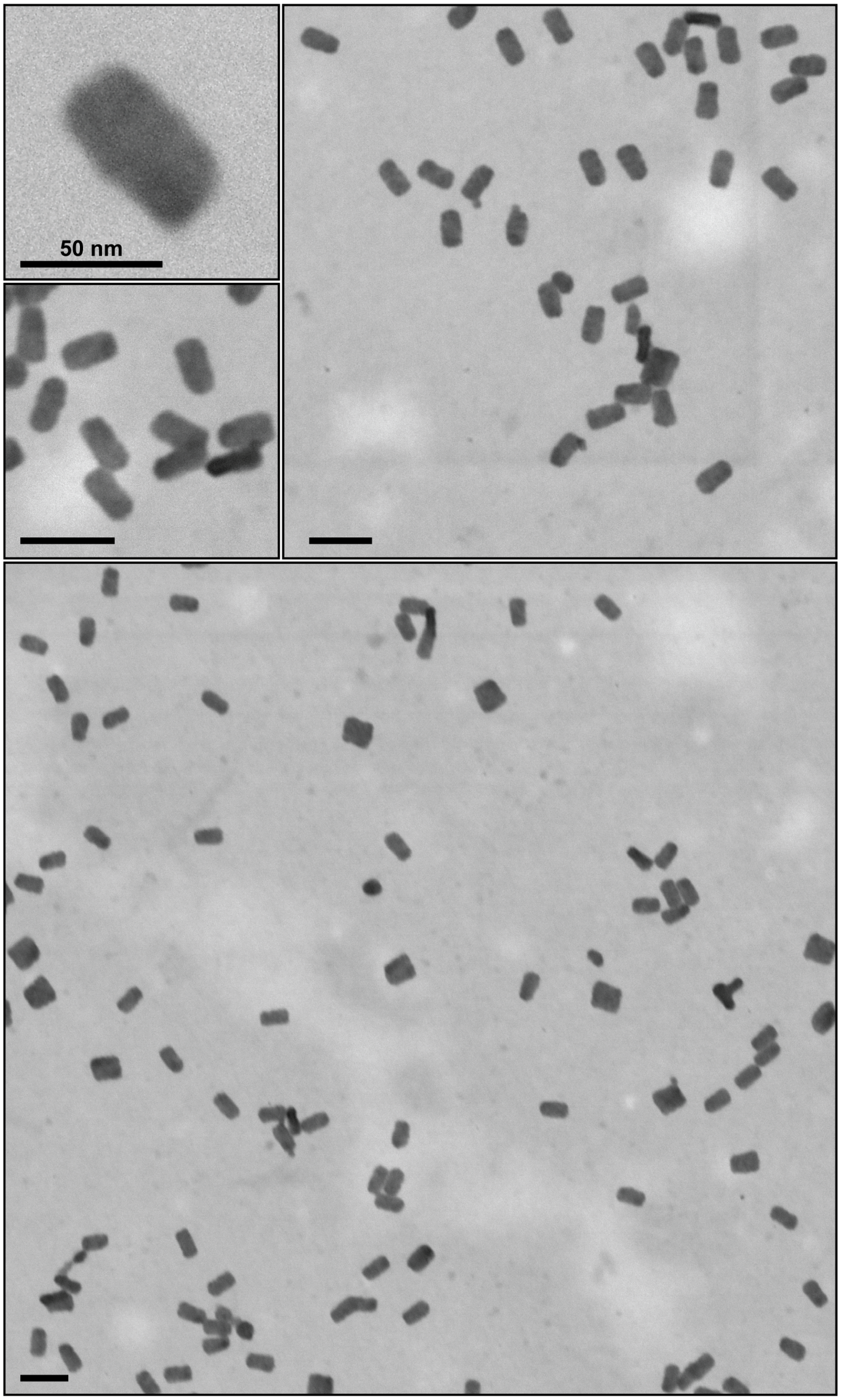

Figure S 17. Electron micrographs representing silica coated DNA brick structures. For micellization, the K10P20k block co-polymers were used. For silica shell growth 150x concentration of TMOS was used. Scale bars = 100 nm (unless specified otherwise). For these structures, the TEM grids were not stained with uranyl formate.

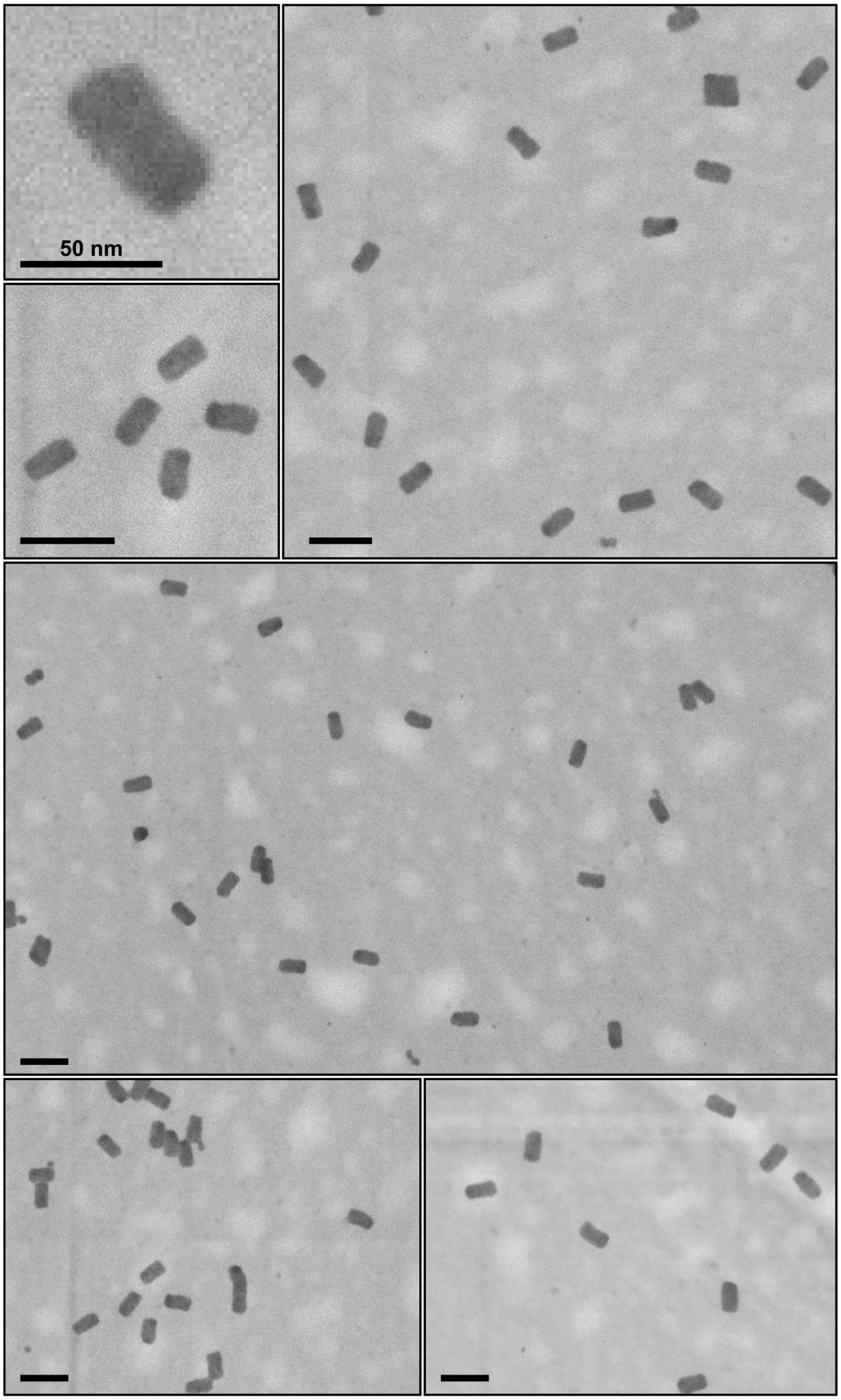

Figure S 18 Electron micrographs representing silica coated DNA brick structures. For micellization, the K10P20k block co-polymers were used. For silica shell growth 200x concentration of TMOS was used. Scale bars = 100 nm (unless specified otherwise). For these structures, the TEM grids were not stained with uranyl formate.

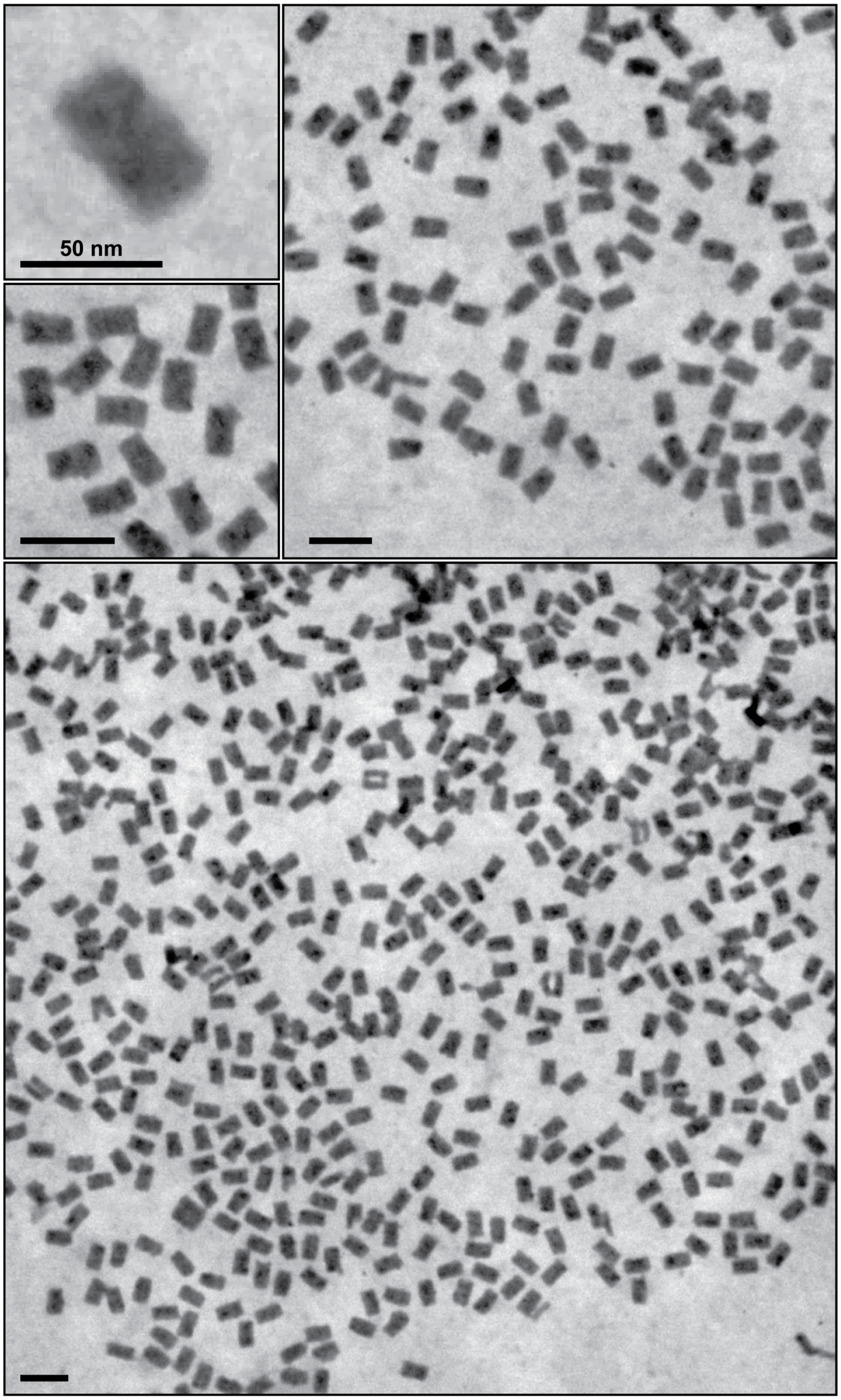

Figure S 19. Electron micrographs of silica coated DNA brick structures. For micellization, the K10P20k block co-polymers were used. For silica shell growth 125x concentration of TMOS was used. The reaction was stopped using PEG-silane followed by the pH transition using HCl. Scale bars = 100 nm (unless specified otherwise). For these structures, the TEM grids were not stained with uranyl formate.

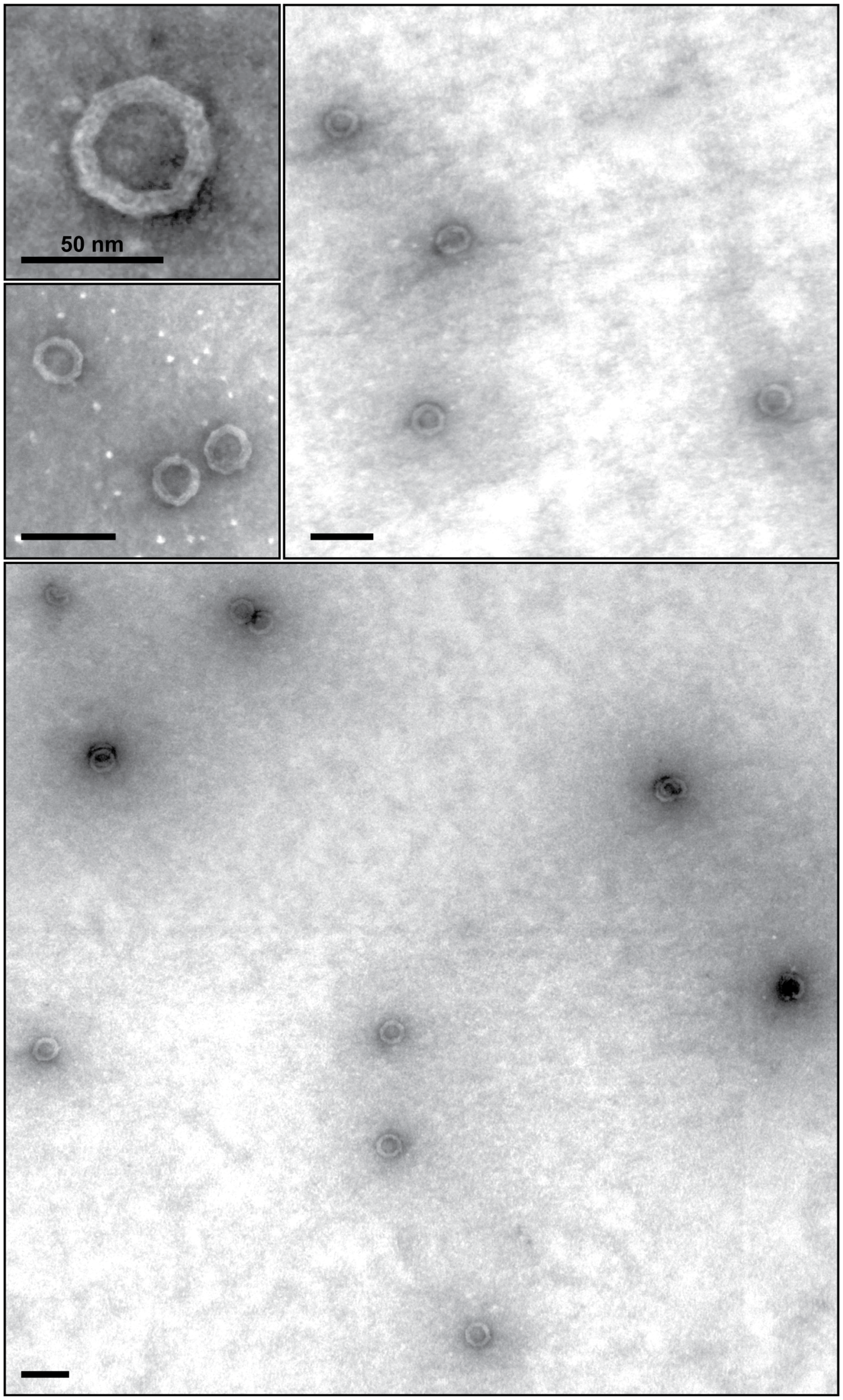

Figure S 20. Electron micrographs representing control DNA ring structures. Scale bars = 100 nm (unless specified otherwise). For these structures, the TEM grids were stained with uranyl formate to facilitate superior contrast to observe the structures.

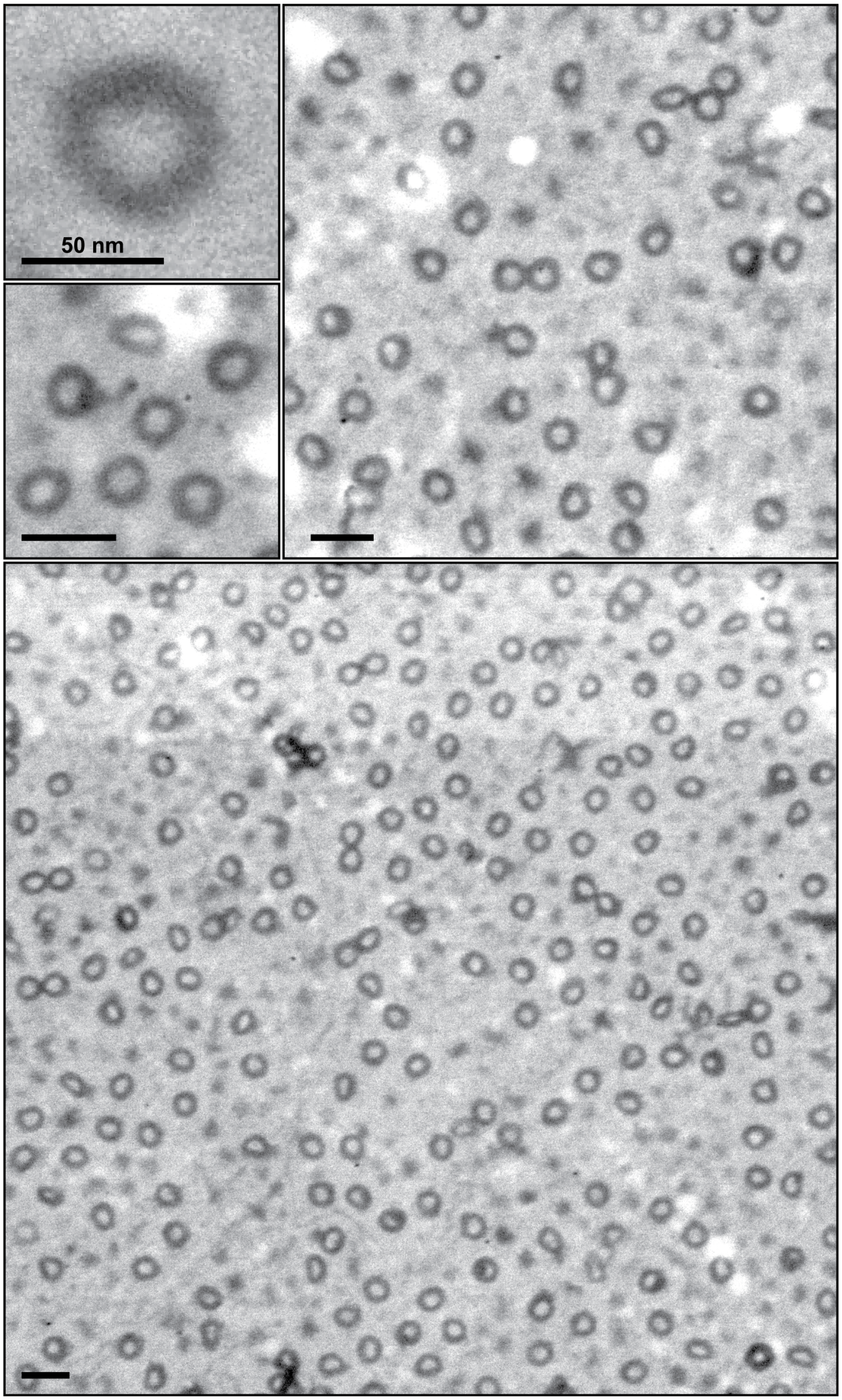

Figure S 21. Electron micrographs representing polyplexed DNA ring structures. For micellization, the K10P20k block co-polymers were used. Scale bars = 100 nm (unless specified otherwise). For these structures, the TEM grids were not stained with uranyl formate.

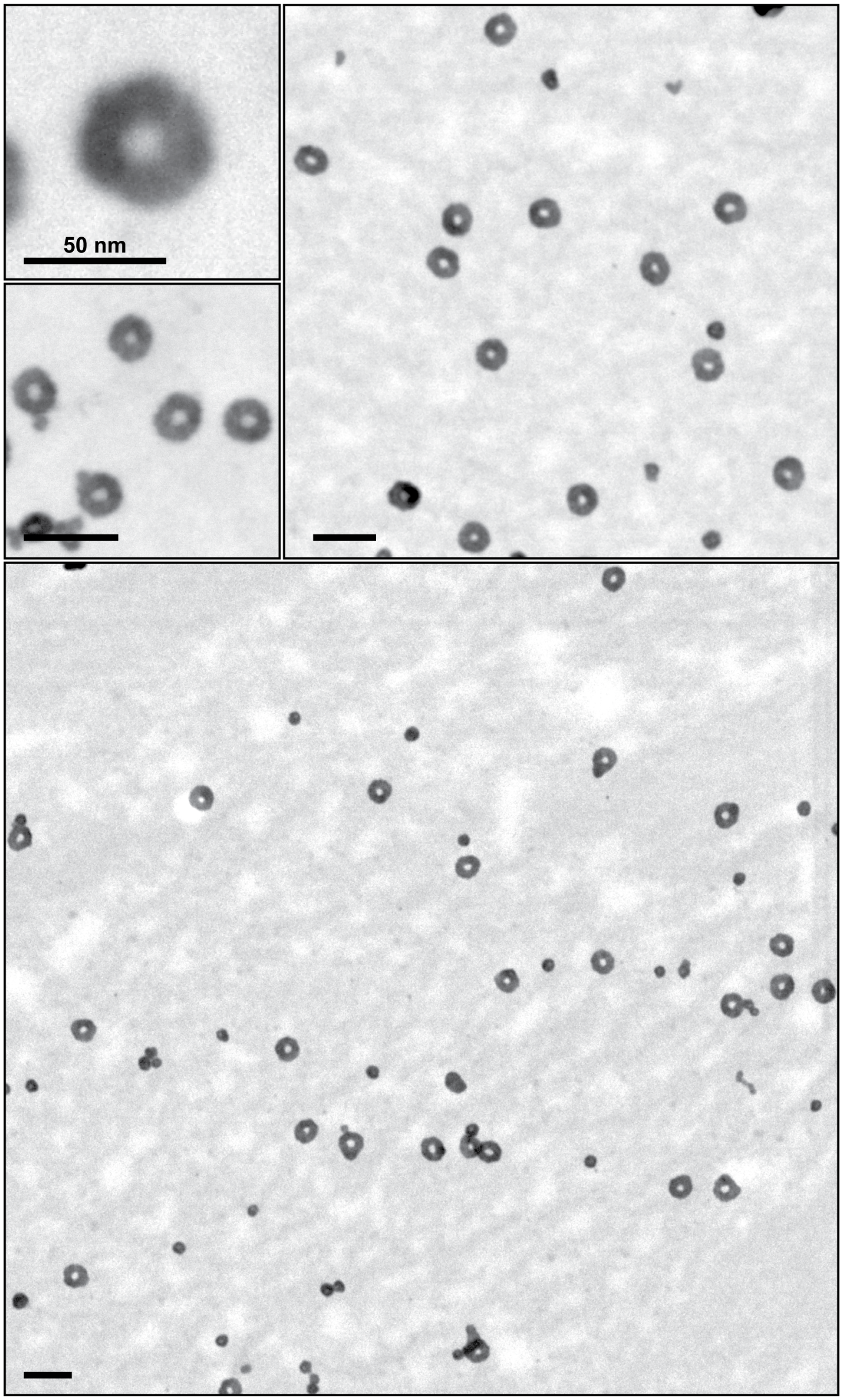

Figure S 22. Electron micrographs representing silica coated DNA ring structures. For micellization, the K10P20k block co-polymers were used. Scale bars = 100 nm (unless specified otherwise). For these structures, the TEM grids were not stained with uranyl formate.

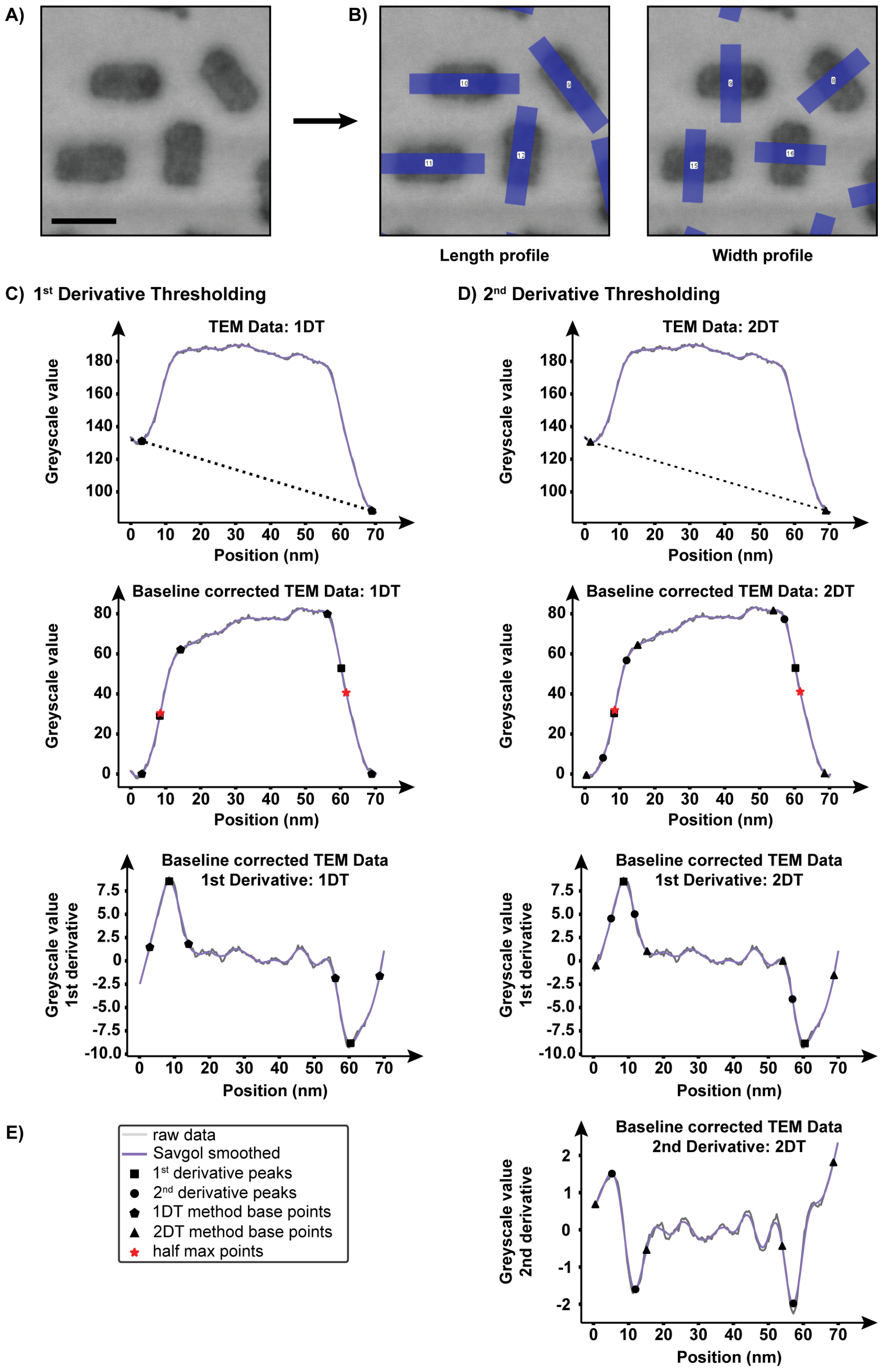

Figure S 23. Example of size measurements of silica coated 42HBs using a semi-automated method on TEM images. A) A small, selected region on the TEM image of silica coated 42HBs. B) Length and width profile (thick blue lines with numbers) around the structures of interest. C and D) 1st and 2nd derivative thresholding with baseline correction, respectively. E) Legend. The scale for all the TEM images = 50 nm.

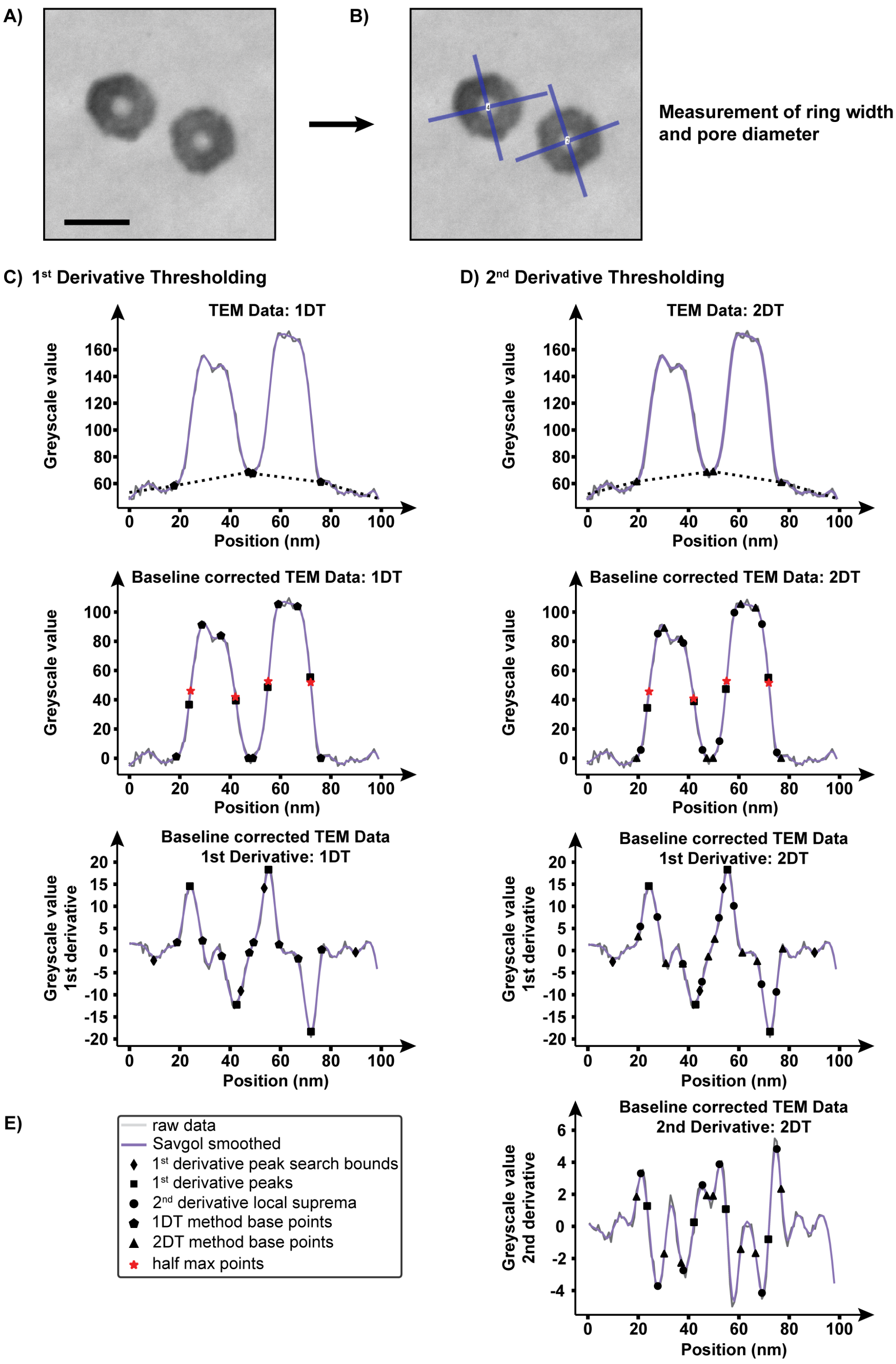

Figure S 24. Example of size measurements of silica coated rings using a semi-automated method on TEM images. A) A small, selected region on the TEM image of silica coated 42HBs. B) Length and width profile (thick blue lines with numbers) around the structures of interest. C and D) 1st and 2nd derivative thresholding with baseline correction, respectively. E) Legend. The scale for all the TEM images = 50 nm.

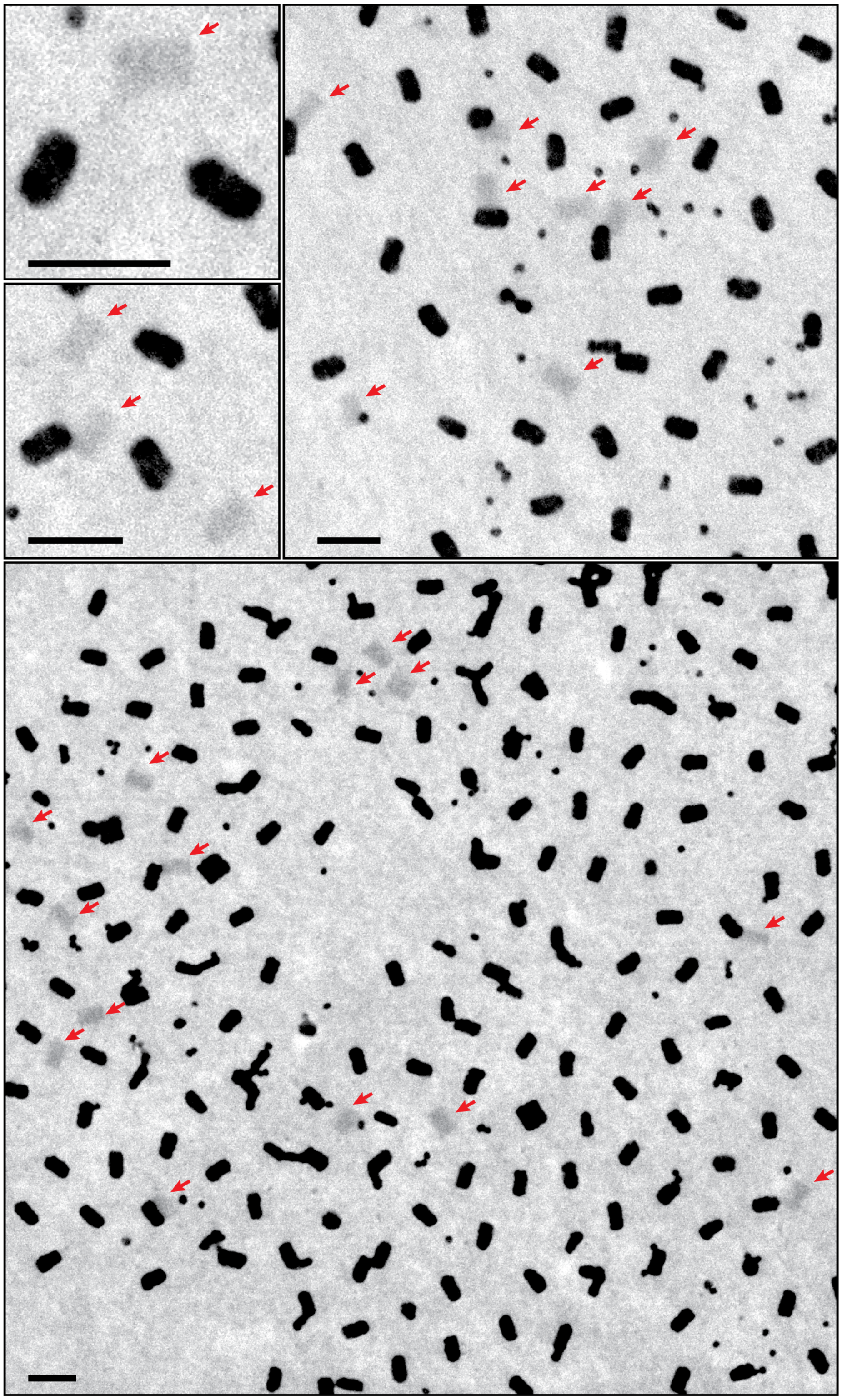

Figure S 25. Electron micrographs representing silica coated DNA brick structures in comparison to the control non-polyplexed, non-silica coated structures (red arrows). For micellization, the K10P20k block co-polymers were used. Scale bars = 100 nm (unless specified otherwise). For this, the TEM grids were not stained with uranyl formate.

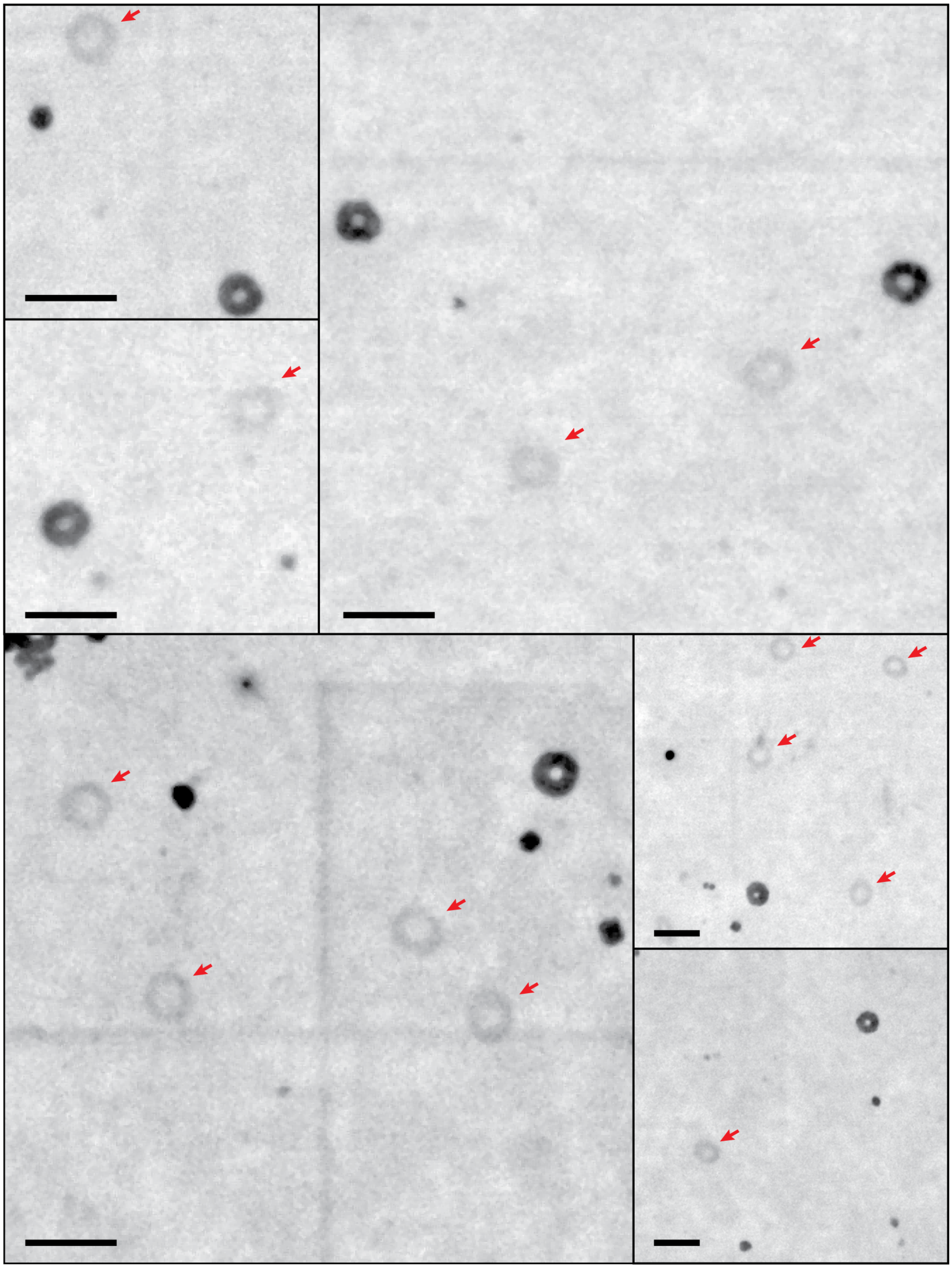

Figure S 26. Electron micrographs representing silica coated DNA ring structures in comparison to the control non-polyplexed, non-silica coated structures (red arrows). For micellization, the K10P20k block co-polymers were used. Scale bars = 100 nm (unless specified otherwise). For this, the TEM grids were not stained with uranyl formate.

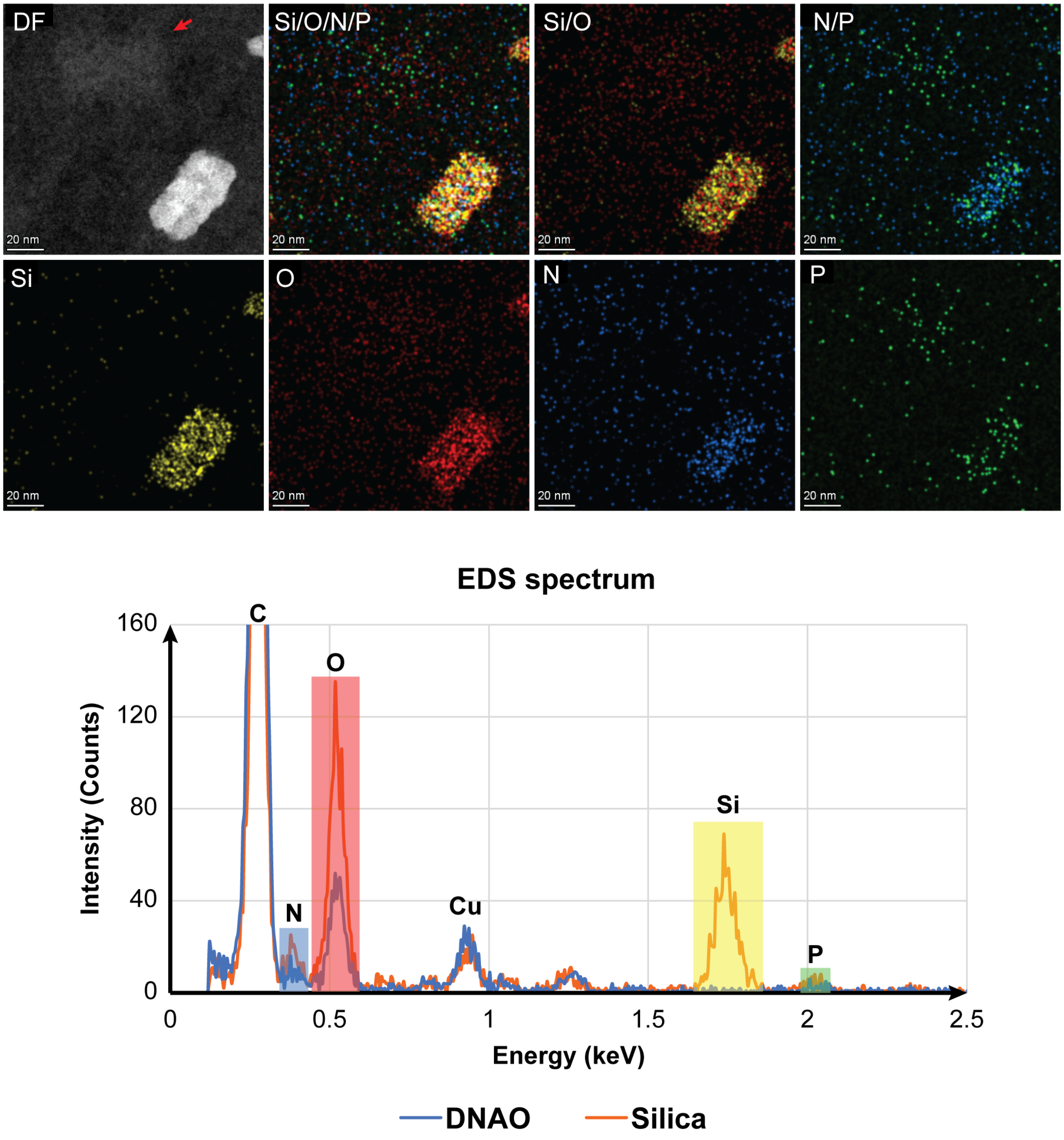

Figure S 27. Energy-Dispersive Spectroscopy (EDS) elemental analysis comparing the non-coated control DNA structures with the silica coated ones. The images represent a collection of electron micrographs overlayed with elemental signal maps as mentioned in the top left of each image (Silica, Si = yellow; Oxygen, O = red; Nitrogen, N = blue; Phosphorus, P = green). Scale bars = 20 nm. In the bottom of the figure, the energy-dispersive spectra against intensity, which has been correlated with the respective elements for both DNAO (blue curve) and silica coated structures (red curve).

A)

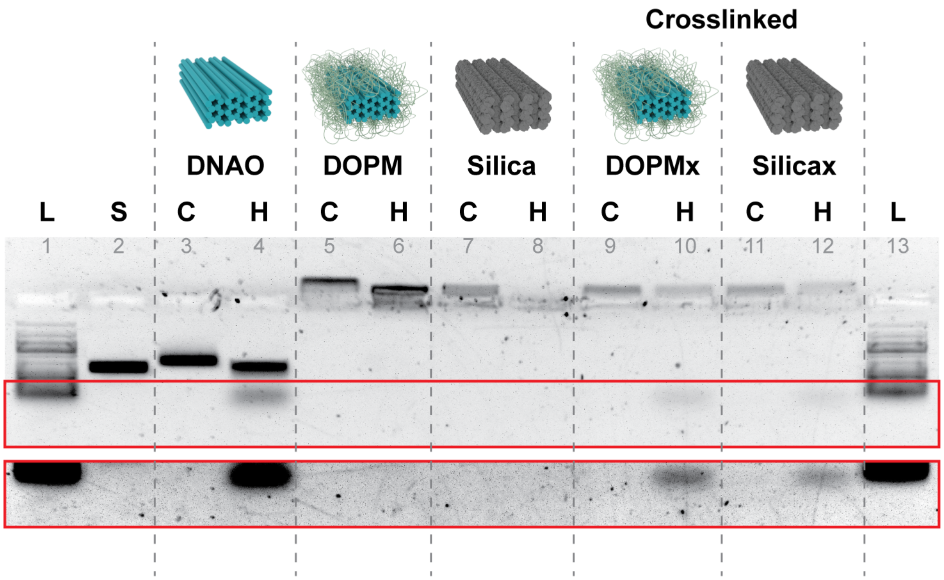

B)

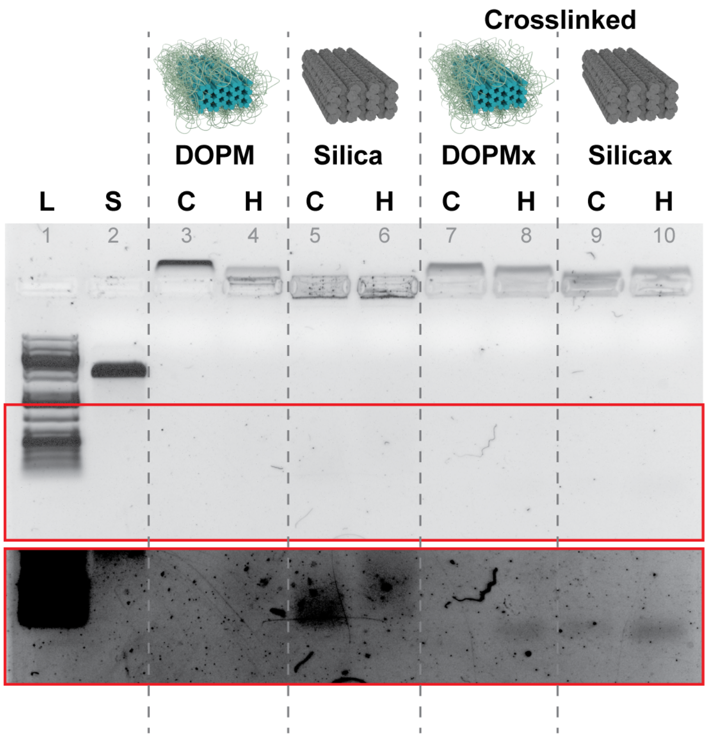

Figure S 28. Agarose gel electrophoresis of the studies with thermal stability of structures.

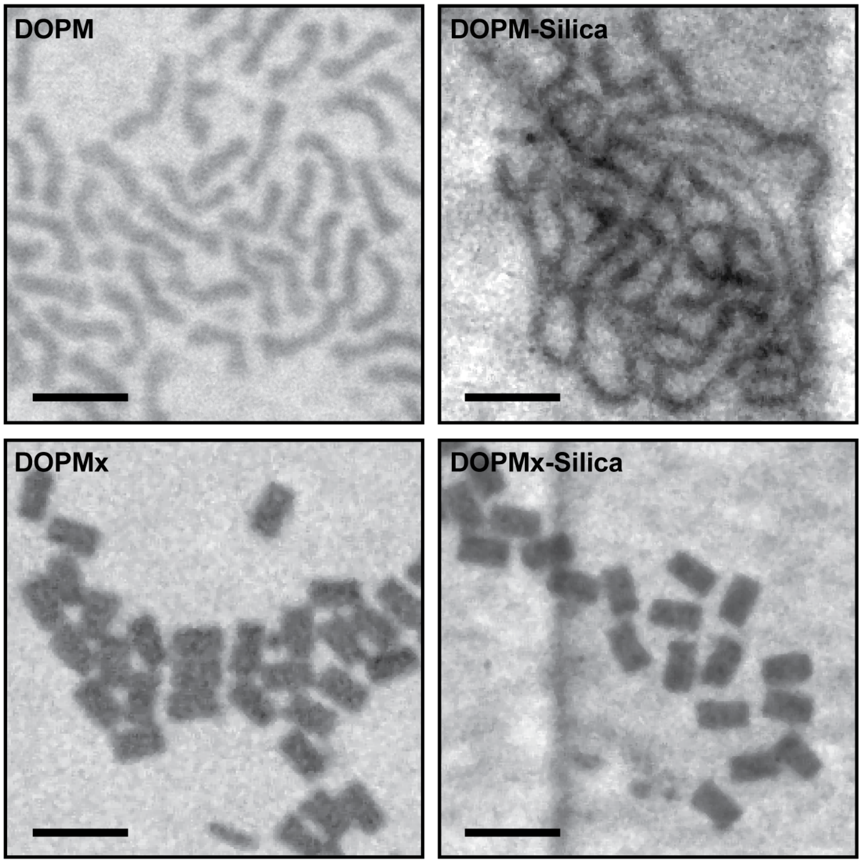

Figure S 29. Evaluating the thermostability of silica-coated structures. Scale = 100 nm. As a control all the variations of the structures involved in the steps leading up to the silica growth were also assessed for thermostability. From the initial tests it was observed that the structures despite of not losing any staples (Figure S 30) lost its structural integrity. From the electron micrographs it can be observed that the structures are no longer brick-shaped and are instead noodle-like. Such observations were made for silica coated structures as well, suggesting that the silica coating does not prevent structural deformation. Our hypothesis is that the electrostatic dynamics of the structure is controlled by the polylysine block in the polymer, which bridge the phosphates in the DNA backbone that maintains the structural integrity. Upon thermal treatment the DNA hybridization is disturbed and the compression forces from the lysines force the structure into a noodle-like formation. However, in the process no staples are lost as they are still strongly held by the electrostatic forces to the polylysine. Such a compression force is also observed for flat-sheet DNA origami structures where the micellization process forces the structure to fold in half.[3]

To reduce such structure-deforming compression forces from the block copolymer, we utilized glutaraldehyde crosslinking established by the Shih group.[4] Glutaraldehyde crosslinks the amines from the polylysines to not only create stronger internal networks that hold the entire structure in place but also reduce the compression forces by reducing the overall positive charge that come from the polymers. From the electron micrographs it can be observed that the structures resemble the brick-shape. However, from the AGE it can be observed that there were some staple leakages. This further proves the previously mentioned hypothesis and that with crosslinking the polylysines the overall charge of the block copolymers is reduced. To avoid the staple leakages, we modified the design to remove all the weakly-bound staples.

From this we can conclude that it is possible to make the structures thermostable to enable applications where DNA origami is typically degraded.

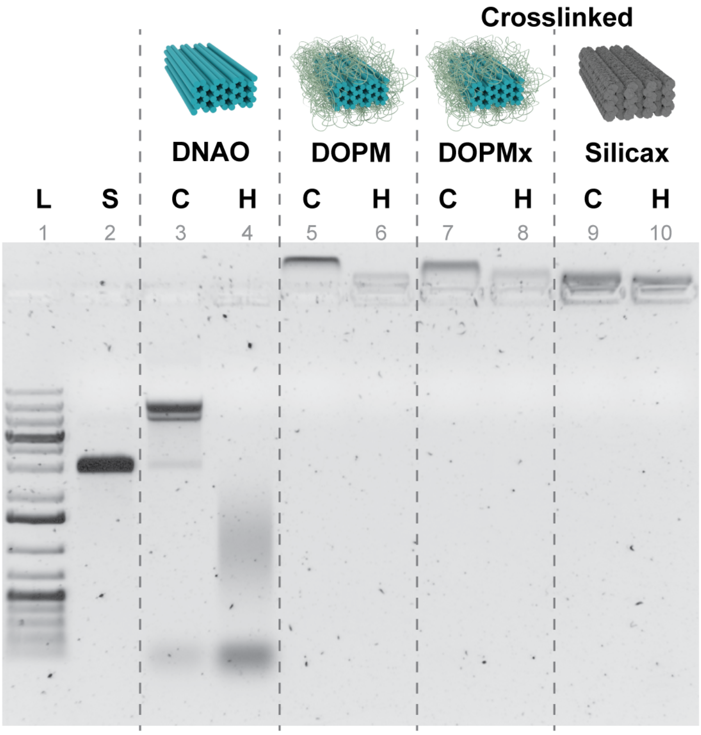

Figure S 30. AGE analysis to confirm the thermostability of silica-coated structures.

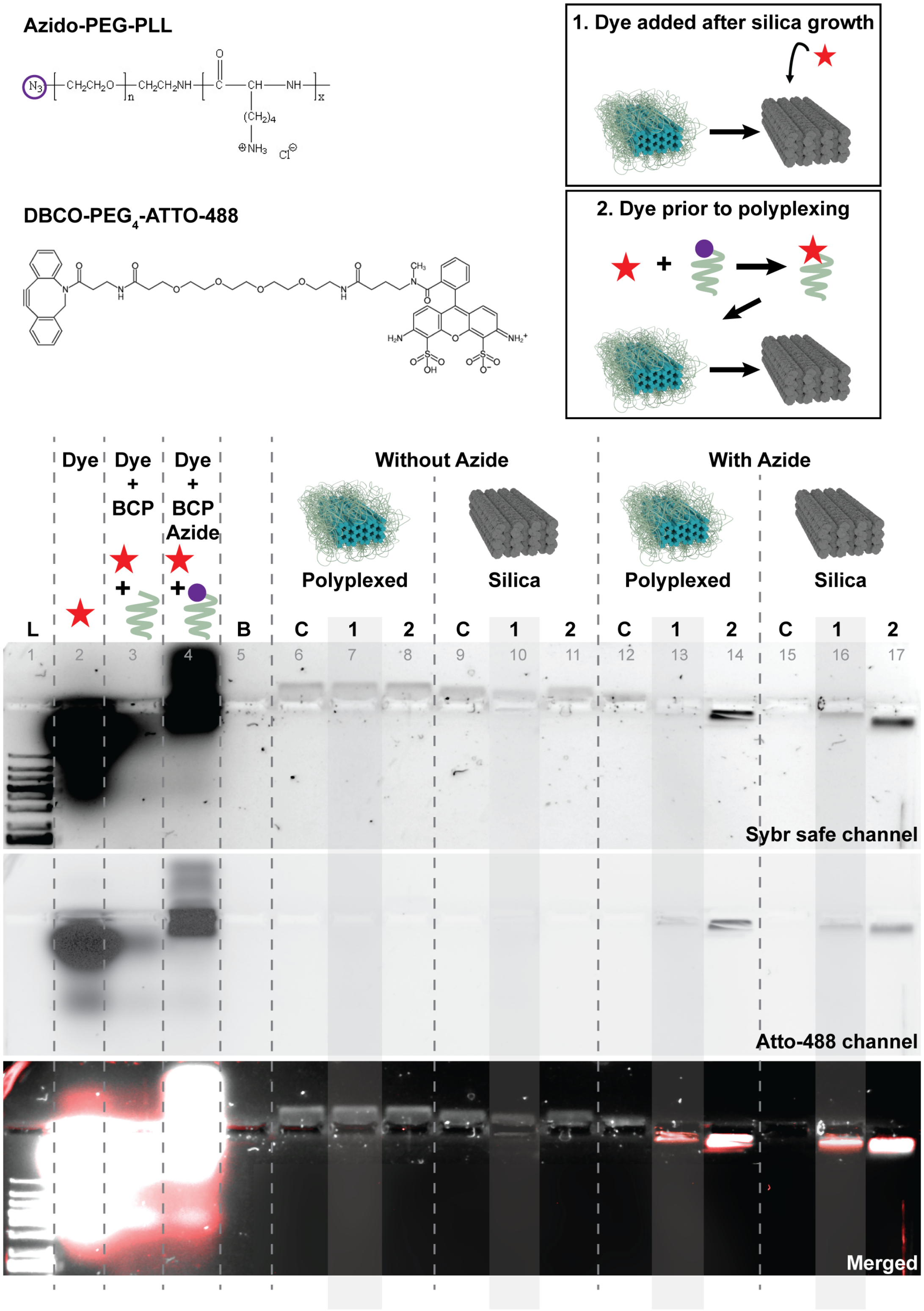

Figure S 31. AGE analysis to confirm the accessibility of functional moieties after silica growth. For this experiment block copolymers with azido group (PLL-PEG-N3) were used. As a control, block copolymers without the azido groups were used (PLL-PEG). For fluorescent tagging of the azido groups, DBCO modified Atto-488 dyes were added to the reaction mixture in two separate ways: 1) The dye was added to the block copolymer prior to the polyplex micellization step. This would serve as the dye control set. 2) The dye was added after silica growth. The two sets can be compared to confirm the accessibility. To facilitate the comparison, agarose gels were scanned for fluorescent signal from the dye using the blue channel, followed by post-staining with Sybr safe DNA stain to observe the fluorescence from the DNA bands. A merged version of the two fluorescent images can be observed in the bottom of the figure, with red fluorescent signal from the dye, while grey bands depict the fluorescence from the DNA stain.

Lanes 2, 3 and 4 are the negative controls without the origami. From lane 4 it can be confirmed that the DBCO-containing fluorescent dye reacts with azido tagged block copolymer. However, the aggregation along with the smear towards the negative terminal of the gel suggests a highly dispersed collection of polymer lengths. It should be noted that this effects the average-migration of the resulting DNA origami polyplex micelles.

Next, lanes 6 to 11 contain negative control structures prepared using block copolymers without the azido group. The absence of fluorescent bands in the Atto-488 channel suggests that there is not any non-specific interaction between the dye and DNA origami. Lanes 12 to 17 contain structures prepared using block copolymer with the azido group. From the lanes 14, 17 (Dye control lanes) and 13, 16, the presence of fluorescence in the Atto-488 channel can be confirmed, further confirming the accessibility of azido group after silica growth. We hypothesize that the difference in band intensities could arise from the loss of structures during the multiple purification steps as compared to the dye-control structures.

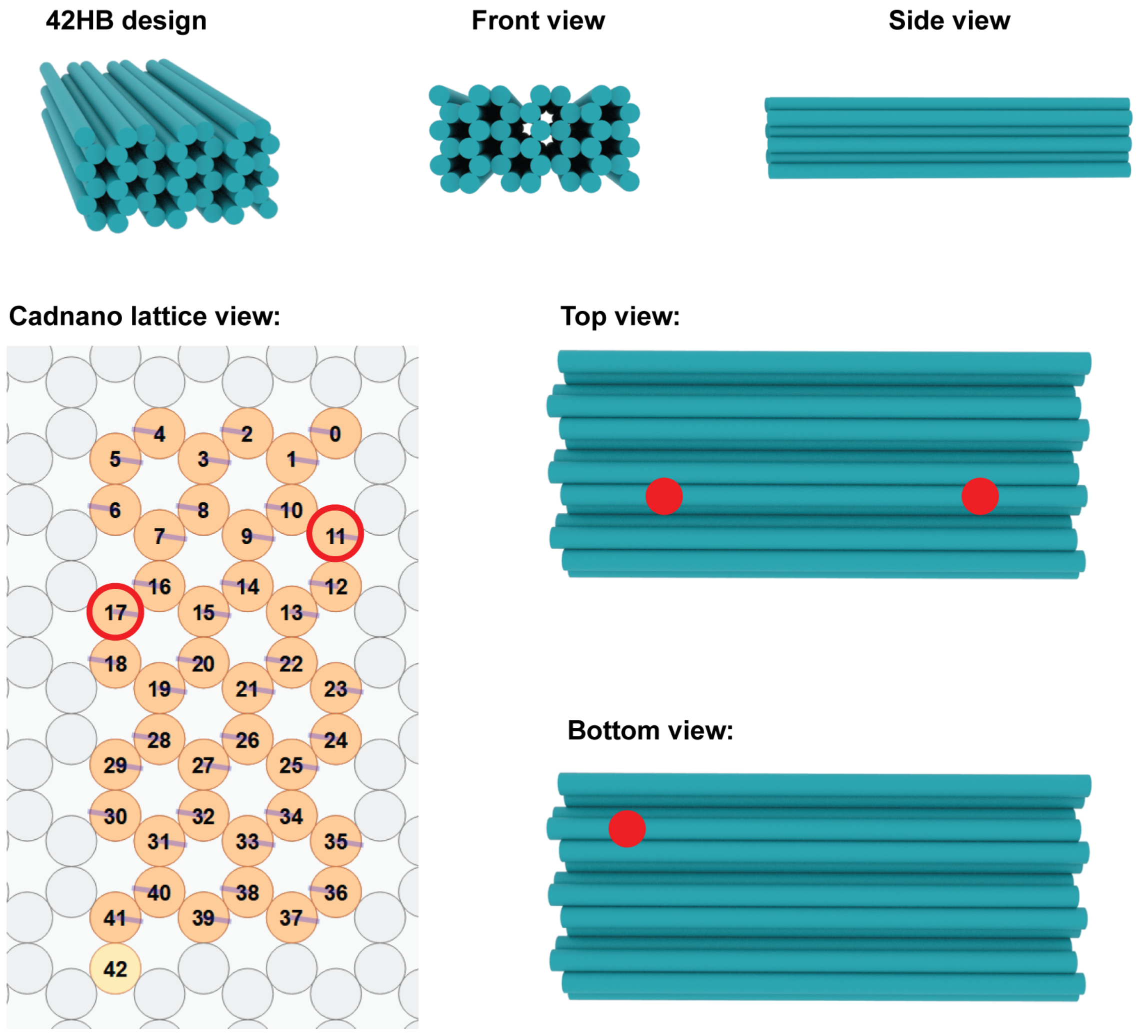

Figure S 32. Representative models for 42HB design. The top row graphics represent the perspective, top and the side view. The cadnano lattice view market with red circles represents the helices that consist of the binding sites. The top and bottom view marked with red dots represent the position of the 3 binding sites on the 42HB design that was used to attach IONPs.

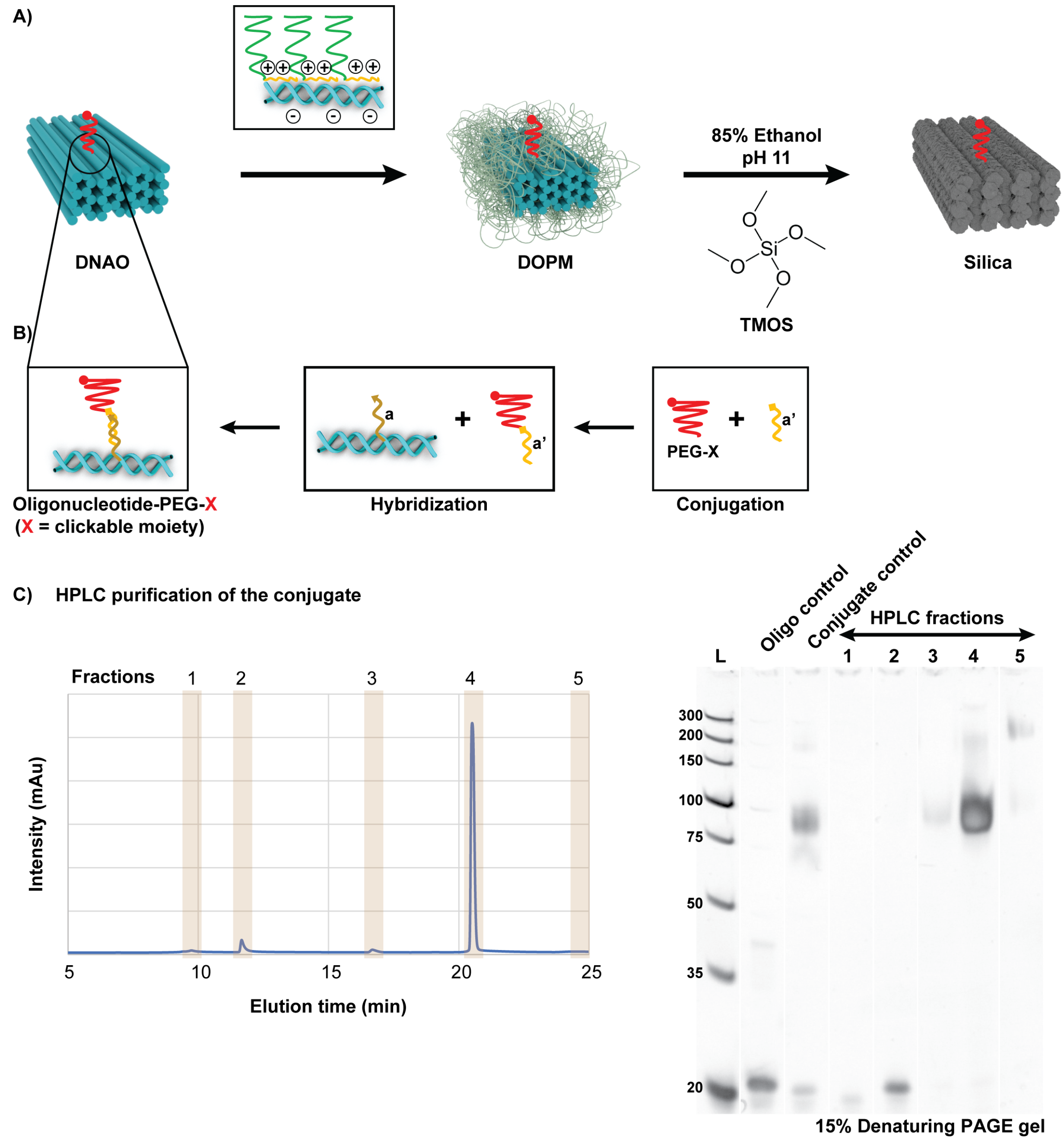

Figure S 33. Preparation of DBCO-PEG-oligonucleotide conjugate that serve as that binding site on the structures. A) Schematic representation of the strategy that enables maintaining the functionality during silica growth. B) Schematic representation of the preparation of the conjugate. C) HPLC purification trace and the corresponding PAGE gel analysis of the different fractions that were collected after the purification.

**1 Binding Site: Silica coated**

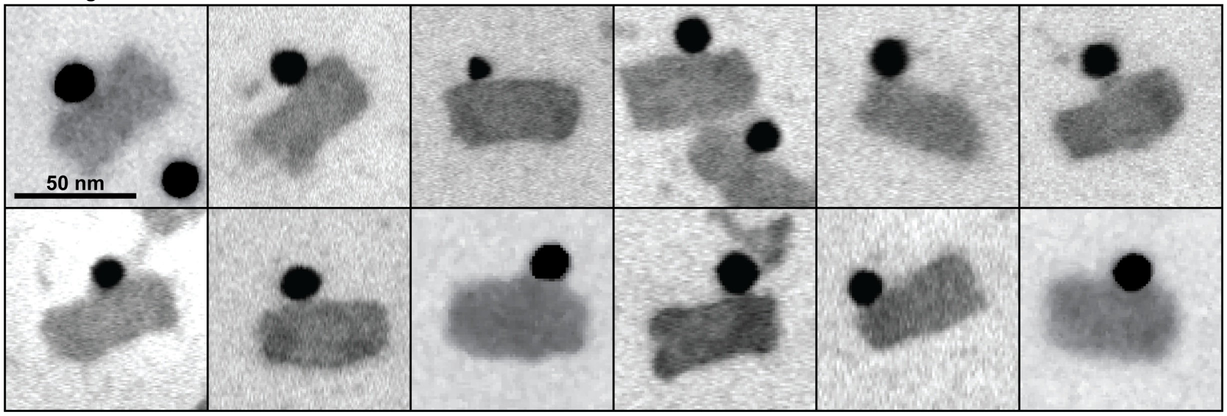

**1 Binding Site: Polyplexed (Stained with Uranyl Formate)**

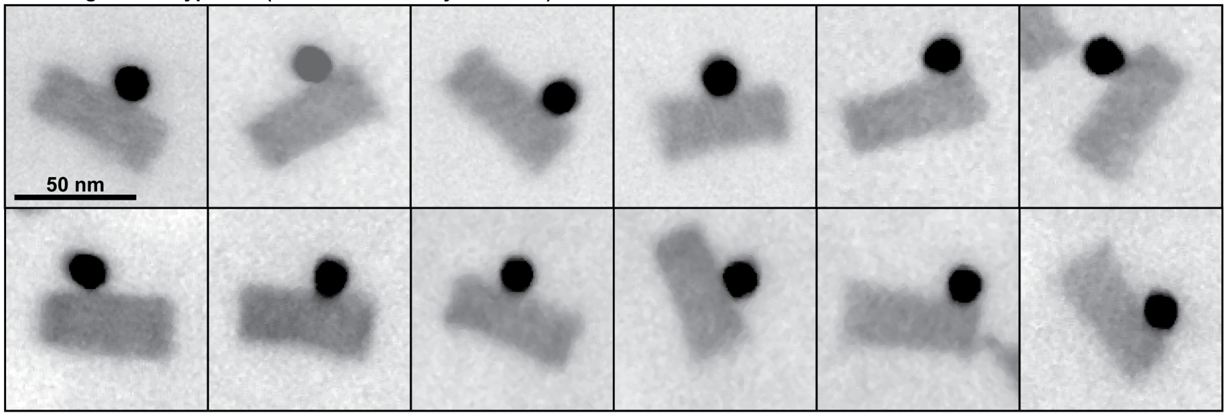

**1 Binding Site: Control (Stained with Uranyl formate)**

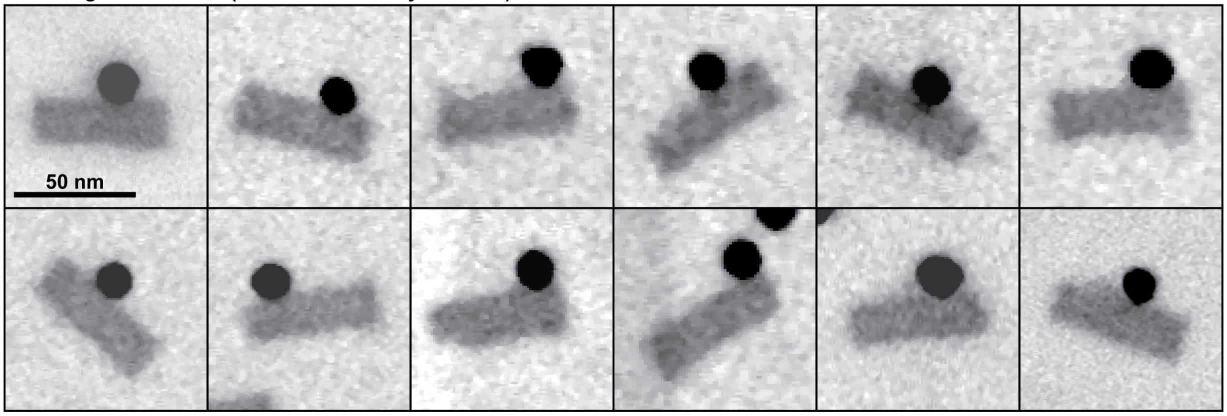

Figure S 34. Collection of electron micrographs for DNA brick structures carrying a single binding site (monofunctional DNA nanoparticle). Scale bars = 50 nm. For micellization, the K10P5k block co-polymers were used.

**2 Binding sites: Silica coated**

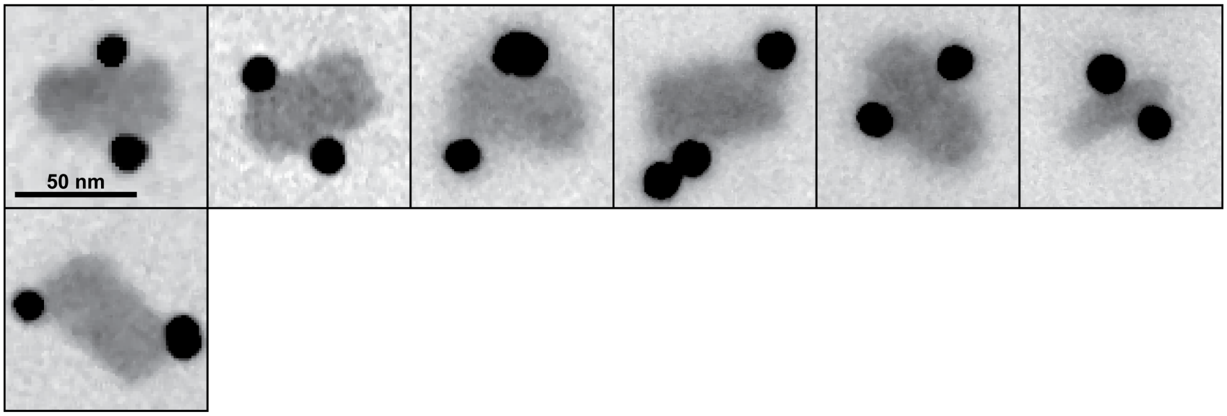

**2 Binding sites: Polyplexed (Stained with Uranyl Formate)**

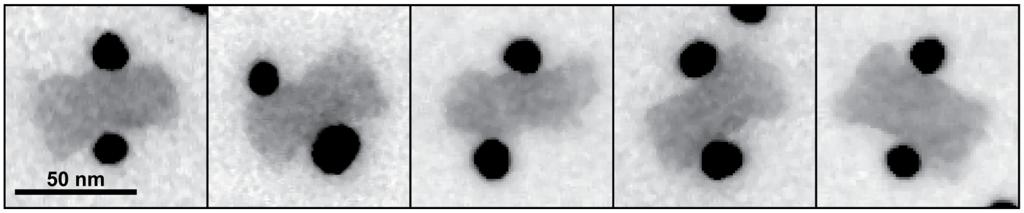

**2 Binding sites: Control (Stained with Uranyl Formate)**

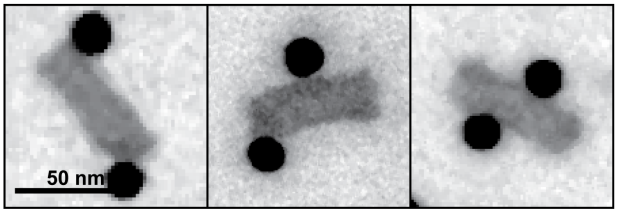

Figure S 35. Collection of electron micrographs for DNA brick structures carrying two binding sites. Scale bars = 50 nm. For micellization, the K10P5k block co-polymers were used.

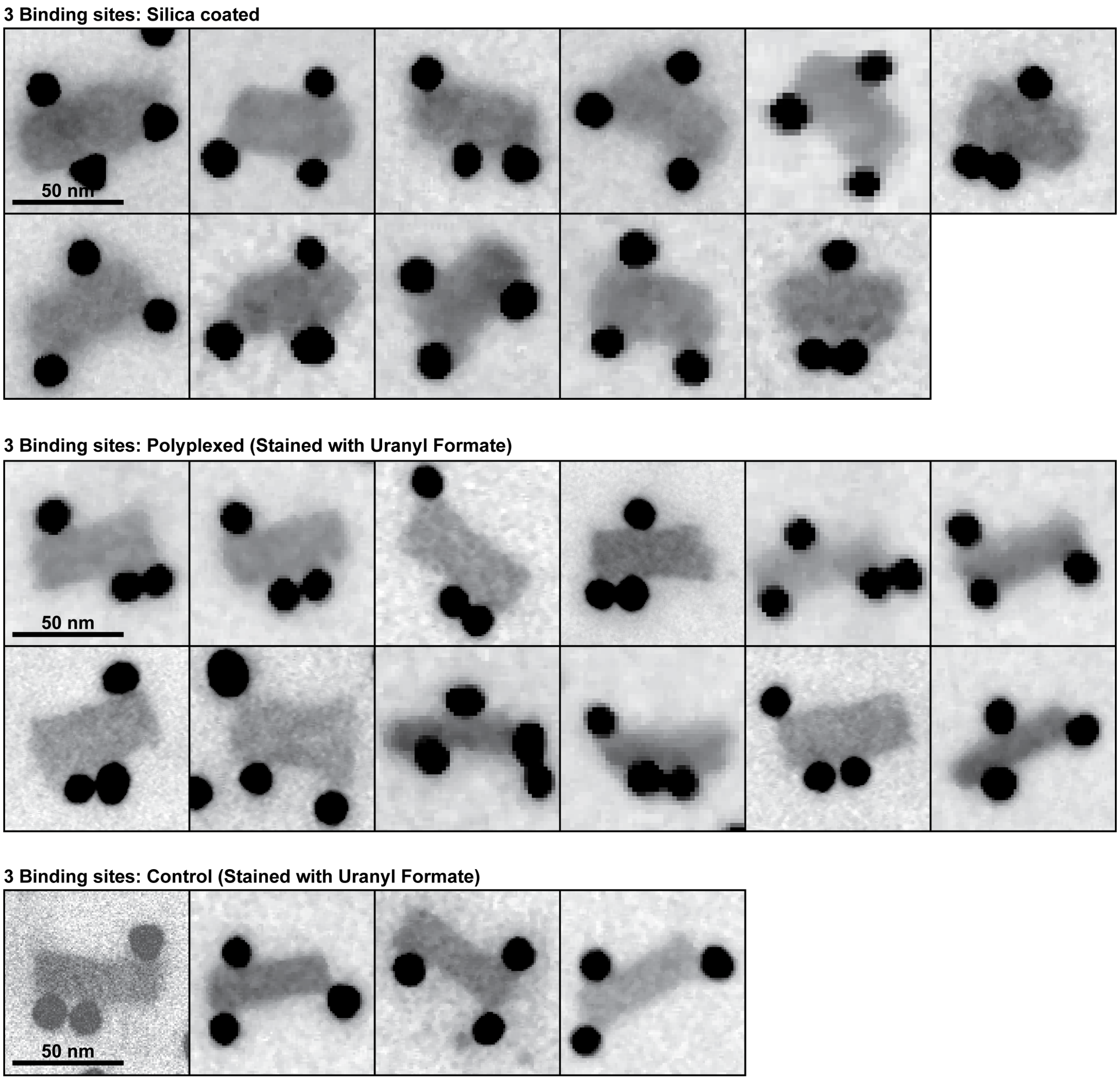

Figure S 36. Collection of electron micrographs for DNA brick structures carrying three binding sites. Scale bars = 50 nm. For micellization, the K10P5k block co-polymers were used.

### References.

(1) Engelhardt, F. A. S.; Praetorius, F.; Wachauf, C. H.; Brüggenthies, G.; Kohler, F.; Kick, B.; Kadletz, K. L.; Pham, P. N.; Behler, K. L.; Gerling, T.; Dietz, H. Custom-Size, Functional, and Durable DNA Origami with Design-Specific Scaffolds. *ACS Nano* **2019**, *13* (5), 5015–5027. https://doi.org/10.1021/acsnano.9b01025.

(2) Fragasso, A.; De Franceschi, N.; Stömmer, P.; van der Sluis, E. O.; Dietz, H.; Dekker, C. Reconstitution of Ultrawide DNA Origami Pores in Liposomes for Transmembrane Transport of Macromolecules. *ACS Nano* **2021**, acsnano.1c01669. https://doi.org/10.1021/acsnano.1c01669.

(3) Agarwal, N. P.; Matthies, M.; Gür, F. N.; Osada, K.; Schmidt, T. L. Block Copolymer Micellization as a Protection Strategy for DNA Origami. *Angew. Chem. Int. Ed.* **2017**, *56* (20), 5460–5464. https://doi.org/10.1002/anie.201608873.

(4) Anastassacos, F. M.; Zhao, Z.; Zeng, Y.; Shih, W. M. Glutaraldehyde Cross-Linking of Oligolysines Coating DNA Origami Greatly Reduces Susceptibility to Nuclease Degradation. *J. Am. Chem. Soc.* **2020**, *142* (7), 3311–3315. https://doi.org/10.1021/jacs.9b11698.
